# Nanopore-based sequence deconvolution of diverse glycosaminoglycans

**DOI:** 10.64898/2026.08.08.743666

**Authors:** Lemuel L. M. Szeto, Anna Yucknovsky, Daniel P. Cole, Hagan Bayley, Benjamin G. Davis, Yujia Qing

## Abstract

Glycosaminoglycan (GAG) polysaccharides play vital roles in animal physiology and disease^1^. Their diverse and intricate patterns of sulfation and epimerization endow them with an extensive potential to encode functional information^2,3^. GAG characterization, however, remains a formidable challenge for state-of-the-art ensemble-based techniques^4,5^. Single-molecule techniques are uniquely suited for analysing complex mixtures^6,7^. Here, we report the single-molecule resolution and counting of diverse GAG di- and oligosaccharides derived from longer heterogeneous chains as part of a deconvolutive nanopore-based workflow that requires no fractionation and is operationally simple. Modular chemical deacylation and amino-selective ring-contractive formation of electrophilic aldehydes enable the parsing of libraries of GAG structures into simplified sets of reactive anhydrosugars for nanopore readout via reversible covalent adduct formation. Discrete clustering of event amplitudes enables direct sugar sizing (∼10 % step change per residue), which can be coupled to precisely resolved amplitude differences that further reveal sugar fine structure—including the number and position of sulfate groups (∼2 % step change per sulfate) alongside single-atom stereochemistry (∼0.5 % step change between epimers). Guided by chemical logic, the reverse mapping of resolved anhydrosugars to their precursors covers ∼84–100 % of all disaccharides and their eliminative digestion variants in natural heparan sulfate (HS). We demonstrate the practical utility and scope of our approach through the compositional analysis of a panel of HS polysaccharides that together encompass natural GAG structural diversity. Moreover, we detect contaminants in heparin, including oversulfated chondroitin sulfate found in an authentic pharmaceutical heparin sample previously implicated in a global healthcare crisis. Together, our results suggest a general chemo-biophysical framework for the precise and sensitive characterization of GAGs that extends to other aminosugar biopolymers. When adapted for portable, widely used nanopore sequencing devices, our approach may offer a path towards the long-sought ‘democratization’ of glycan analysis.

## Introduction

Glycosaminoglycans (GAGs), a major class of linear aminosugar polysaccharides, are integral and ubiquitous components of the animal cell glycocalyx and extracellular matrix^1^. By interacting with hundreds of proteins^8^ in ways that can strongly depend on GAG fine structure^3,9,10^, GAGs modulate diverse physiological processes—including cell growth and differentiation^1,11–13^, development^14^, microbial pathogenesis^15,16^, inflammation^17,18^, coagulation^19^, and neurodegeneration^20^. GAGs of the heparan sulfate (HS) subclass appear to dominate such functional interactions^10^. HS chains are initially assembled as compositionally homogeneous precursors of repeating GlcA-GlcNAc disaccharide units (GlcA = glucuronic acid and GlcNAc = *N*-acetylglucosamine); these precursors are subsequently elaborated into heterogeneous structures containing up to 27 unique disaccharide units, each defined by a specific combination of *N*-substitution, sulfation, and uronic acid (UA) epimerization^2,3^ (Extended Data Fig. 1). Yielding ∼27*^n^* sequence permutations in a chain of *n* units, HS exhibits tremendous sequence complexity and is therefore considered one of the most ‘information-dense’ molecules in nature^2^.

Consequently, the characterization of HS and other GAGs for structure–function correlation remains a formidable challenge^4,5^. State-of-the-art analytical techniques such as tandem mass spectrometry (MS/MS)^21–23^, ion mobility MS^24–28^, and cryogenic infrared (IR) spectroscopy^29,30^ currently resolve only a limited subset of GAG di- and oligosaccharides (Supplementary Discussion 1). Because MS differentiates ions strictly by their mass-to-charge ratios, it is inherently blind to GAG regio- and stereoisomerism. Although solutions to this ‘isomer problem’ exist via various fragmentation and ion mobility augmentations to the basic MS configuration, their implementation can pose new challenges (e.g., ion activation causing GAG desulfation)^4,5^. Furthermore, MS and its variants typically operate at the ensemble level and therefore may lack the power needed to identify GAG structures within complex mixtures^7^; this conundrum is not entirely resolved by their coupling with fractionation procedures that can be lengthy and laborious^31,32^, thereby limiting application and throughput. Moreover, the size, sophistication, and cost of many state-of-the-art glycoanalytical instruments can present major barriers to their widespread adoption^33^.

Single-molecule techniques can pinpoint the individual components of challenging mixtures of compounds^6,7^. Protein nanopores, in particular, can sense and resolve a broad spectrum of analytes at the single-molecule level^34^; this capability, together with their ready incorporation into low-cost, portable devices such as the MinION^35^, positions them as potentially powerful tools for glycan characterization^7^. Although initial approaches using solid-state^36–40^ and protein^41–45^ nanopores have yielded signals for various GAGs (Supplementary Discussion 2), the often fleeting nature of these signals and their overlapping amplitudes have thus far precluded the resolution of individual sugar residues, much less their fine structural features. Crucially, these preliminary studies have illustrated the need for a fundamental shift in strategy to simultaneously overcome two key challenges pertaining to the kinetics of sensing and GAG structural diversity. First, event lifetimes— which have largely been limited to the sub-millisecond regime, yielding event ‘spikes’ with poor signal-to-noise ratios—would need to be drastically prolonged. Second, analytical capability would require significant expansion to resolve large libraries of unique GAG structures; this would necessarily encompass disaccharides, the fundamental GAG repeat unit, while ideally generalizing to longer oligosaccharides.

Here, we present a strategy of deconvolutive glycan ring contraction^46–49^ that overcomes both challenges through (i) the installation of electrophilic aldehydes, which enable a signal-rich mode of nanopore sensing via reversible covalent adduct formation, and (ii) the convergent transformation of libraries of GAG structures into simplified readout sets in a way that logically decodes precursor identity. Together, these allow the precise resolution and counting of diverse di- and oligosaccharides derived from longer chains via a fractionation-free and operationally simple workflow (Fig. 1).

**Fig. 1.**
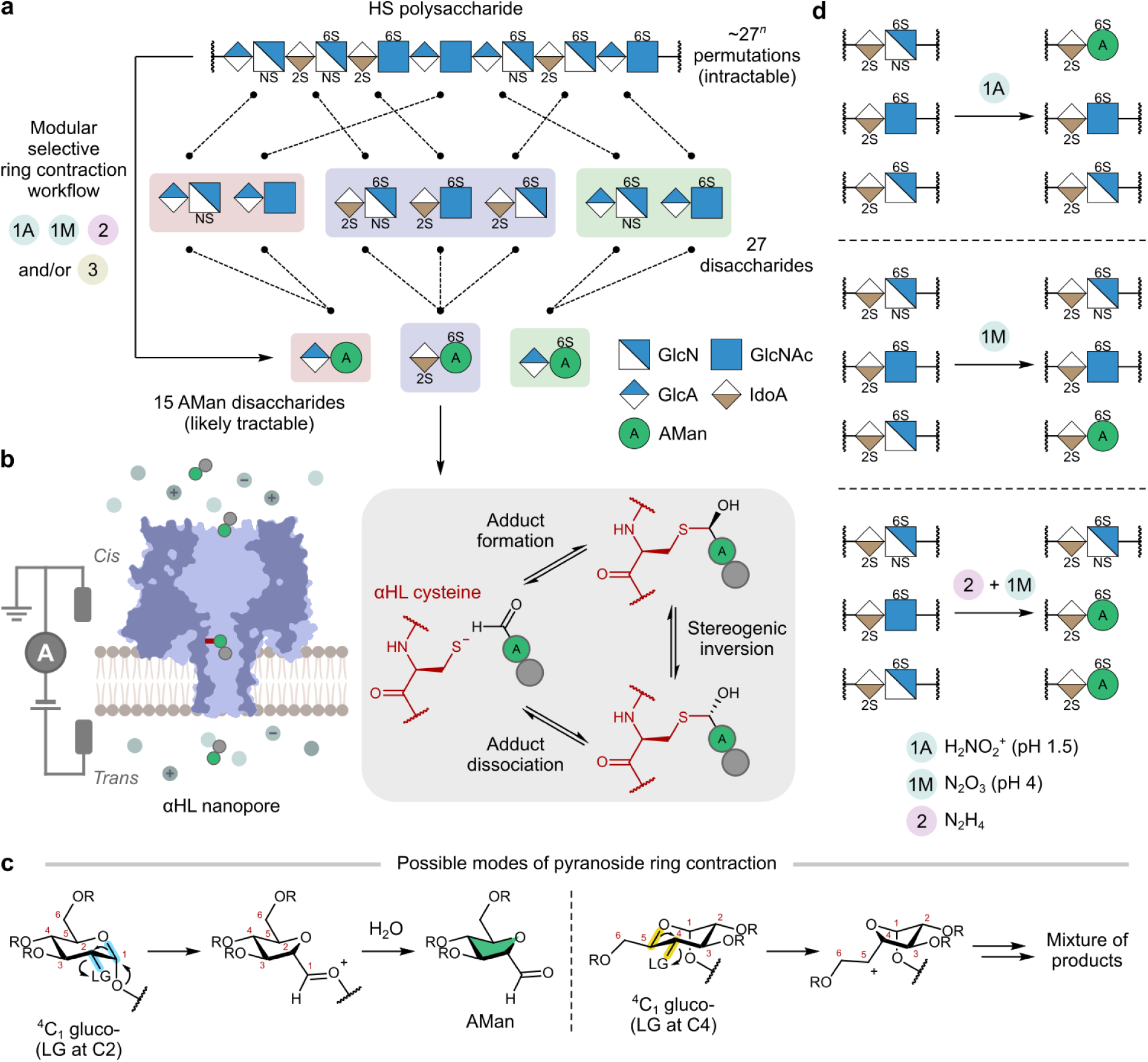
Glycan ring contraction enables sequence deconvolution and covalent sensing. **a,** A modular selective ring contraction workflow for the deconvolution of diverse GAG di- and oligosaccharides derived from longer chains into simplified sets of anhydrosugars, each bearing a reactive aldehyde. The dashed lines between the top and middle rows indicate the conceptual grouping of disaccharide units within the depicted HS polysaccharide by identity; the dashed lines between the middle and bottom rows indicate the chemical mapping of these grouped disaccharides to their corresponding anhydrosugars. The 27 unique disaccharides that emerge from HS biosynthesis map to 15 AMan disaccharides. *n*, number of disaccharide units. **b,** Anhydrosugar readout using αHL nanopores via reversible covalent adduct formation between anhydrosugar C1 aldehydes and a single cysteine thiolate. The resulting adducts undergo characteristic stereogenic inversion at the thiohemiacetal carbon. **c,** Possible modes of pyranoside ring contraction leveraging the strong ^4^C_1_ conformational preference of glucopyranosides, coupled with the presence of a suitable leaving group (LG) at C2, to generate AMan^49^. Ring carbon atoms are numbered, and the antiperiplanar O–C and C–LG bonds in each ring contraction mode are highlighted. R = H, SO_3_H, or any generic carbon substituent. **d,** The conditional generation of reactive anhydrosugars in a way that decodes precursor *N*-substituent identity.

## Results

### Glycan ring contraction designed for sequence deconvolution and covalent sensing

The elucidation of GAG fine structural features would ideally require the generation of prolonged sensing events with lifetimes spanning milliseconds to seconds. Covalent sensing, wherein nanopores and analytes form covalent adducts, offers a potentially powerful solution; however, it remains largely limited to functional groups absent in GAGs (e.g., disulfides and *cis* diols^50–56^). Among reactive functional groups endogenous to glycans, the aldehyde at the reducing-end anomeric centre (C1) has long been exploited as an electrophilic handle for selective chemical functionalization^57^. While a putative target for sensing via adduct formation with protein nucleophiles (e.g., cysteine thiolates^55^), this aldehyde is masked as a lactol, and its severely diminished reactivity typically permits only poor functionalization yields even under vigorous conditions^58^.

Instead, we reasoned that the generation of unmasked aldehydes in glycans, preferably under mild and efficient conditions, would overcome this reactivity barrier and allow covalent sensing. The endocyclic ring oxygen of pyranosides can, under certain circumstances, participate in stereospecific migratory ring contraction via intramolecular nucleophilic substitution of a leaving group at C2 or C4 and concomitant cleavage of the O–C1 or O–C5 bond, respectively^46,59^ (Fig. 1c). Diazonium groups serve as excellent leaving groups in such migratory rearrangements and can be readily generated from amines via classical Tiffeneau–Demjanov *N-*nitrosation^60^. Applied to the C2 nitrogen in *N*-sulfated and *N*- unsubstituted glucosamine (GlcNS and GlcN) residues in HS, this would facilitate a ring contraction to 2,5-anhydromannose (AMan), within which the newfound C1 aldehyde is exocyclic and unmasked^47,61^. Furthermore, *N*-sulfate (NS) and free amine (N) substituents can be rapidly converted to diazonium groups with high selectivity (∼87–98 %) using distinct reagents^61^; specifically, the former reacts with H_2_NO_2_ ^+^ at pH 1.5 (**<u>a</u>**cidic condition **1A**; Fig. 1d), whereas the latter reacts with N_2_O_3_ at pH 4 (**<u>m</u>**ildly acidic condition **1M**; Fig. 1d; see Extended Data Fig. 2 for mechanism; *N*-acetyl (NAc) groups are inert).

Intriguingly, this suggested a utility of the Tiffeneau–Demjanov ring contraction not only in generating powerful aldehyde electrophiles for covalent sensing but also, by virtue of its pH- addressable chemoselectivity, in establishing a deconvolutive strategy in which libraries of GAG di- and oligosaccharides are converted into simplified sets of anhydrosugars in a way that would logically decode *N*-substituent identity (NS vs N vs NAc). Crucially, this structural simplification would inherently minimize potential overlap in nanopore signal amplitudes. Because the C2 amide (NAc) in the GlcNAc residues shared by HS and other GAGs is inert to both conditions **1A** and **1M**, we further envisaged the use of chemoselective deacylating hydrazinolysis^62^ to convert the amide to a free amine (condition **2**; Fig. 1d), thereby vitally broadening the scope of deconvolution to encompass all natural GAG *N*-substituent variants (NS, N, and NAc). By design, all three reactions (**1A**, **1M**, and **2**) preserve all other GAG structural features, including sulfate groups, glycosidic linkages, and UA stereochemistry^61,62^. In this way, their modular combination (e.g., **1A**, **1M**, or **2 + 1M**; Fig. 1d) would enable the deconvolution of 17 out of the 27 unique disaccharide units that emerge from HS biosynthesis—which collectively account for ∼84–100 % by molar abundance of all units in natural HS (Table S4)—into a mere seven AMan disaccharides (Extended Data Fig. 3a).

We engineered two heteroheptameric α-haemolysin (αHL) pores—αHL-Cys115 and αHL- Cys117—with lumen volumes anticipated to be suitable for the capture of diverse sugars as nucleophilic ‘nanoreactors’ for covalent sensing. Both pore variants were assembled in an A_6_B_1_ subunit stoichiometry (Methods). To eliminate potential side reactions such as metal chelation and imine formation^55^, the six A subunits and the single B subunit in each variant were mutated to collectively produce lysine- and methionine-free lumens; the B subunit was additionally equipped with a single cysteine residue at either position 115 or 117 to provide a thiolate nucleophile.

### Discrete clustering of event amplitudes enables direct sugar sizing

To test this design, we subjected a sample of polysaccharidic porcine intestinal heparin to ring contraction at pH 1.5 (condition **1A**) and subsequent covalent sensing without further fractionation (Extended Data Fig. 4a). Ring contraction in these and other polysaccharides is necessarily concomitant with glycosidic bond cleavage (Extended Data Fig. 2), yielding AMan fragments each bearing the desired, unmasked C1 aldehyde. Gratifyingly, upon introducing these heparin-derived fragments to the αHL-Cys117 pore from the *cis* side under an applied voltage of +150 mV, numerous rectangular events with millisecond-to-second lifetimes immediately emerged (Extended Data Fig. 4b and Methods); these were consistent with the reversible nucleophilic addition of the single αHL cysteine thiolate to unmasked AMan aldehydes to form thiohemiacetal adducts (Fig. 1b). Control experiments using non-ring-contracted heparin failed to produce any such covalent events, consistent with the expectedly poor reactivity of aldehydes sequestered in lactol form (Fig. S2).

An analysis of the residual current values of these heparin-derived events—expressed as *I*_res_ = 100 × (*I*_e_/*I*_o_), where *I*_e_ and *I*_o_ respectively denote the event and open-pore currents (Methods)—and their subsequent comparison with structure-defined AMan standards (Supplementary Methods) revealed a striking correlation with sugar chain length. When benchmarked individually, mono-, di-, and tetrasaccharides yielded highly distinct *I*_res_ values of ∼96 %, ∼87 %, and 64–67 %, respectively (*N* ≥ 3 pores for each length type), with each additional sugar residue contributing a ∼10 % step decrease (Fig. 2a and Table S5). Furthermore, the corresponding covalent events exhibited lifetimes of 140 ± 40 ms, 270 ± 40 ms, and 200 ± 70 ms (mean ± s.d.), respectively, thus providing an ample window for accurate *I*_res_ extraction. The lack of events observed with disaccharides bearing free anomeric centres, even when used at high concentrations, again confirmed the insufficient reactivity of aldehydes masked as lactols (Fig. S3). Identical experiments with ring-contracted heparin and AMan standards using the alternative αHL-Cys115 pore revealed clustering around similar *I*_res_ values (Extended Data Figs 4c, 5a, and Table S6).

**Fig. 2.**
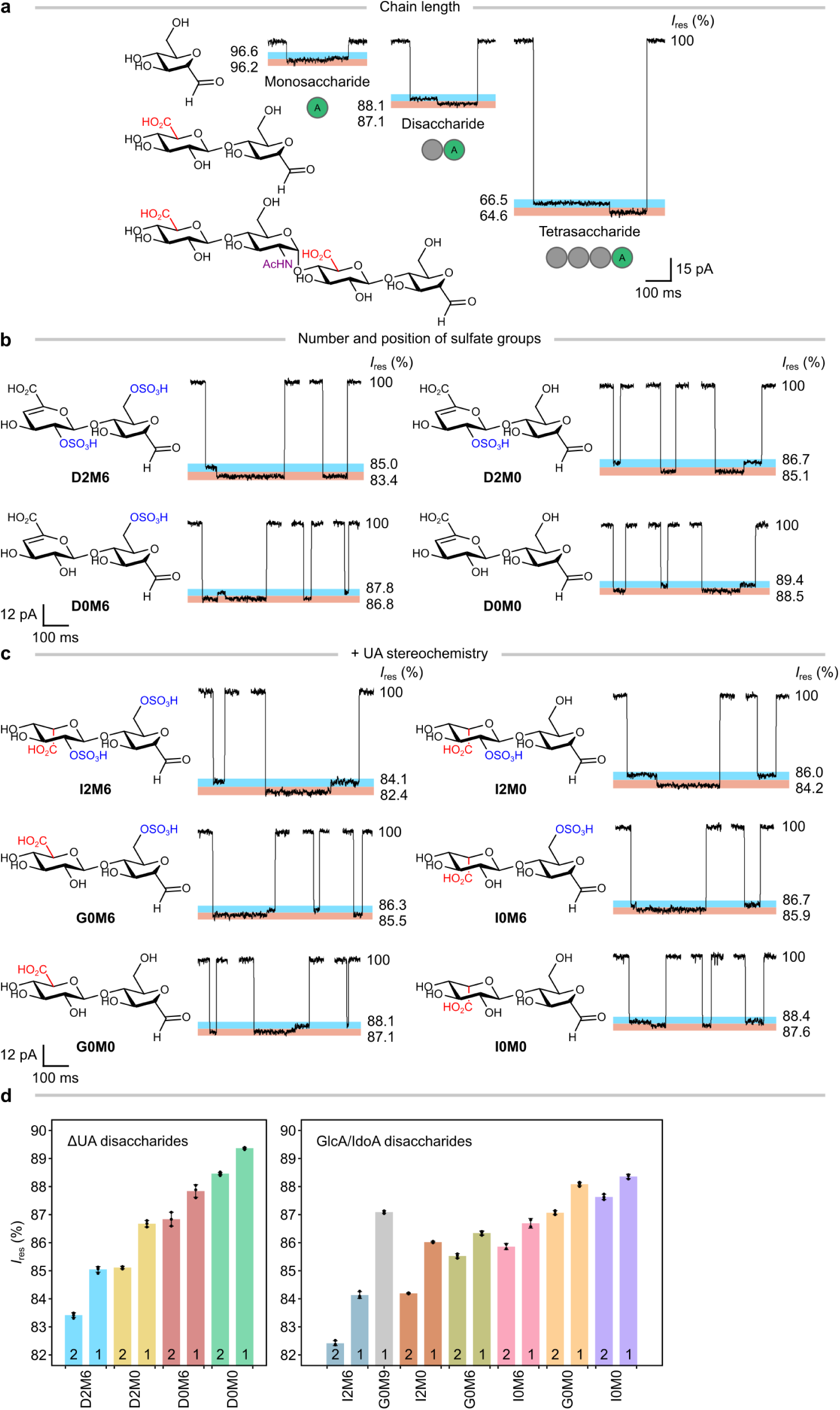
Resolution of GAG chain length and fine structural features using the αHL-Cys117 pore. **a,** Mono-, di-, and tetrasaccharides (M0, G0M0, and G0A0G0M0, respectively) yielded distinct *I*_res_ values, with each additional sugar residue contributing a ∼10 % step decrease. AMan and non-AMan residues are depicted as green and grey circles, respectively. **b–c,** ΔUA disaccharides with various sulfation states (**b**) and UA disaccharides with various sulfation and epimerization states (**c**) were resolved based on differences in the *I*_res_ values of interconverting levels, highlighted in blue and pink (e.g., ∼2 % step decrease per sulfate and ∼0.5 % step decrease from IdoA to GlcA). **d,** The bar plots show the mean *I*_res_ levels of 11 AMan disaccharides recorded across *N* ≥ 3 pores, except for I0M6 (*N* = 2) due to limited material (also see Table S5). Points indicate *I*_res_ levels from independent pores; vertical error bars indicate s.d. The bars are numbered either ‘1’ or ‘2’; for each sugar with paired interconverting levels, ‘1’ indicates the level with the higher *I*_res_ value. **Recording conditions:** 4 M LiCl, 20 mM HEPBS, 40 μM EDTA, titrated to pH 8 using KOH; sugars (*cis*); +150 mV (*trans*); 23.8 ± 1 °C. Sugar concentrations ranged from 0.19 mM to 0.84 mM, corresponding to 42–139 µg of material. All *I*_res_ values are reported to one decimal place.

We then tested whether sugars within complex mixtures could be quantified according to chain length. Probed by the αHL-Cys115 pore, the heparin-derived fragments yielded discrete clusters that were immediately discernible by pronounced differences in *I*_res_ (∼10 % or more; Extended Data Fig. 4c). Having established the signatures of various length types through the use of standards, we confirmed that these clusters arose from mono-, di-, and oligosaccharides, comprising 10.1 ± 1.4 %, 70.1 ± 2.9 %, and 19.8 ± 2.2 % of all events, respectively (mean ± s.d. from *N* = 3 pores; Table S7). Moreover, within the oligosaccharide cluster, events were separable into a major fraction centred within the tetrasaccharide range (*I*_res_ 60–70 %), accounting for 14.4 ± 1.4 % of total sugar events, and a tailing fraction presumed to correspond to higher-order saccharides (*I*_res_ < 60 %; Extended Data Fig. 4d). These resolved event proportions were in good agreement with relative molar abundances determined using bulk fractionation methods^63^. Notably, previous glycan sizing approaches have been limited to compositionally homogeneous GAG chains of > 54 kDa, producing only broad, overlapping size distributions^36,40^; by comparison, our approach enabled direct sizing down to individual residue lengths alongside quantification within complex mixtures via real-time event counting.

### Precisely resolved interconverting events reveal sugar substructural features

Within the event clusters delineating chain length, we observed smaller *I*_res_ subclusters that hinted at a sensitivity towards more subtle structural features, such as sulfation and epimerization—a capability we termed ‘subsizing’ (Extended Data Fig. 4). Moreover, many individual events displayed a pair of *I*_res_ levels demonstrating spontaneous, discrete level-to-level transitions in real time, or ‘interconversions’. To investigate both phenomena, we prepared and analysed a panel of 11 AMan disaccharide standards that together spanned diverse combinations of sulfation, epimerization, and UA saturation states (Extended Data Fig. 3c and Supplementary Methods; GAG moieties are referred to by their disaccharide structure code^64^).

Consistent with our results on sizing, disaccharides probed by the αHL-Cys117 pore yielded *I*_res_ levels within the 82.4–89.4 % range (Figs 2b–d and Extended Data Fig. 6a). Interconverting paired *I*_res_ levels were observed for all disaccharides but one (G0M9); such interconversions were consistent with stereogenic inversion at the thiohemiacetal carbon^65,66^, a feature previously reported with simple aliphatic and aromatic aldehydes^55,67^. Importantly, the additional information encoded in a second, temporally coupled *I*_res_ level—with both arising within a single molecular capture event and together yielding a characteristic *I*_res_ ‘doublet’—enabled sugars that shared an individual level to be discriminated, thereby expanding analytical capability. The *I*_res_ data for each level acquired from any single pore were generally well modelled by Gaussian distributions with low half-width at half-maximum (HWHM) values of ∼0.1 %, indicating excellent precision; furthermore, a comparison of these distributions across independent pores revealed high reproducibility, with centroid s.d. values of ∼0.1 % or less (Fig. S5 and Table S5). Together, the negligible probability of coincidental doublet overlap and the high reproducibility and precision of our measurements enabled structure resolution, even within complex mixtures (see below).

Clear trends relating to GAG substructural features were observed. First, more heavily sulfated disaccharides produced, on average, lower *I*_res_ values than their less sulfated counterparts, consistent with increased volume imparted by sulfation; for instance, the disulfated D2M6 yielded lower values than the monosulfated D2M0, whose values in turn fell below those of the non-sulfated D0M0 (Fig. 2b). Thus, *I*_res_ values were strongly correlated with the level of sulfation (∼2 % step decrease per sulfate), providing a powerful framework for its direct estimation (see below) alongside the resolution of sugar chain length.

Second, our means of subsizing extended to sulfation regioisomers: 2-*O*-sulfated disaccharides exhibited consistently lower *I*_res_ values (by 1–1.5 %) than their 6-*O*-sulfated counterparts—as seen, for instance, when comparing D2M0 against D0M6 (Fig. 2b). Accordingly, a sulfate group at the secondary C2 position is sensed by the pore as ‘larger’ than one at the primary C6 position.

Third, and remarkably, even stereoisomeric disaccharides, as well as those differing solely in their UA saturation states (Δ^4,5^-unsaturated UA (ΔUA) vs saturated UA, corresponding to a mass difference of 18 Da arising from elimination), yielded distinct *I*_res_ values. GlcA disaccharides produced consistently lower *I*_res_ values (by ∼0.5 %) than their iduronic acid (IdoA) C5 epimers, whose values in turn fell below those of their ΔUA counterparts (by ∼1 %; e.g., see G0M0, I0M0, and D0M0 in Figs 2b–c). Together, our results suggest a nanopore- sensed volume order of GlcA > IdoA > ΔUA. Notably, even single-atom stereochemistry, the most subtle of GAG structural features, was readily resolved by our approach.

These structurally correlated *I*_res_ differences, combined with the reproducibility and precision of our measurements and the enhanced resolving power afforded by level-to-level interconversions, enabled the unambiguous discrimination of all pairs among ten disaccharides (Figs 2b–c) using the αHL-Cys117 pore (Supplementary Note 1). Similar *I*_res_ values and trends were observed using the αHL-Cys115 pore (Extended Data Figs 5b–d, 6a, Fig. S6, and Table S6). Moreover, the two pores proved complementary: on rare occasions when a specific disaccharide pair was challenging to differentiate using one pore (such as G0M0 vs G0M9 using αHL-Cys117), it was readily resolved by the other (Extended Data Fig. 7). Ultimately, the αHL-Cys115 pore alone sufficed to identify any given ΔUA disaccharide from the four examined, whereas both pores in combination sufficed to identify any given disaccharide across the entire 11-member panel (Supplementary Note 1).

Moreover, substructural features such as sulfation were clearly discernible in longer oligosaccharides. In both pores, the di-6-*O*-sulfated AMan tetrasaccharide G0A6G0M6 yielded lower interconverting *I*_res_ levels (by ∼2 %) than the non-sulfated tetrasaccharide G0A0G0M0 (Fig. 2a and Extended Data Figs 5a, 6a). This ∼1 % step decrease in *I*_res_ per 6- *O*-sulfate group—consistent with our results in disaccharides—suggests that the *I*_res_ shifts imparted by individual sulfate groups remain conserved in these longer chains, thereby supporting sulfate counting beyond the disaccharide level.

Finally, a comparative analysis of unique GAG structures before and after ring-contractive deconvolution revealed that deploying our strategy at the disaccharide level might provide the greatest reduction in library size, and therefore maximum structural coverage for a given set of anhydrosugars (Extended Data Fig. 3). Indeed, our resolved panel of 11 disaccharides covers ∼84–100 % by molar abundance of all disaccharides, including their eliminative digestion variants, in natural HS (Table S4).

### A modular deconvolutive workflow enables compositional analysis of diverse polysaccharides

Consequently, we integrated hydrazinolysis (condition **2**), chemoselective ring contraction variants (conditions **1A** and **1M**), and eliminative digestion (condition **3**) into a modular deconvolutive workflow to resolve and count the di- and oligosaccharide constituents of GAG polysaccharides. This workflow was applied to a panel of six polysaccharide samples designed to encompass natural GAG structural diversity: (i) unfractionated porcine intestinal heparin (pUFH); (ii) 2-*O*-desulfated heparin (2DSH); (iii) 6-*O*-desulfated heparin (6DSH); (iv) heparosan (Hp), the unmodified biosynthetic precursor of HS; (v) fully *N*-sulfated heparosan (NSHp)^68^; and (vi) epimerized fully *N*-sulfated heparosan (ENSHp)^69^.

To begin, a subset of the panel comprising the three heparins (pUFH, 2DSH, and 6DSH) was subjected to eliminative digestion (condition **3**) followed by ring contraction at pH 1.5 (condition **1A**); the unpurified product mixtures, retaining all added buffer salts, enzymes, and cryoprotectant non-GAG sugars (Fig. S7), were loaded directly into our nanopore setup and probed using the αHL-Cys115 pore (Fig. 3a). Clear sensing events contributed by the heparins were observed and assigned based on direct matching with our resolved standards (Supplementary Methods and Table S8). No background events occurred in the absence of the heparins, demonstrating exceptional tolerance to salts and typical non-target biomolecules, and obviating the need for any prior fractionation (Fig. S8).

**Fig. 3.**
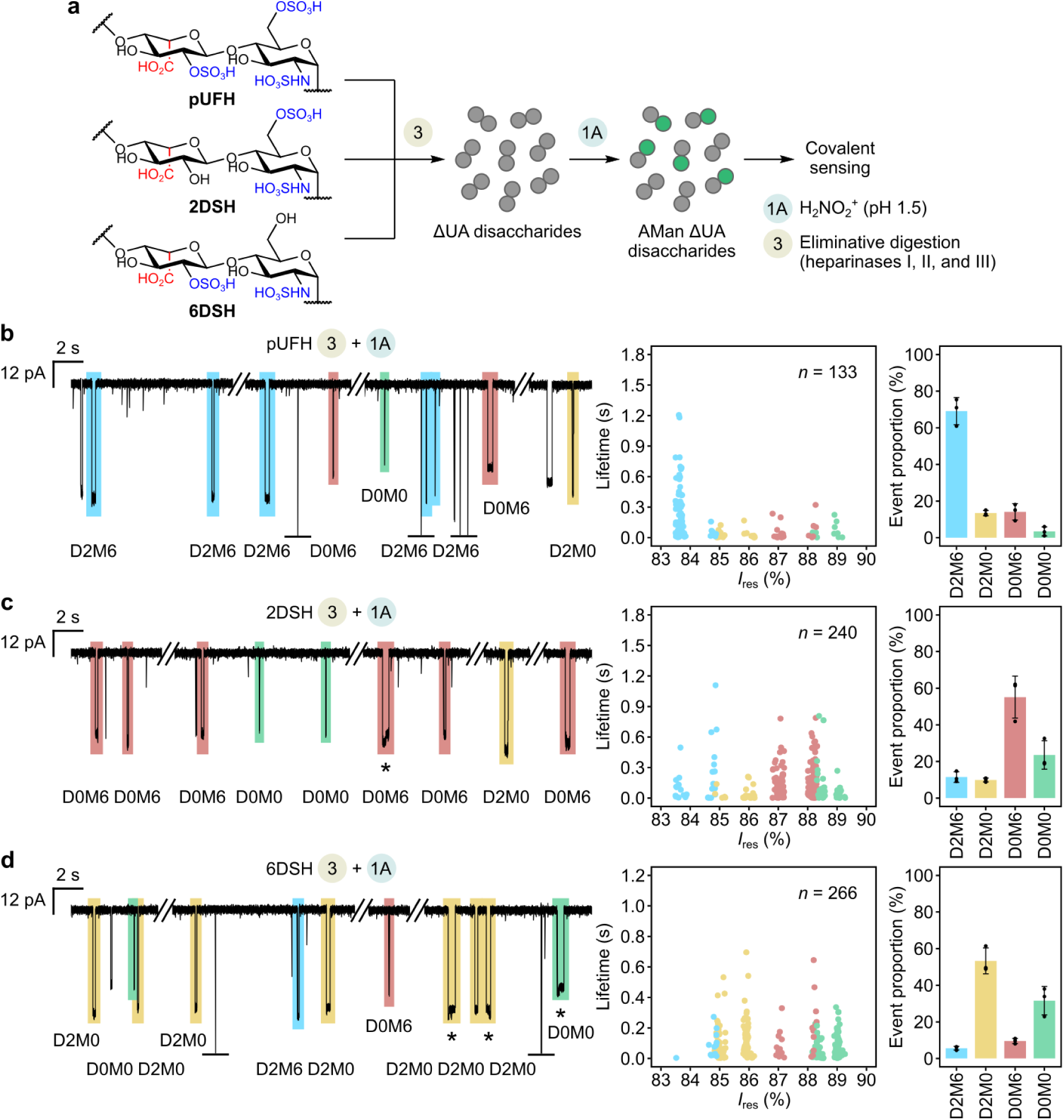
Deconvolutive compositional analysis of heparin polysaccharides. **a,** pUFH, 2DSH, and 6DSH (representative structures shown) were subjected to eliminative digestion using heparinases (condition **3**), followed by ring contraction at pH 1.5 (condition **1A**). The resulting product mixtures (∼0.3 mg) were directly introduced to the αHL-Cys115 pore. AMan and non-AMan residues are depicted as green and grey circles, respectively. **b–d,** The nanopore fingerprints of pUFH (**b**), 2DSH (**c**), and 6DSH (**d**) revealed disaccharides in various proportions, thereby enabling compositional analysis. Representative current traces are shown, along with assignments for the four ΔUA disaccharides D2M6, D2M0, D0M6, and D0M0 (Supplementary Methods). Interconversion events are marked with an asterisk. Graphically truncated events are marked with a horizontal bar. The scatter plots show assigned ΔUA disaccharide event levels from single pores (*n* = number of events), and levels are colour-coded as in the traces. The bar plots show mean ΔUA disaccharide event proportions across *N* = 3 pores; points indicate values from independent pores, and vertical error bars indicate s.d. **Recording conditions:** 4 M LiCl, 20 mM HEPBS, 40 μM EDTA, titrated to pH 8 using KOH; sugars (*cis*); +150 mV (*trans*); 23.8 ± 1 °C. Events with lifetimes under 2 ms were discarded (Methods).

Mean event proportions in what we termed the pUFH ‘fingerprint’ directly indicated the composition of pUFH, falling in the order D2M6 > D0M6 > D2M0 > D0M0; here, D2M6 emerged as the major component, accounting for 69.1 ± 7.4 % of all ΔUA disaccharide events (mean ± s.d. from *N* = 3 pores, 57–266 assigned events per pore; Fig. 3b). This result was consistent with the disaccharide compositions of various porcine intestinal heparins determined by high-performance liquid chromatography (HPLC)^70^ (Fig. S9). By comparison, mean event proportions in the 2DSH and 6DSH fingerprints displayed the respective orders D0M6 > D0M0 > D2M6 > D2M0 and D2M0 > D0M0 > D0M6 > D2M6— with D0M6 comprising 55.1 ± 11.5 % of the ΔUA pool in the 2DSH fingerprint (Fig. 3c) and D2M0 comprising 53.3 ± 7.1 % of the pool in the 6DSH fingerprint (Fig. 3d).

Across all heparins, independent experiments replicating the entire **3 + 1A** reaction sequence yielded reproducible fingerprints (Figs S10a–c), thereby indicating the robustness of our workflow. Typically, a sampling depth of only ∼50 assigned ΔUA disaccharide events, accumulated in under ∼30 min of recording, sufficed to establish stable event proportions. Such was the resolving power of our approach that we observed even unanticipated saturated UA disaccharides, which were unambiguously identified via paired interconversion events (Extended Data Figs 8a–c); these disaccharides were attributed to the ring- contractive cleavage of recalcitrant components within a digestion process otherwise deemed exhaustive by high-performance thin-layer chromatography (Fig. S10d). In this way, our deconvolutive workflow, when applied to complex polysaccharides, enabled ready and detailed compositional analysis, with a sensitivity for low-abundance components beyond those of bulk methods.

We next evaluated the three heparosans (Hp, NSHp, and ENSHp) under an alternative set of deconvolutive modules (conditions **1A**, **1M**, and **2**); these simultaneously cleave GAG chains and generate reactive anhydrosugars for covalent sensing, while preserving UA stereochemistry. Using appropriate module combinations to decode precursor *N*-substituent identity, Hp was subjected to conditions **2 + 1M**, whereas NSHp and ENSHp were directly reacted under condition **1A** (Fig. 4a). Upon introducing the unpurified product mixtures to the αHL-Cys117 pore, events were observed and again assigned based on direct matching with standards (Supplementary Methods and Table S9). While the Hp and NSHp fingerprints yielded almost exclusively G0M0 (Figs 4b–c), consistent with their apparent homogeneous compositions (Figs S12–13), the ENSHp fingerprint was dominated by two major components: G0M0 and I0M0 accounted for 59.7 ± 3.7 % and 40.3 ± 3.7 % of these events, respectively (mean ± s.d. from *N* = 3 pores, 83–145 assigned events per pore; Fig. 4d). By identifying IdoA disaccharides, the I0M0 event proportion provided a direct readout of the degree of epimerization in ENSHp. Notably, this single-molecule readout was in excellent agreement with the bulk metric determined using ^1^H nuclear magnetic resonance (NMR) spectroscopy (40.3 ± 3.7 % vs 38 %; Fig. S14).

**Fig. 4.**
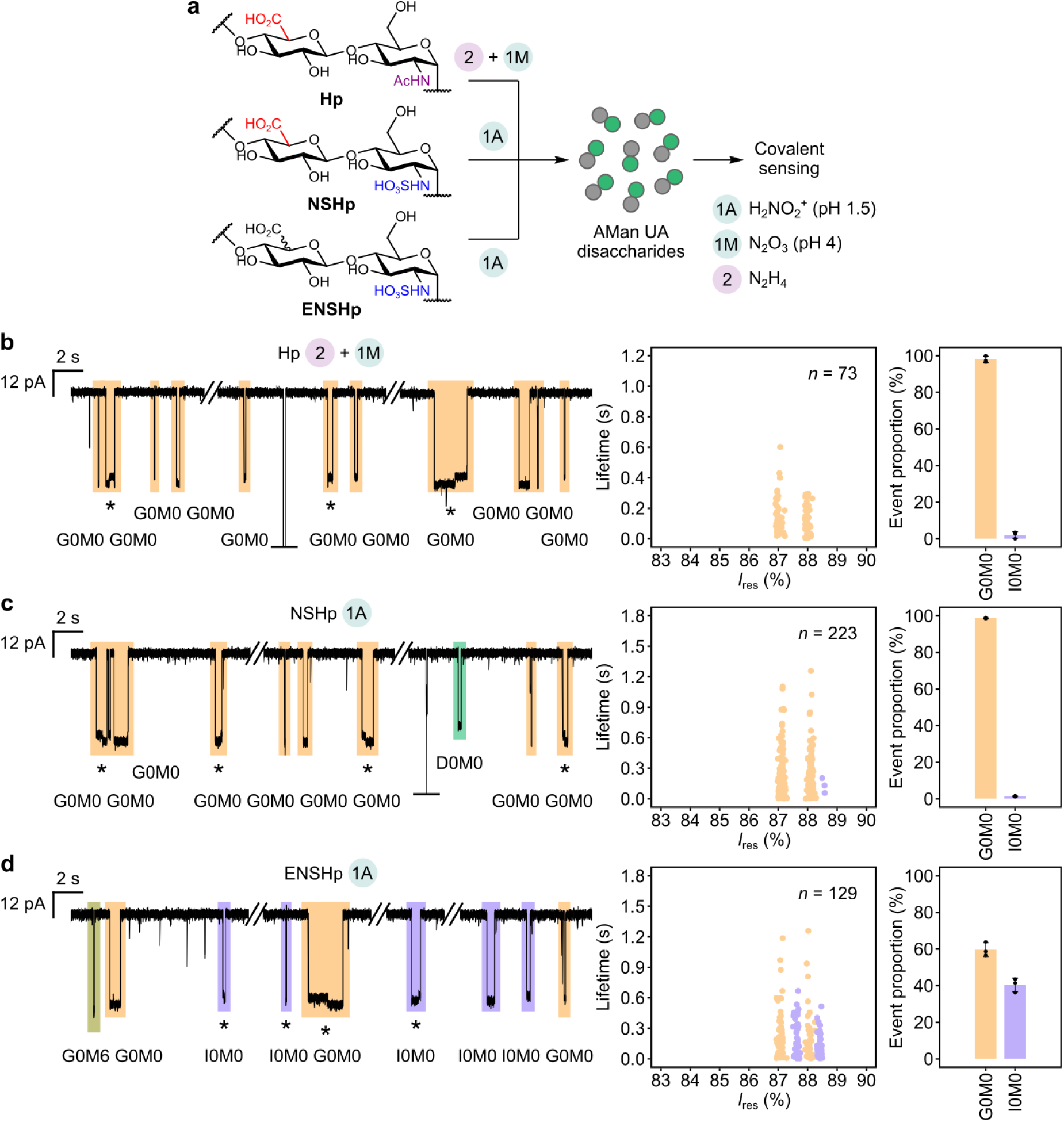
Deconvolutive compositional analysis of heparosan polysaccharides. **a,** Hp, NSHp, and ENSHp were deconvoluted via the modular application of hydrazinolysis and both ring contraction variants (conditions **2**, **1A**, and **1M**, with concomitant chain cleavage). The resulting product mixtures (∼0.3 mg) were directly introduced to the αHL-Cys117 pore. AMan and non-AMan residues are depicted as green and grey circles, respectively. **b–d,** The nanopore fingerprints of Hp (**b**), NSHp (**c**), and ENSHp (**d**) revealed disaccharide compositions and provided a direct, single-molecule readout of the degree of heparosan epimerization. Representative current traces are shown, along with assignments for G0M0, I0M0, D0M0 (rare), and G0M6 (rare); see Supplementary Methods for assignment criteria. Interconversion events are marked with an asterisk. Graphically truncated events are marked with a horizontal bar. The scatter plots show assigned G0M0 and I0M0 levels from single pores (*n* = number of events), and levels are colour-coded as in the traces. The bar plots show mean G0M0 and I0M0 event proportions across *N* = 3 pores; points indicate values from independent pores, and vertical error bars indicate s.d. **Recording conditions:** 4 M LiCl, 20 mM HEPBS, 40 μM EDTA, titrated to pH 8 using KOH; sugars (*cis*); +150 mV (*trans*); 23.8 ± 1 °C. Events with lifetimes under 2 ms were discarded (Methods).

Again, the resolving power of our approach enabled the discovery and precise characterization of unanticipated components. These included *O*-sulfation and eliminative (ΔUA) products, which exposed minor or even unforeseen chemical reaction pathways (Extended Data Figs 8d–f). For instance, both NSHp and ENSHp were obtained as synthetic commercial materials prepared from heparosan precursors via alkaline *N*-deacetylation optimized to minimize depolymerization^68,69^, followed by chemical *N*-sulfation, commonly regarded as exclusively *N*-selective^71^. Nonetheless, our NSHp and ENSHp fingerprints revealed D0M0 and G0M6, both unambiguously identified via paired interconversion events, suggesting their artefactual generation from base-catalysed eliminative cleavage and off-target *O*-sulfation, respectively (Extended Data Figs 8e–f). Notably, other than the ∼2 % abundance of ΔUA residues subsequently verified in NSHp using ^1^H NMR spectroscopy (Fig. S13), none of these side products were detected using a variety of classical techniques, including ^1^H and ^13^C NMR spectroscopy, size-exclusion chromatography (SEC), and HPLC disaccharide compositional analysis (Figs S15–16). Together, these results reinforced the utility of our workflow for compositional analysis and moreover demonstrated its capability to elucidate otherwise hidden molecular mechanisms by interrogating components that evade conventional ensemble methods.

### Detection of contaminants in adulterated pharmaceutical heparin

Heparin is a widely used clinical anticoagulant^19^. In 2008, a global healthcare crisis ensued when an adulterant in pharmaceutical heparin distributed across the US, Europe, and Asia was linked to increased occurrences of severe anaphylactoid reactions and deaths^72,73^. This adulterant was painstakingly identified as a chemically oversulfated form of the GAG chondroitin sulfate (OSCS)^74^. The close structural resemblance between OSCS and heparin, along with the antiquated tests for heparin quality control, had allowed OSCS to evade detection^73^. To this day, OSCS and other oversulfated polysaccharides, which are numerous and can elicit similar adverse reactions^75^, continue to pose threats to the global heparin supply chain^72,76,77^. While the current US Pharmacopeia (USP) heparin monograph mandates practical tests for OSCS detection—namely ^1^H NMR spectroscopy, strong anion-exchange HPLC, SEC, and assays that measure potency against specific coagulation factors^73^—these are targeted in scope and may struggle to detect other contaminants^76–78^. Alternative approaches based on, for instance, ^1^H NMR spectroscopy paired with multivariate modelling^79,80^, two-dimensional NMR spectroscopy^78,81^, LC-MS^76,82^, and IR spectroscopy^83^ are capable of identifying OSCS and other oversulfated polysaccharides. These approaches, however, currently rely on more centralized instrumentation and/or extensive operator expertise^73^; some can also demand substantial amounts of sample (e.g., 10–25 mg for NMR spectroscopy^78–81^).

To evaluate our workflow in the context of heparin quality control, we used a sample of pUFH after spiking to 20 % (w/w) with fully *N*-sulfated oversulfated heparosan (NSOSHp; Figs 5a, c), a structurally similar yet previously untested potential contaminant. This concentration closely approximated that of OSCS determined in multiple batches of adulterated heparin^76,84^. Unlike the case for OSCS^74^, the ^1^H NMR spectrum of NSOSHp (Fig. S17) lacks any signals clearly resolved from those of heparin, thus complicating its detection by USP monograph methods. This spiked sample was subjected to condition **1A** and probed using the αHL-Cys115 pore (Supplementary Methods). The resulting fingerprint immediately revealed a striking unrecognized event doublet that comprised 4.1 ± 1.1 % of all events (mean ± s.d. from *N* = 3 pores; Fig. 5d); this doublet was absent from the corresponding pUFH fingerprint (Extended Data Fig. 9b). Sizing and subsizing according to chain length and substructure, respectively, enabled the attribution of this doublet, with mean interconverting *I*_res_ values of 80.8 / 82.6 %, to an anhydrosugar disaccharide bearing three or four *O*-sulfate groups; coupling this with a logically deduced *N*-substituent identity (through the use of condition **1A**) revealed an *N*-sulfated precursor. Therefore, in a blind test, this collective information would allow a direct, rapid convergence to the small number of candidate contaminants containing *N*-sulfated tri- and/or tetra-*O*-sulfated disaccharide units—one of which is NSOSHp—from within broad GAG sequence space. A subsequent identical deconvolutive analysis of NSOSHp alone confirmed the unknown sugar as the tetrasulfated AMan disaccharide G5M9 (Extended Data Fig. 10a, Fig. S6, and Table S6). Notably, each such fingerprint was generated using only ∼0.3 mg of sample, and the full analytical workflow, from ring contraction to sensing, was completed in only ∼1.5 h. These results established proof of principle for the rapid detection and identification of a potential (here spiked and previously untested) contaminant.

**Fig. 5.**
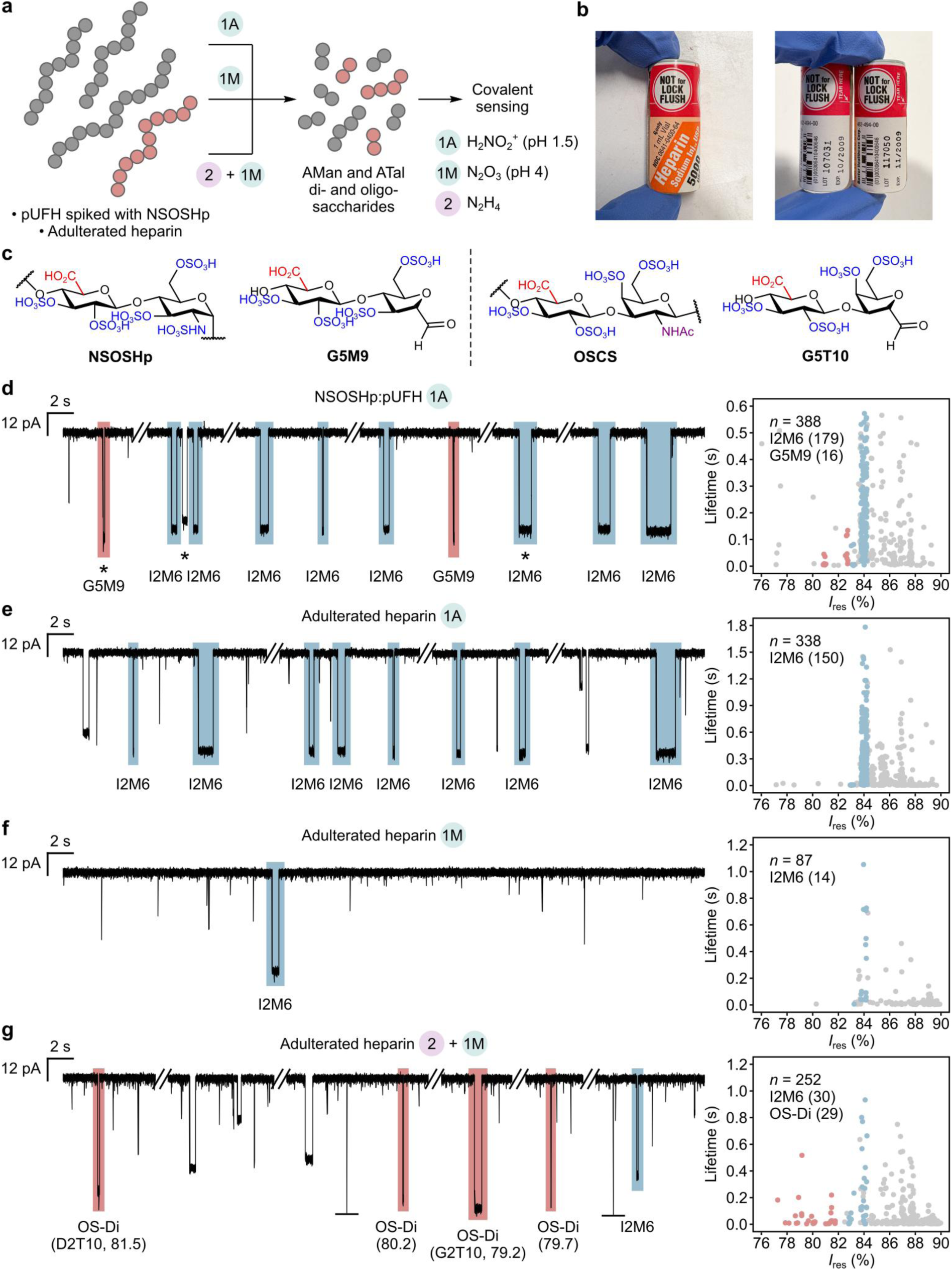
Detection of GAG contaminants in heparin. **a,** Conditions **1A**, **1M**, and **2** were differentially applied to two heparin samples: (i) pUFH spiked with NSOSHp to a final concentration of 20 % (w/w) and (ii) a pharmaceutical heparin sample withdrawn from the US market in 2008 due to adulteration. The resulting anhydrosugars (∼0.3–0.7 mg) were probed by the αHL-Cys115 pore (Supplementary Methods). Heparin and GAG contaminants are depicted in grey and red, respectively. **b,** Sealed vials of adulterated heparin; our data here correspond to lot 107031. **c,** The structures of NSOSHp, OSCS, and their corresponding anhydrosugar disaccharides G5M9 and G5T10, respectively. **d–g,** The fingerprints of the heparin samples revealed the structural signatures and therefore the presence of NSOSHp and OSCS. Representative current traces are shown for the NSOSHp:pUFH mixture (**d**) and the adulterated heparin sample (**e–g**), along with assignments for I2M6 (to guide the eye; see Supplementary Methods for assignment criteria) and G5M9. The *I*_res_ values of OSCS-derived sugars in the *I*_res_ 76–82 % cluster (OS-Di) are reported to one decimal place. G2T10 and D2T10 are tentatively assigned (see Extended Data Fig. 9e for OSCS data and Extended Data Fig. 10b for sugar structures and *I*_res_ predictions). Interconversion events are marked with an asterisk. Graphically truncated events are marked with a horizontal bar. The scatter plots in **d** and **e** show event levels from single pores. Due to the low GlcN and GlcNAc content of heparin, the scatter plots in **f** and **g** show combined event levels from two and three pores, respectively. Event levels in the scatter plots are colour-coded as in the traces, and levels from all other sugars are depicted in grey. Within the scatter plots, event counts for I2M6, G5M9, and OS-Di are given in parentheses. *n*, total number of events. **Recording conditions:** 4 M LiCl, 20 mM HEPBS, 40 μM EDTA, titrated to pH 8 using KOH; sugars (*cis*); +150 mV (*trans*); 23.8 ± 1 °C. Events with lifetimes under 2 ms were discarded (Methods).

Encouraged by this initial demonstration of utility, we next challenged our workflow with an archived pharmaceutical heparin sample withdrawn from the US market in 2008 due to adulteration (Fig. 5b). The parallel application of conditions **1A**, **1M**, and **2 + 1M**, followed by sensing using the αHL-Cys115 pore, yielded three complementary fingerprints (Figs 5e–g and Supplementary Methods). Under condition **1A**, the fingerprint of the adulterated sample proved essentially identical to that of an uncontaminated control (pUFH; Fig. 5e and Extended Data Fig. 9b); similarly matching fingerprints were generated under condition **1M** (Fig. 5f and Extended Data Fig. 9c). Together, these results allowed us to rule out the presence of any *N*-sulfated or *N*-unsubstituted contaminants.

By contrast, the use of conditions **2 + 1M** generated unrecognized characteristic events in the *I*_res_ 76–82 % range in the fingerprint of the adulterated sample but not in that of the pUFH control; these arose from multiple sugars and accounted for 11.0 ± 4.1 % of all events (Fig. 5g and Extended Data Fig. 9d). Sizing and subsizing, coupled with deductive logic afforded by the parallel application of three distinct chemistries, indicated a contaminant bearing *N*- acetylated tri- and/or tetra-*O*-sulfated disaccharide units, consistent with OSCS identified by alternative methods^74^. Once again, in a blind test, this collective information would allow a rapid convergence to a small number of candidate oversulfated contaminants, which in this instance includes OSCS.

We then subjected an OSCS standard to an identical workflow employing conditions **2 + 1M**. The resulting fingerprint closely resembled that of the adulterated sample, exhibiting a characteristic event cluster in the same *I*_res_ 76–82 % range corresponding to tri- and/or tetra- *O*-sulfated disaccharides; these comprised 12.6 ± 5.7 % of all events (Extended Data Fig. 9e). Moreover, this cluster contained individual events with *I*_res_ values closely matching those originating from the adulterated sample. These included events with single levels (singlets) at 79.1 % and events with interconverting levels (doublets) at 80.5 / 81.5 %, which we tentatively attributed to the trisulfated anhydrotalose (ATal) disaccharides G2T10 and D2T10, respectively, based on predicted *I*_res_ values (Extended Data Fig. 10b). Together, these results support a mode of ‘correlative ranging’, in which sugars in an otherwise unoccupied *I*_res_ range, rather than exhibiting specific values, are diagnostic of a contaminant.

The OSCS fingerprint also contained events throughout the *I*_res_ 82–90 % range, coinciding with the window where all pUFH-derived events resided (Extended Data Figs 9d–e). Sizing and subsizing indicated that these OSCS-derived events corresponded to various disaccharides bearing zero to two *O*-sulfate groups. Their heterogeneity in sulfation, together with the increased chemical shift dispersity of signals in the ^1^H NMR spectrum of OSCS post-hydrazinolysis (Fig. S18), suggested they arose as hydrazinolytic degradation products, consistent with a previous report^77^. Additional events arising from these degraded sugars could potentially enable a ‘combined ranging’ strategy to achieve even greater OSCS specificity. Importantly, no such degradation occurred for pUFH (Fig. S19).

The CS-type GAG dermatan sulfate is a common process-related impurity in heparin^85^. Therefore, as a test of impurity detection and a further test of ring-contractive deconvolution of non-HS GAGs, we also analysed a sample of porcine intestinal dermatan sulfate (DS) under conditions **2 + 1M** (Supplementary Methods). The resulting αHL-Cys115 fingerprint revealed two distinct event doublets (Extended Data Fig. 9f). Based on both predicted *I*_res_ values and the reported compositions of typical dermatan sulfates^86^, these doublets, with mean interconverting *I*_res_ levels of 84.9 / 87.0 % and 87.6 / 89.1 %, were tentatively attributed to the mono- and non-sulfated ATal disaccharides I0T4 and I0T0, respectively (Extended Data Fig. 10c). Notably, both ATal doublets were fully resolved from all previously characterized AMan standards. With these signatures established, we re-examined the corresponding adulterated heparin and pUFH fingerprints. While the former showed no evidence of dermatan sulfate–derived sugars, the latter revealed unambiguous interconverting I0T4, thus indicating low, perhaps even trace, levels of dermatan sulfate within the supplier-certified pure pUFH (Extended Data Fig. 9d). Strikingly, parallel attempts to corroborate this result using two-dimensional NMR spectroscopy—the gold standard in heparin quality control^78^—proved unsuccessful (Fig. S21).

Finally, as well as detecting the previously untested NSOSHp, our approach enabled us to readily rule out the presence of other untested contaminants. In 2010, the precise nature of the adulterant in 2008 US heparin products was called into question, when two studies claimed to have detected not only OSCS but also oversulfated HS (OSHS)^82,87^. Under our workflow, the presence of certain sugars (e.g., G5M9 generated under condition **1A**) would unambiguously signal OSHS adulteration; however, no such sugars were detected within the authentic crisis-associated heparin sample in our possession (Fig. 5e). This result is consistent with a 2014 investigation confirming that, for samples within this specific heparin lot, OSCS was the sole adulterant^76^.

Thus, by deploying two distinct analytical modes—(i) predictive and direct event matching and (ii) predictive and correlative ranging—our workflow enabled us to rapidly survey contaminated heparins of diverse origin, including an authentic historical adulterated pharmaceutical sample, and detect or exclude contaminants using minimal material. Across all evaluated samples, generating a full set of fingerprints required at most 1.5 mg of material, and entire analytical workflows were completed in only ∼1.5–5.5 h.

## Discussion

Nanopore sensing has revolutionized genomics^35^ and is projected to similarly transform proteomics^88,89^. Its extension to glycomics is therefore tantalizing. Whereas previous nanopore approaches have highlighted the challenge of signal transience inherent in glycan analytes (see refs ^41,90^ for early examples), here we have overcome this bottleneck through a strategy of deconvolutive glycan ring contraction that generates powerfully electrophilic aldehydes for covalent sensing. In this way, we have resolved, for the first time in GAGs, individual sugar residues alongside their underlying fine structural features.

One next opportunity is to resolve a more comprehensive set of oligosaccharides, as functional GAG–protein interactions typically involve tetrasaccharide or higher-order sequence motifs^3,9,10^. Beyond this, elucidating their linear order, spacing, and frequency within a given GAG chain would ultimately require the sequencing of intact polysaccharides by processive reading. An immediate challenge lies in developing a ratcheting mechanism to control polysaccharide translocation, for which methods based on unassisted electrophoresis^36–44^ have consistently failed to deliver the prolonged, millisecond-to-second residence times that, as we have shown here, enable structural resolution; tothis end, various solutions based on either dynamic covalent or non-covalent chemistries (e.g., enzymeless disulfide-based ‘hopping’^52^ and helicase ratcheting of DNA conjugates^45,91^) may be considered. Although such long reads have not been the focus of this work, our ability to pinpoint fine structural features in *immobilized* di- and oligosaccharides—such as the position of sulfate groups and single-atom stereochemistry—represents a necessary first step towards doing so in *moving* polysaccharides. Moreover, ring contraction furnishes reactive aldehydes that enable facile conjugation^92–94^ at rates up to two orders of magnitude above those of conventional reducing-end strategies^58^; therefore, through the use of conditions that induce minimal polysaccharide cleavage, our approach is uniquely suited for integration with various ratcheting mechanisms that require adaptor installation^52,91^.

Limitations of our approach include the risk of relying on potentially degradative chemistries to deconvolute NAc sugars and the relatively low rates of covalent sensing (e.g., rate constants governing thiohemiacetal formation between the αHL-Cys115 pore and a representative panel of non-, mono-, and disulfated disaccharides were in the range 0.16– 0.34 mM^−1^ s^−1^; Extended Data Fig. 11 and Supplementary Methods). First, while we have determined, using NMR spectroscopy and MS, that all reactions employed here (**1A**, **1M**, and **2**) preserve the fine structural features of HS (Supplementary Methods)—consistent with historical bulk fractionation data for HS^61,62^ and many other GAGs^86^—a small subset of structures may prove less stable. Should degradation occur, as observed here during OSCS hydrazinolysis, milder reactions (e.g., direct *N*-nitrosation and subsequent ring contraction of NAc sugars^49^) can be explored. Second, the relatively low rates of covalent sensing currently limit throughput and mass sensitivity. For each experiment, a single nanopore was used alongside tens to hundreds of micrograms of analyte to produce sufficient events for analysis within a 0.5–1.5 h window. By comparison, a state-of-the-art ion mobility MS platform for resolving HS ΔUA disaccharides requires only tens of nanograms of analyte and several minutes for data acquisition^28^. This disparity, however, also reflects the limitations inherent to the use of single nanopores, which can be bypassed by multi-pore parallelization (see below).

The toxicity of numerous oversulfated polysaccharides^75^, as contaminants in pharmaceutical heparin, calls for detection or, preferably, identification methods with a broad scope. Here, sugar sizing and subsizing coupled with deductive chemical logic revealed characteristic structural information that enabled the detection of OSCS and the identification of an additional potential contaminant, NSOSHp. In a demonstration of real-world utility, we detected and excluded contaminants in a pharmaceutical heparin sample withdrawn during the 2008 global heparin contamination crisis. Moreover, our resolution of multiple contaminants and a process impurity (dermatan sulfate) across various heparins suggests the feasibility of simultaneous multi-GAG screening. Thus, our work builds upon an earlier attempt to detect contaminants using solid-state nanopores, which, while proving successful, was limited to a simulant 1:1 heparin–OSCS mixture and was unable to identify either polysaccharide^37^. Although several other general and consequently powerful methods exist for contaminant identification^76,78–83^, these remain underutilized due to practical challenges relating to instrument size, operator expertise, and cost (see Extended Data Table 1 for a comparison of various methods). By contrast, our approach is simple in both execution and data interpretation, and is adaptable to low-cost, portable nanopore sequencing devices such as the MinION^35^. Such adaptation will also enable parallelization across hundreds of pores to drastically enhance throughput and mass sensitivity; even in our current single-pore setup, assay times and material requirements already match those of alternative approaches (∼1.5–5.5 h in total for sample preparation and sensing; ∼0.3–0.7 mg per sensing run).

At the heart of our deconvolutive strategy is a reduction in the number of unique GAG structures that must be interrogated by nanopores, whereby multiple precursors can be chemoselectively converted into the same anhydrosugar. Notably, our resolved panel of seven UA and four ΔUA AMan disaccharides covers 29 disaccharide units and their eliminative digestion variants in natural HS (Extended Data Fig. 3); this count closely approaches the theoretical maximum of 33, reflecting 11 anhydrosugars each mapping to three natural *N*-substituent precursors (NS, N, and NAc)—the four remaining disaccharides are theoretically plausible but have yet to be found in nature. Consequently, this establishes, to our knowledge, the largest number of HS disaccharides tractable by a single analytical method reported to date (Supplementary Discussion 3). Including the NSOSHp-derived, and therefore non-natural, AMan disaccharide G5M9 further extends this ceiling of tractable HS-type disaccharides to 36. This expanded analytical capability—together with an unprecedented power to size and even subsize sugar molecules according to chain length and substructural features in a fractionation-free manner—paves the way for the discovery of novel oligosaccharide structures directly within native biological mixtures.

While GAGs are vital in animal physiology and disease, they are but one class of aminosugar biopolymer. Present across all domains of life, aminosugars have few parallels in structural breadth^33^. The exceptional chemoselectivity and functional group compatibility of hydrazinolysis^62^ and the ring contraction variants^61^ suggest a true generality of scope, which would encompass other aminosugar biopolymers—including chitin, chitosan, *N*- and *O*-type glycans (as part of glycoproteins, glycoRNA, and glycolipids), milk oligosaccharides, and bacterial capsular and lipopolysaccharides^33^. For instance, the conditions developed here could be directly applied to selectively cleave *N*-glycans from glycoproteins^95,96^ to generate corresponding reactive sugars for sensing. The exquisite sensitivity of covalent sensing to GAG molecular shape that we have demonstrated here should prove similarly powerful in the characterization of *N*-glycans, which often occur as complex mixtures of constitutional (branched) isomers^33^. Paired with devices such as the MinION, our approach paves the way for the long-sought ‘democratization’^33^ of glycan analysis, which will reveal new structure– function correlations and drive discoveries in fundamental biology and medicine.

## Methods

### Nanopores

αHL pores were prepared according to an *in vitro* transcription–translation (IVTT) protocol adapted from the literature^1^. The αHL-Cys115 and αHL-Cys117 pores are heteroheptameric constructs comprising six A subunits and one cysteine-bearing B subunit. All A and B subunits carry K8A, M113G, K131G, and K147G mutations, whereas B subunits uniquely carry an additional T115C or T117C mutation alongside a C-terminal octa–aspartic acid tag. A commercial IVTT system (*E*. *coli* T7 S30 Extract System for Circular DNA, Promega) was used. For each pore variant, a 50 µL reaction was set up according to supplier instructions with 3.56 μg of plasmid encoding the A subunit and 0.44 μg of plasmid encoding the B subunit (∼8:1 plasmid molar ratio). Following the addition of 1–2 μL of ^35^S methionine (15 mCi mL^−1^, MP Biomedicals) and 1 μL of rifampicin solution (5 ng μL^−1^) to inhibit endogenous *E*. *coli* RNA polymerase, the mixture was incubated at 37 °C for 1 h (400 rpm). To induce pore assembly, 3 µL of rabbit erythrocyte membrane solution (3 mg protein mL^−1^) was added, followed by incubation at 37 °C for a further 1 h (400 rpm). The mixture was then subjected to two sequential rounds of centrifugation at 18,000 × *g* for 15 min and pellet resuspension in 200 μL of MBSA buffer (10 mM MOPS, 150 mM NaCl, 1 mg mL^−1^ BSA, pH 7.4). Following a final centrifugation at 18,000 × *g* for 15 min and removal of the supernatant, the resulting pellet was resuspended in 30 μL of 2.5 % βME in Laemmli buffer without heating and electrophoresed on a 5.5 % polyacrylamide gel (Tris·HCl, 0.1 % SDS, pH 8.8) of ∼20 cm height at 70 V and room temperature for 16 h. Sodium thioglycolate and DTT were added to the electrophoresis buffer to final concentrations of 1 mM and 2 mM, respectively. The buffer was flushed with inert gas before use, and electrophoresis was performed under an inert atmosphere.

Afterwards, the gel was placed on filter paper, vacuum-dried at room temperature for 6 h, and exposed to photographic film for 6–12 h. The film was developed, and the band that migrated the second slowest in the 100–150 kDa region, corresponding to the desired αHL- Cys115 or αHL-Cys117 variants, was excised. The excised product was rehydrated in 200 µL of TE buffer (10 mM Tris, 0.1 mM EDTA, 0.5 mM DTT, pH 8), homogenized, and incubated at room temperature for 2 h. The mixture was then passed through a 0.2-μm- diameter membrane filter (Proteus Mini Clarification Spin Column, Generon) at 14,000 × *g* and 4 °C. The filtrate was aliquoted into 10 µL portions, flash-frozen with liquid N_2_, and stored at −80 °C until use.

The αHL-A_7_ pore, used in control experiments (Fig. S4), is a cysteine-free homoheptameric pore comprising seven A subunits and was isolated as a byproduct during the preparation of the cysteine-bearing pore variants. For this purpose, the gel band that migrated the slowest in the 100–150 kDa region was excised. Subsequent isolation and purification steps were carried out as detailed above.

### Sugars

Unless stated otherwise, all sugars were purchased as sodium salts and used as received. The following sugars were purchased from Iduron: ΔUA2S-GlcNS6S (D2S6, HD001, batch numbers 5–7), ΔUA2S-GlcNS (D2S0, HD002, batch numbers 5–6), ΔUA- GlcNS6S (D0S6, HD004, batch number 16), ΔUA-GlcNS (D0S0, HD005, batch numbers 5 and 7), 2-*O*-desulfated heparin (2DSH, DSH001/2, batch number 1), and 6-*O*-desulfated heparin (6DSH, DSH002/6, batch number 1). The following sugars were purchased from Glycan Therapeutics: 6S 7-Mer C (GT77-AZ-021, lot GT221-VP070516), 2S 7-Mer A (GT77- AZ-027, lot HS-20250702-GS), and 6S IdoA 7-Mer B (GT77-AZ-083, lot GT77-283-VP051117). The following sugars were purchased from GlycoNovo: heparosan (Hp, C- LMWK5PS, lot 1B60H1H, mass-average molecular mass = 12.2 kDa, number-average molecular mass = 9.6 kDa), fully *N*-sulfated heparosan (NSHp, C-CNSK5PS, lot 1B30C2H, mass-average molecular mass = 59.2 kDa, number-average molecular mass = 57.9 kDa), epimerized fully *N*-sulfated heparosan (ENSHp, C-EPICNSK5PS, lot 1B60H1H, molecular mass = 84 kDa), and *N*-sulfated oversulfated heparosan (NSOSHp, C-CNOSK5PS, lot 2B1OI1C, molecular mass = 85.3 kDa). The following sugars were purchased from Sigma-Aldrich: glucosamine hydrochloride (GlcN, G1514) and porcine intestinal dermatan sulfate (DS, C3788, lot 0000417795). Unfractionated porcine intestinal heparin (pUFH) was purchased from Alfa Aesar (A16198, lot 10198820). Fondaparinux was purchased from Biosynth (OF04094, lot 040941550). Oversulfated chondroitin sulfate (OSCS) was purchased from the US Pharmacopeia (1133580, lot R188K0). Sealed vials of adulterated pharmaceutical heparin (Heparin Sodium Injection, 1 mL vial, Baxter Healthcare, 5,000 US Pharmacopeia units mL^−1^, lots 107031 and 117050) were gifted by Robert J. Linhardt and Fuming Zhang. In this work, only lot 107031 was analysed; the concentration of OSCS in this specific lot was previously determined to be 19.5 % (w/w) (ref. ^2^). All sugars were stored at −20 °C until use.

### General procedure for hydrazinolysis (condition 2)

This was adapted from a literature protocol^3^. 1 mg of sugars was dissolved in 66.7 µL of freshly prepared hydrazine reagent (1 % N_2_H_4_·H_2_SO_4_ in N_2_H_4_·H_2_O) in a polypropylene microcentrifuge tube. **Caution:** Hydrazine and its fumes are highly toxic; all handling and reaction steps should be performed within a fume hood. The tube was flushed with inert gas, tightly sealed, and the solution incubated at 100 °C for 4 h (1,000 rpm). Following incubation, the mixture was cooled to room temperature under open air, and lyophilized to yield *N*-deacetylated sugars containing residual N_2_H_4_ as an amorphous yellow or brown cake.

### General procedure for ring contraction at pH 1.5 (condition 1A)

This was adapted from a literature protocol^4^. 1 mg of sugars, alongside stock solutions of 0.5 M H_2_SO_4_ and 1 M NaNO_2_, was placed on ice. 500 µL of NaNO_2_ solution was added to 500 µL of H_2_SO_4_ solution; the mixture (pH 1.5) was swirled for 3 s, then a 60 µL aliquot was immediately transferred to the sugars. **Caution:** The mixing of NaNO_2_ and H_2_SO_4_ generates toxic HNO_2_ fumes; this step should be performed within a fume hood. The reaction mixture was protected from direct light and allowed to warm to room temperature over 10 min, followed by the addition of 10.5 µL of 2 M Na_2_CO_3_ to quench the reaction and adjust its pH to 8. The resulting solution was either taken immediately for nanopore sensing or stored at −80 °C until use.

### General procedure for ring contraction at pH 4 (condition 1M)

This was adapted from a literature protocol^4^. 500 µL of 5.5 M NaNO_2_ solution was added to 200 µL of 0.5 M H_2_SO_4_ solution; the mixture (pH 4) was swirled for 3 s, then a 40 µL aliquot was immediately added to 1 mg of sugars. **Caution:** The mixing of NaNO_2_ and H_2_SO_4_ generates toxic HNO_2_ fumes; this step should be performed within a fume hood. When performed on *N*-deacetylated sugars containing residual N_2_H_4_, 3 M H_2_SO_4_ (typically 3–4 µL) was further added to adjust the pH to 4. The solution was protected from direct light and allowed to react at room temperature for 10 min, followed by the addition of 4.3 µL of 2 M Na_2_CO_3_ to quench the reaction and adjust its pH to 8. The resulting solution was either taken immediately for nanopore sensing or stored at −80 °C until use.

### Eliminative digestion using heparinases (condition 3)

Heparin polysaccharides (pUFH, 2DSH, and 6DSH) were digested according to a protocol adapted from the literature^5^. Recombinant *Pedobacter heparinus* heparinases I, II, and III in their lyophilized forms were purchased from IBEX Pharmaceuticals (60-010, 60-018, and 60-020, respectively; *Pedobacter heparinus* was formerly known as *Flavobacterium heparinum*). The heparinases were each reconstituted in H_2_O to a concentration of 2,000 mU µL^−1^, aliquoted, stored at −80 °C, and used immediately after thawing. The heparins were each dissolved in H_2_O to a concentration of 100 µg µL^−1^. Digestion reactions were prepared as follows:

| Component | Volume (µL) |
| --- | --- |
| pUFH, 2DSH, or 6DSH (1 mg) | 10 |
| Heparinase I (4,000 mU) | 2 |
| Digestion buffer (10 mM NaP<sub>i</sub> pH 7) | 100 |

The mixture was initially incubated at 37 °C for 2 h. Next, heparinase III (4,000 mU) was added, followed by a further incubation at 37 °C for 2 h. Heparinase II (4,000 mU) was subsequently added, after which the mixture was incubated at 37 °C for 24 h. Finally, the mixture was incubated at 100 °C for 2 min to denature all heparinases. A small aliquot (∼0.4 µL) was analysed on HPTLC silica gel 60 aluminium sheets (Sigma-Aldrich) according to a literature protocol^6^. Under these digestion conditions, only single spots corresponding to disaccharides were observed, indicating exhaustive digestion (Fig. S10d, but see main text and Extended Data Figs 8a–c). The remaining mixture was lyophilized and subjected to ring contraction as detailed above.

### Nanopore sensing and data analysis

1,2-Diphytanoyl-*sn*-glycero-3-phosphocholine (DPhPC) was purchased from Avanti Polar Lipids (850356). Ag:AgCl electrodes were prepared by joining copper wires with silver wires, followed by bleaching of the exposed silver for 24 h. These electrodes were protected from direct light and equipped with fresh salt bridges of 3 M KCl and 3 % low-melt agarose before use. Electrolyte solutions were similarly protected from light and stored at 4 °C until use.

All nanopore recordings were conducted in a Faraday cage on an anti-vibration table at 23.8 ± 1 °C. Planar lipid bilayer membranes were prepared using the Montal–Mueller technique^7^. A 25-μm-thick Teflon film with an aperture 70–80 μm in diameter was fixed between two Delrin compartments. The aperture was coated with 1 % hexadecane in pentane (1–2 µL) on each side and left to dry for 5 min. Each compartment was subsequently filled with 500 μL of electrolyte (4 M LiCl, 20 mM HEPBS, 40 μM EDTA, titrated to pH 8 using KOH). Following electrode insertion into each compartment, 0.25 % DPhPC in pentane (1–2 µL) was added to both sides, and the solutions were gently mixed to allow the self-assembly of a planar lipid bilayer across the aperture. A small volume (1–2 µL) of αHL protein solution was then introduced into the grounded compartment. Upon the insertion of a single pore into the bilayer, the grounded compartment (here referred to as *cis*) was transfused with fresh electrolyte to remove superfluous protein. Following a brief assessment of the current– voltage characteristics and stability of the pore (see Fig. S1 for the open-pore characteristics of the αHL-Cys115 and αHL-Cys117 variants), analyte solution (typically 10–25 µL) was added to the *cis* compartment, and the solution was gently mixed for ∼3 min before data acquisition at +150 mV.

Current traces were generated with a patch-clamp amplifier (Axopatch 200B, Molecular Devices) and processed with a digitizer (Axon Digidata 1440A, Molecular Devices). Signals were acquired at an output gain of 10×, sampled at 100 kHz, and filtered at 10 kHz using an internal 4-pole low-pass Bessel filter. Data were transferred to Clampfit (Molecular Devices, v10.7), further filtered at 500 Hz using an 8-pole low-pass Bessel filter, and idealized using the Single-Channel Search function. The open-pore current was determined by fitting a single-term Gaussian distribution to an all-points histogram using the Fit function with default parameters, and updated manually when necessary. The relative amplitudes of all other levels were held constant, and events with lifetimes of 2 ms or less were discarded. An exception was made when M0 was analysed using the αHL-Cys117 pore: transferred data were instead filtered at 250 Hz using an 8-pole low-pass Bessel filter, and events with lifetimes of 4 ms or less were discarded to minimize artefacts at this reduced bandwidth. The residual current was defined as *I*_res_ = 100 × (*I*_e_/*I*_o_), where *I*_e_ and *I*_o_ denote the event amplitude found by Single-Channel Search and the open-pore current, respectively. When benchmarking individual AMan sugars, a small amount of D0M0 (exhibiting complete *I*_res_ resolution) was added and used as an internal standard (e.g., see Figs S23–25). Rate constants were extracted from idealized lifetime data using QuB (www.qub.buffalo.edu; Extended Data Fig. 11 and Supplementary Methods)^8,9^.

## Acknowledgements

This work was funded by Oxford Nanopore Technologies, the Gates Foundation (INV- 074158), a European Research Council Starting Grant under the UK Research and Innovation Horizon Europe Guarantee scheme (EP/Z000351/1; to Y.Q.), and a Croucher Scholarship (to L.L.M.S.). Next Generation Chemistry at the Rosalind Franklin Institute has been funded by grants from the Engineering and Physical Sciences Research Council (EP/V011359/1, EP/T012021/1, EP/T012005/1, EP/V011367/1, and EP/X527245/1). We thank Robert J. Linhardt and Fuming Zhang for their provision of adulterated heparin, Coral Mycroft for assistance with NMR spectroscopy, Victor Mikhailov for assistance with MS, and Georgios Papadakis, Zhong Hui Lim, Dimitrios Mamalis, Anthony J. Devlin, Lan Na, and Ajay Jha for discussions.

## Author contributions

L.L.M.S., H.B., B.G.D., and Y.Q. conceived the work and designed experiments. L.L.M.S., A.Y., and D.P.C. performed experiments and analysed data. L.L.M.S., H.B., B.G.D., and

Y.Q. wrote the manuscript. All authors contributed to the final manuscript.

---

**Extended Data Fig. 1.**
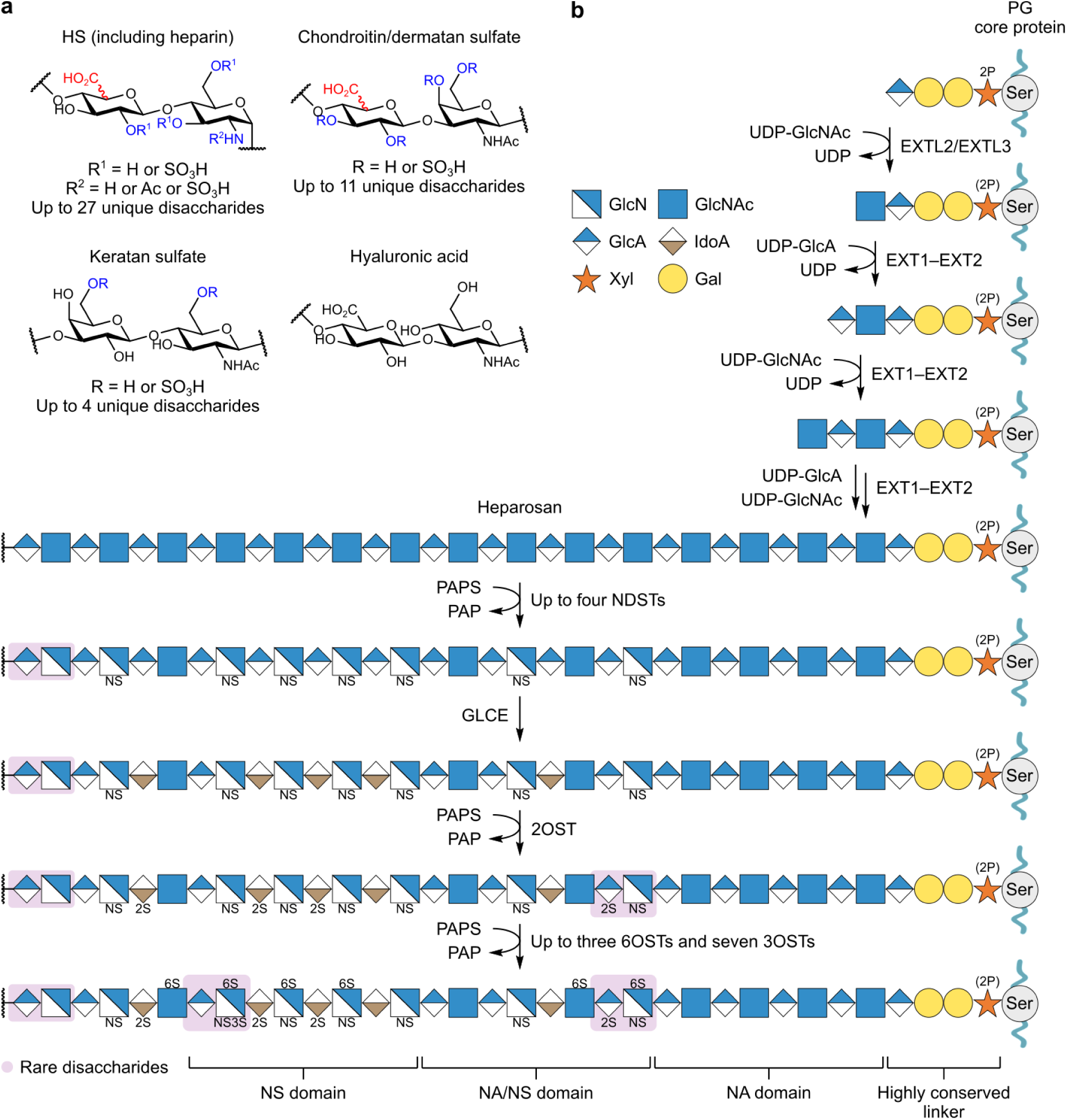
Natural GAG structural diversity and HS biosynthesis. **a,** HS, chondroitin/dermatan sulfate, and hyaluronic acid GAGs consist of repeating disaccharide units in which the first residue is either GlcA or its C5 epimer, IdoA, and the second residue is either GlcN in one of three *N*-substitution states (NS, N, or NAc) or *N*- acetylgalactosamine^1^. By contrast, keratan sulfate GAGs comprise Gal-GlcNAc disaccharide units (Gal = galactose)^1^. While HS, chondroitin/dermatan sulfate, and keratan sulfate are sulfated at various residue positions in a typically heterogeneous fashion, hyaluronic acid is not sulfated^1,33^. HS represents the most structurally diverse GAG subclass, with up to 27 unique disaccharide units emerging from its biosynthesis (Table S1). **b,** HS is synthesized on highly conserved oligosaccharide linkers on proteins as part of proteoglycans (PGs)^1–3,33^. Within the Golgi apparatus, chains of repeating GlcAβ1-4GlcNAcα1-4 disaccharide units (heparosan) are initially assembled on these linkers via the stepwise addition of GlcA and GlcNAc by the glycosyltransferases EXT1–EXT2 and EXTL3. A subset of GlcNAc residues is then converted to GlcNS via tightly coupled deacetylation and sulfation reactions catalysed by *N*-deacetylase-*N*-sulfotransferases (NDSTs)^3^; occasional decoupling of these reactions results in unsubstituted GlcN residues^3^. NDSTs typically generate clusters of GlcNS residues that direct subsequent enzymatic elaboration: the epimerase GLCE converts GlcA residues to IdoA, while up to 11 distinct sulfotransferases modify the 2-*O*-positions of GlcA/IdoA residues and the 3- and 6-*O*-positions of GlcNS/GlcNAc/GlcN residues^1^. The resulting mature, heterogeneous HS chains exhibit regions rich in GlcA/GlcNAc (NA domains) and those rich in IdoA/GlcNS and their highly sulfated derivatives (NS domains), flanked by sequences of intermediate modification (NA/NS domains). Notably, chain polymerization and *N*-sulfation may be coupled; moreover, the sequence of subsequent epimerization and sulfation reactions—although depicted here in order of broad enzymatic preference—is not strictly defined^3^. Ser, serine; Xyl, xylose; 2P, 2-phospho; UDP, uridine 5′- diphosphate; PAPS, 3′-phosphoadenosine-5′-phosphosulfate; PAP, 3′-phosphoadenosine-5′- phosphate; 2OST, 2-*O*-sulfotransferase; 6OST, 6-*O*-sulfotransferase; 3OST, 3-*O*- sulfotransferase.

**Extended Data Fig. 2.**
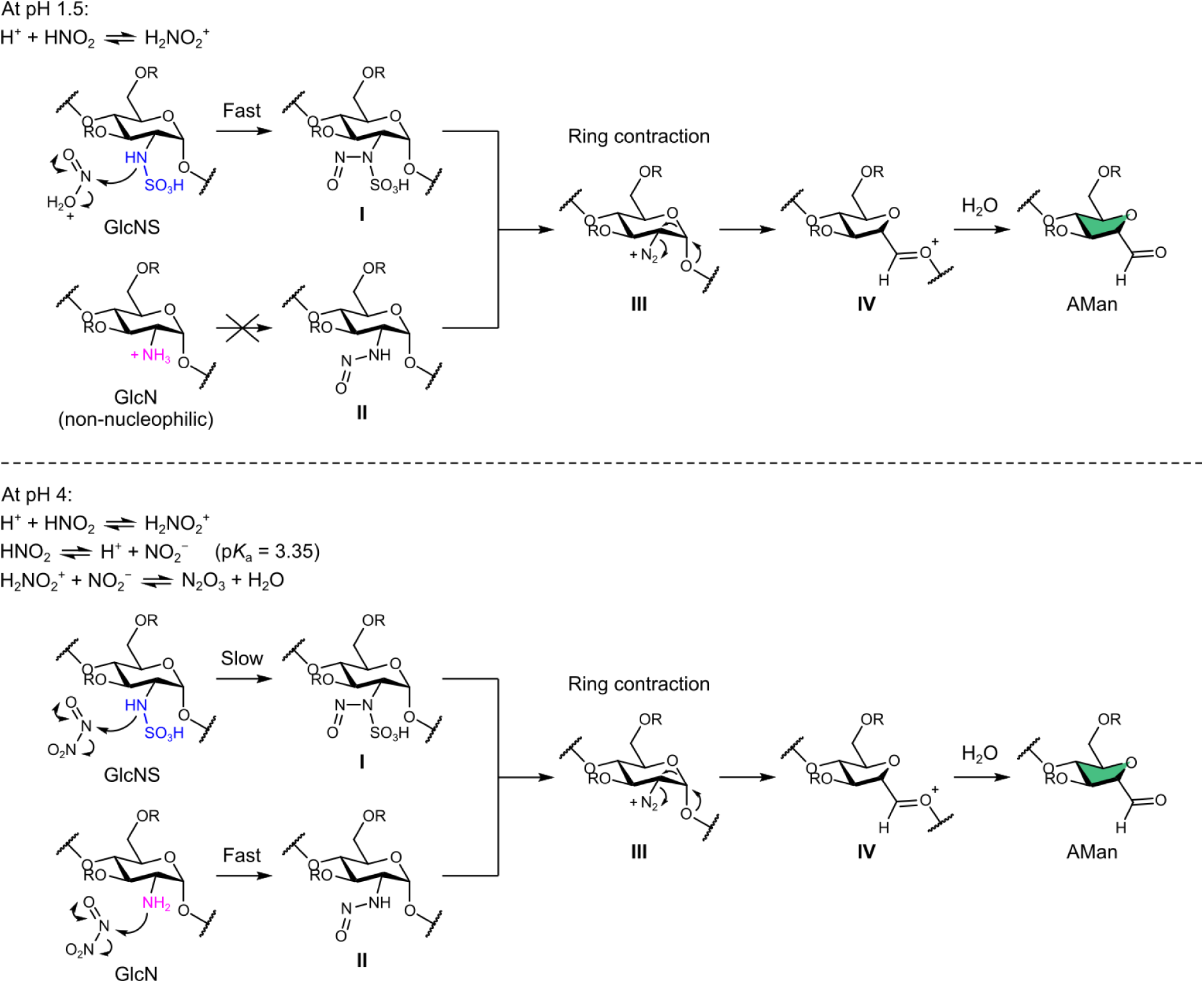
Mechanism of the ring contraction reaction and origin of chemoselectivity. The overall reaction is initiated by the rate-determining *N*-nitrosation of GlcNS and GlcN residues, producing intermediates **I** and **II**, which rapidly collapse to a common diazonium intermediate **III** (refs ^47,61^). This intermediate then undergoes a Tiffeneau–Demjanov rearrangement^47^ (which may proceed through the concerted mechanism depicted here, or via a stepwise pathway) to form oxocarbenium **IV**, which is subsequently hydrolysed to AMan. The chemoselectivity of the overall reaction arises from pH-dictated differences in the rates of *N*-nitrosation^61^. Under condition **1A** (pH 1.5), the strongly electrophilic nitrosating agent H_2_NO_2_^+^ is generated and reacts rapidly with the GlcNS nitrogen. Here, the GlcN nitrogen is essentially fully protonated and therefore non-nucleophilic. Under condition **1M** (pH 4), the alternative nitrosating agent, N_2_O_3_, is generated; this agent reacts rapidly with the GlcN nitrogen, which is less protonated than at pH 1.5 and therefore exhibits superior nucleophilicity. R = H or SO_3_H.

**Extended Data Fig. 3.**
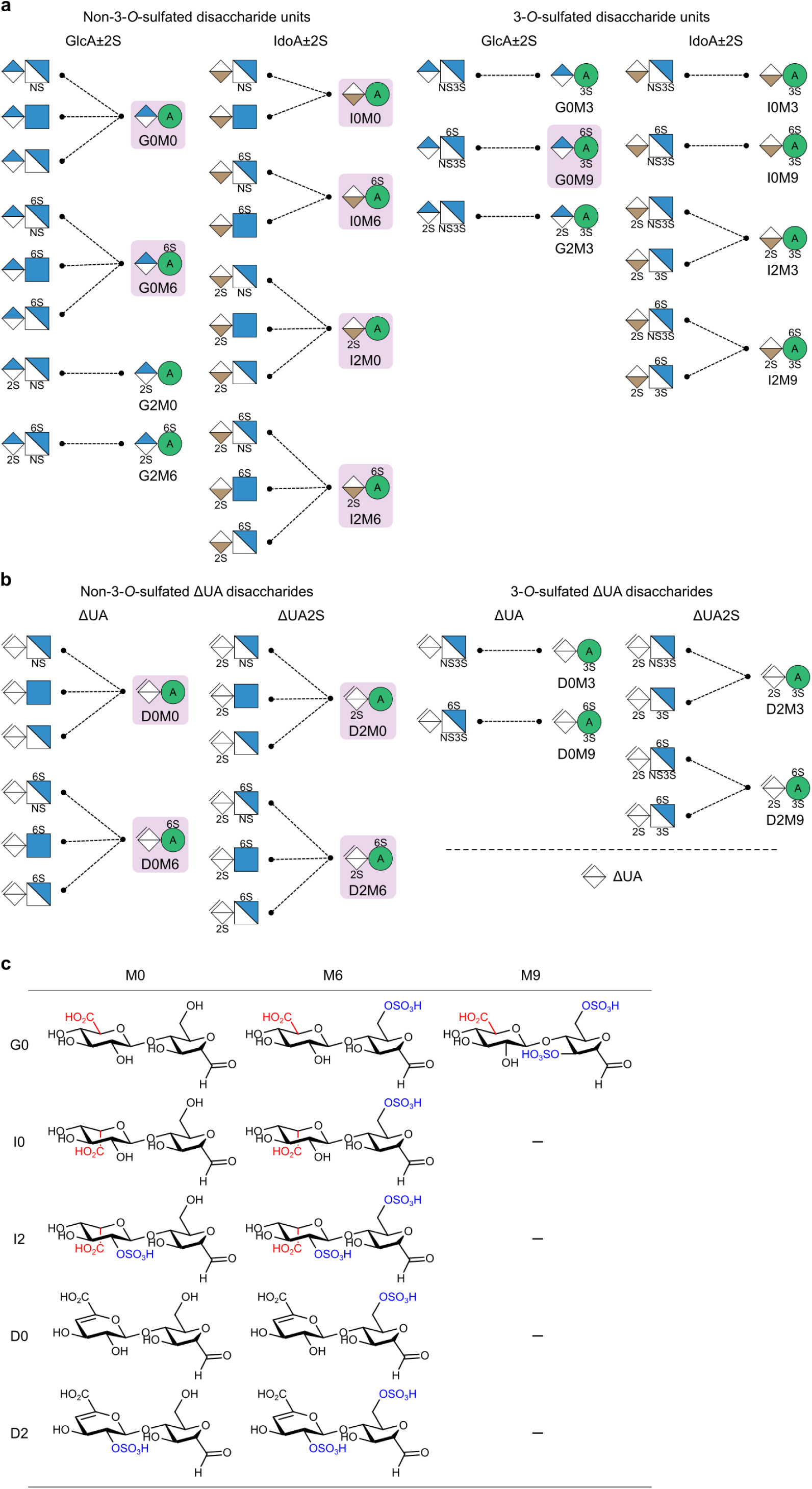
GAG sequence deconvolution at the disaccharide level. **a,** The 27 unique disaccharide units that emerge from HS biosynthesis (Table S1) can be deconvoluted into 15 AMan disaccharides. **b,** Similarly, the 18 ΔUA disaccharides that emerge from the eliminative digestion of HS can be deconvoluted into eight AMan ΔUA disaccharides. For **a** and **b**, the AMan disaccharides examined in this work are highlighted in pink. **c,** Structures of the examined disaccharide standards.

**Extended Data Fig. 4.**
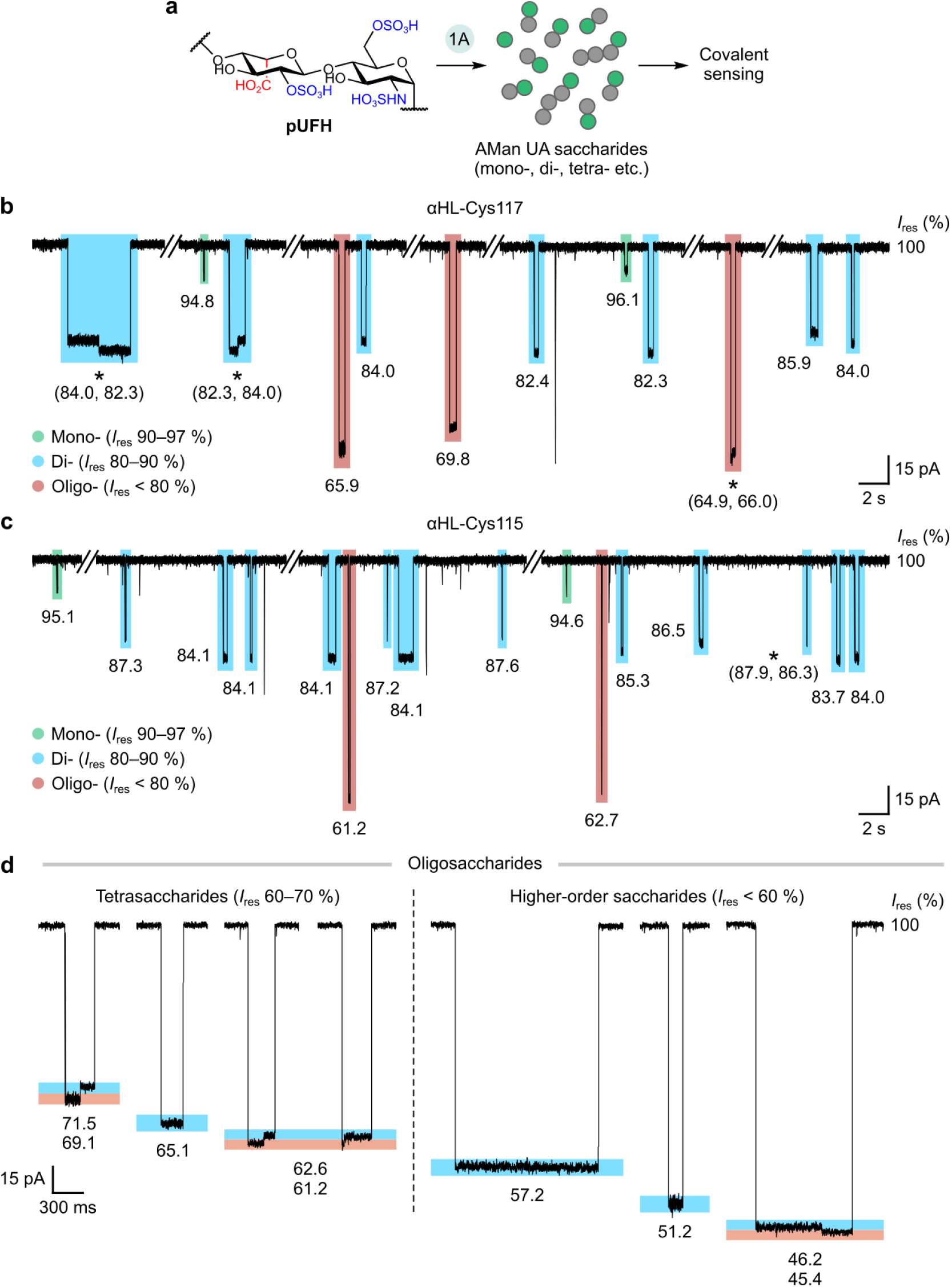
Direct, real-time sugar sizing within complex mixtures. **a,** pUFH (representative structure shown) was ring-contracted at pH 1.5 (condition **1A**) and subsequently probed without further fractionation. AMan and non-AMan residues are depicted as green and grey circles, respectively. **b–c,** Current traces for αHL-Cys117 (**b**) and αHL-Cys115 (**c**) both revealed discrete event clusters of mono-, di-, and oligosaccharides (see Table S7 for event proportions). Event *I*_res_ values are listed; interconversion events are marked with an asterisk. **d,** Representative events within the oligosaccharide cluster probed using the αHL-Cys115 pore are shown alongside their *I*_res_ values (levels highlighted in blue and pink). **Recording conditions:** 4 M LiCl, 20 mM HEPBS, 40 μM EDTA, titrated to pH 8 using KOH; sugars (*cis*); +150 mV (*trans*); 23.8 ± 1 °C. For each experiment, ∼0.3 mg of ring-contracted pUFH was used. Events with lifetimes under 2 ms were discarded (Methods). All *I*_res_ values are reported to one decimal place.

**Extended Data Fig. 5.**
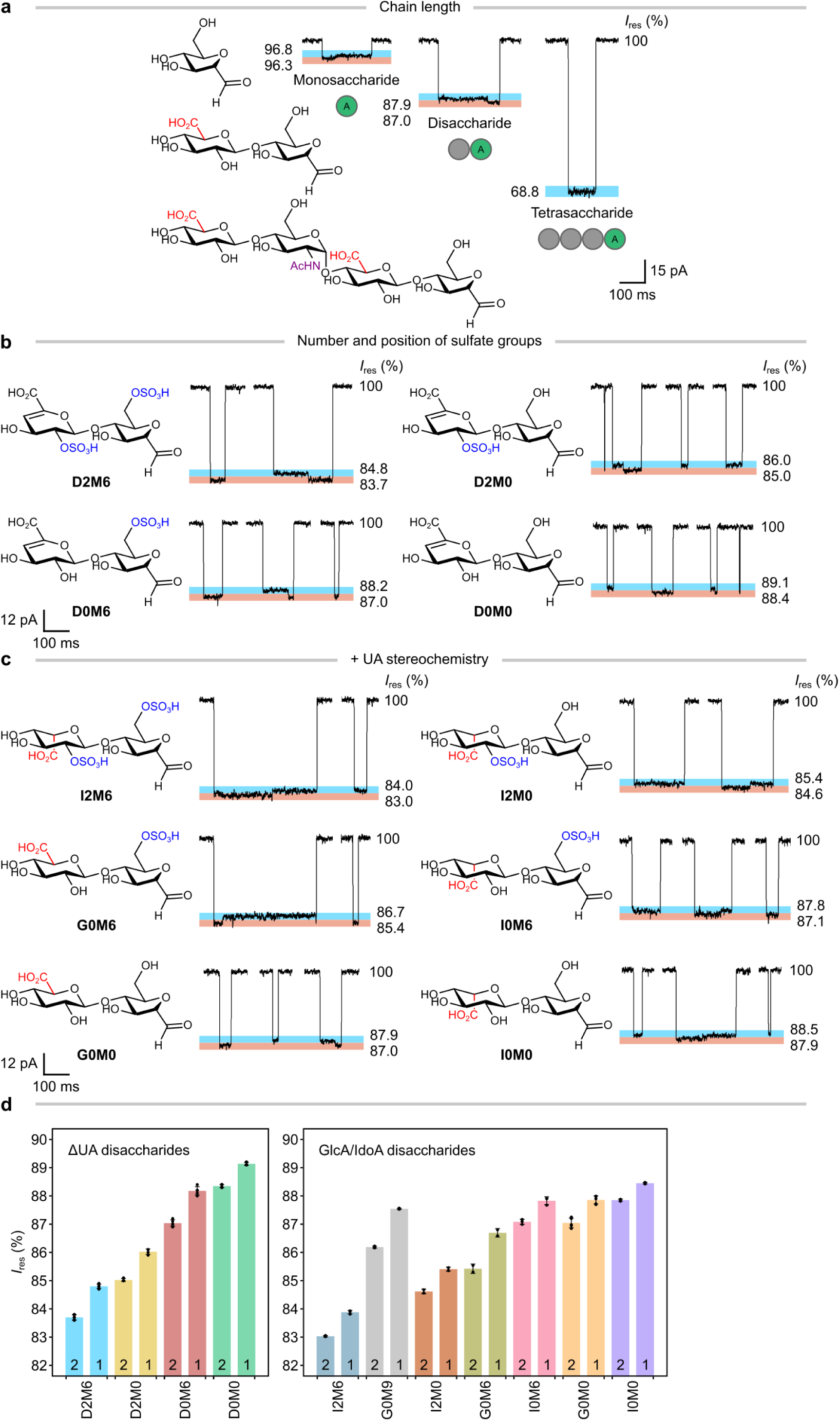
Resolution of GAG chain length and fine structural features using the αHL-Cys115 pore. **a,** Mono-, di-, and tetrasaccharides (M0, G0M0, and G0A0G0M0, respectively) yielded distinct *I*_res_ values, with each additional sugar residue contributing a ∼9 % step decrease; notably, only a single level was observed for G0A0G0M0. AMan and non-AMan residues are depicted as green and grey circles, respectively. **b–c,** ΔUA disaccharides with various sulfation states (**b**) and UA disaccharides with various sulfation and epimerization states (**c**) were resolved based on differences in the *I*_res_ values of interconverting levels, highlighted in blue and pink (e.g., 1.5–3 % step difference per sulfate and ∼1 % step decrease from IdoA to GlcA). **d,** The bar plots show the mean *I*_res_ levels of 11 AMan disaccharides recorded across *N* ≥ 3 pores, except for I2M0 and G0M6 (*N* = 2) due to limited material (also see Table S6). Points indicate *I*_res_ levels from independent pores; vertical error bars indicate s.d. The bars are numbered either ‘1’ or ‘2’; for each sugar with paired interconverting levels, ‘1’ indicates the level with the higher *I*_res_ value. **Recording conditions:** 4 M LiCl, 20 mM HEPBS, 40 μM EDTA, titrated to pH 8 using KOH; sugars (*cis*); +150 mV (*trans*); 23.8 ± 1 °C. Sugar concentrations ranged from 0.19 mM to 0.84 mM, corresponding to 42–139 µg of material. All *I*_res_ values are reported to one decimal place.

**Extended Data Fig. 6.**
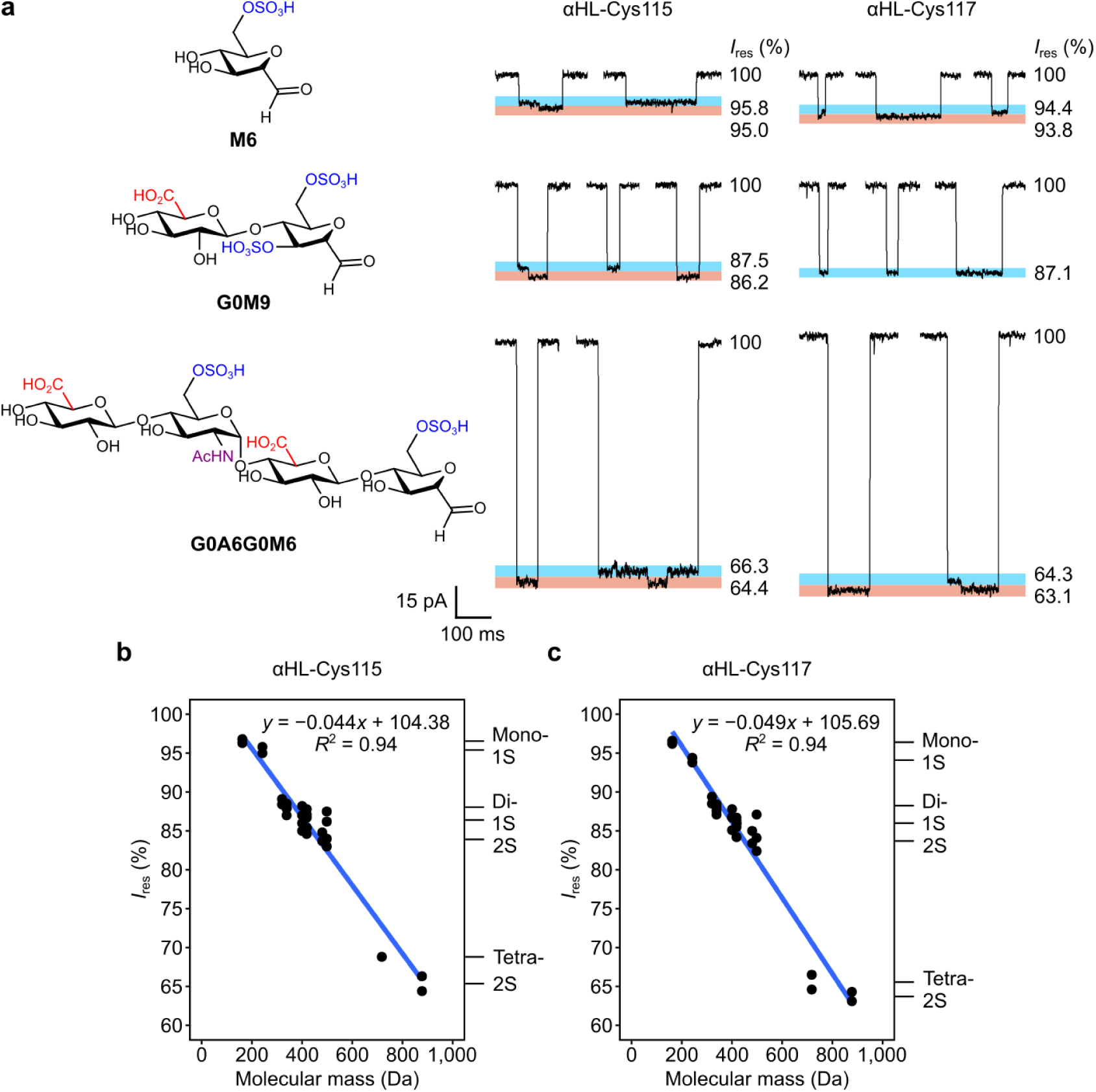
*I*_res_ values correlate with sugar size and sulfation. **a,** Representative events (levels highlighted in blue and pink) for a different set of mono-, di-, and tetrasaccharides (M6, G0M9, and G0A6G0M6, respectively; also see Tables S5–6). *N* = 3 pores were recorded for each sugar, except for αHL-Cys117/G0A6G0M6 (*N* = 2) due to limited material. Notably, only one *I*_res_ level was observed for αHL-Cys117/G0M9. Sugar concentrations ranged from 0.15 mM to 0.34 mM, corresponding to 44–90 µg of material. All *I*_res_ values are reported to one decimal place. **b–c,** The mean *I*_res_ levels of 15 AMan standards (comprising two monosaccharides, 11 disaccharides, and two tetrasaccharides), recorded using independent αHL-Cys115 (**b**) and αHL-Cys117 pores (**c**), are plotted as points against sugar molecular mass. The NSOSHp-derived disaccharide G5M9 is omitted. Blue lines indicate fits from ordinary least squares linear regression; vertical error bars (not visible at the present scale) indicate s.d. The markers of chain length and sulfation are approximate. *R*^2^, coefficient of determination. **Recording conditions:** 4 M LiCl, 20 mM HEPBS, 40 μM EDTA, titrated to pH 8 using KOH; sugars (*cis*); +150 mV (*trans*); 23.8 ± 1 °C.

**Extended Data Fig. 7.**
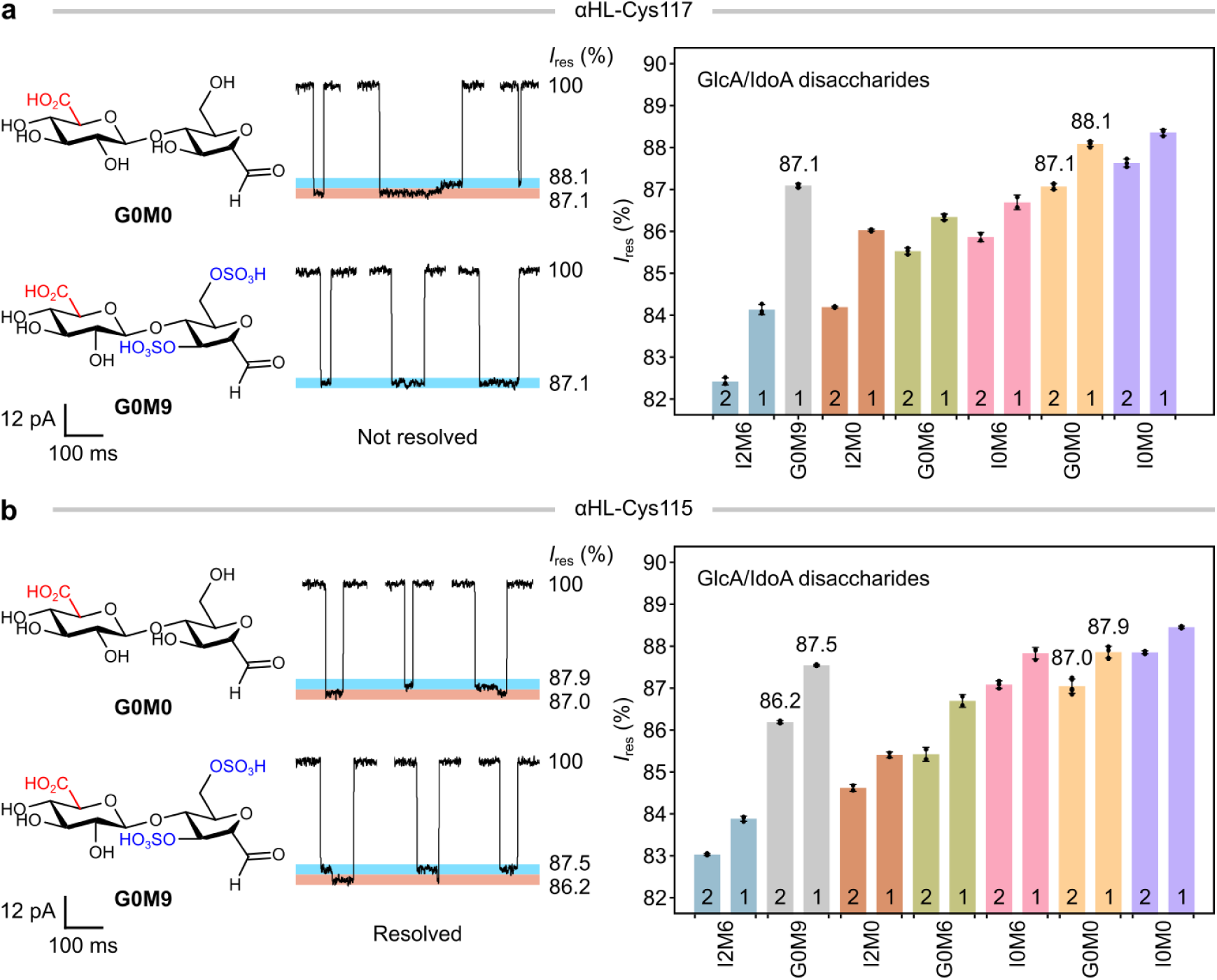
Complementary analytical performance of the αHL-Cys117 and αHL-Cys115 pores. **a–b,** A direct comparison of disaccharide resolving power between the αHL-Cys117 (**a**) and αHL-Cys115 (**b**) pores. This complementarity is exemplified in the resolution of G0M0 and G0M9. In the αHL-Cys117 pore, the single *I*_res_ level of G0M9 completely overlaps with a G0M0 level at 87.1 %; by comparison, in the αHL-Cys115 pore, the disaccharide pair together yields four distinct and fully resolved levels. The bar plots show the mean *I*_res_ levels of seven saturated UA disaccharides recorded across *N* ≥ 3 pores, except for αHL-Cys117/I0M6, αHL-Cys115/I2M0, and αHL-Cys115/G0M6 (*N* = 2). Points indicate *I*_res_ levels from independent pores; vertical error bars indicate s.d. The bars are numbered either ‘1’ or ‘2’; for each sugar with paired interconverting levels, ‘1’ indicates the level with the higher *I*_res_ value. **Recording conditions:** 4 M LiCl, 20 mM HEPBS, 40 μM EDTA, titrated to pH 8 using KOH; sugars (*cis*); +150 mV (*trans*); 23.8 ± 1 °C. All *I*_res_ values are reported to one decimal place.

**Extended Data Fig. 8.**
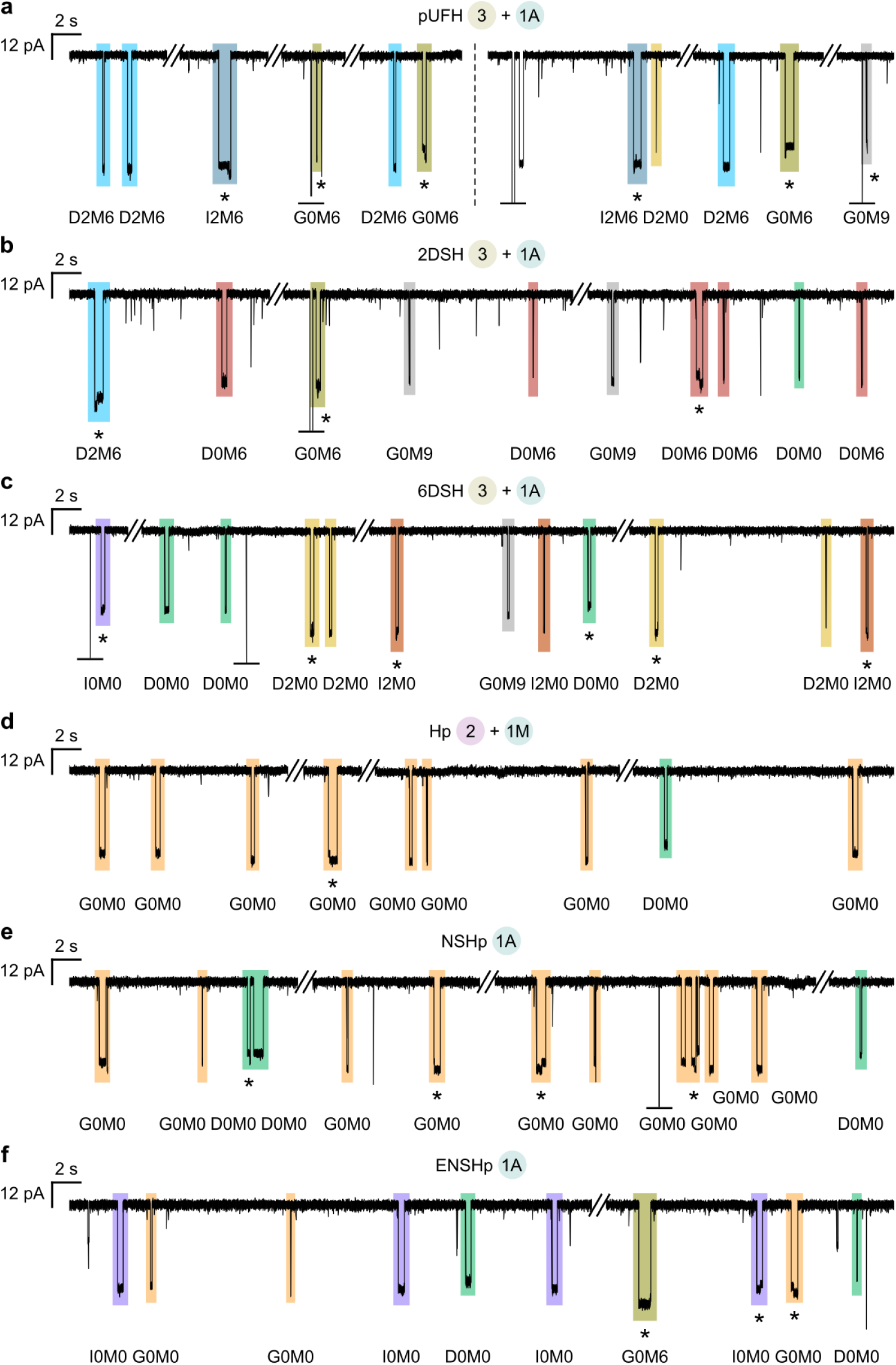
Heparin and heparosan fingerprints reveal unanticipated low-abundance components. **a–c,** The pUFH (**a**), 2DSH (**b**), and 6DSH (**c**) fingerprints generated using the αHL-Cys115 pore under conditions **3 + 1A** (Fig. 3) each exhibited events distinct from D2M6, D2M0, D0M6, and D0M0 (see Table S8 for percentage of events assigned); a subset of these was identified as saturated UA disaccharides. The displayed pUFH-derived events are extracted from separate traces. **d–f,** The Hp (**d**), NSHp (**e**), and ENSHp (**f**) fingerprints generated using the αHL-Cys117 pore under the differential application of conditions **1A**, **1M**, and **2** (Fig. 4) each contained a small number of events distinct from G0M0 and I0M0 (see Table S9 for percentage of events assigned), some of which were identified as D0M0, alongside low levels of G0M6 identified within the ENSHp fingerprint. Interconversion events are marked with an asterisk. Graphically truncated events are marked with a horizontal bar. **Recording conditions:** 4 M LiCl, 20 mM HEPBS, 40 μM EDTA, titrated to pH 8 using KOH; sugars (*cis*); +150 mV (*trans*); 23.8 ± 1 °C. Events with lifetimes under 2 ms were discarded (Methods).

**Extended Data Fig. 9.**
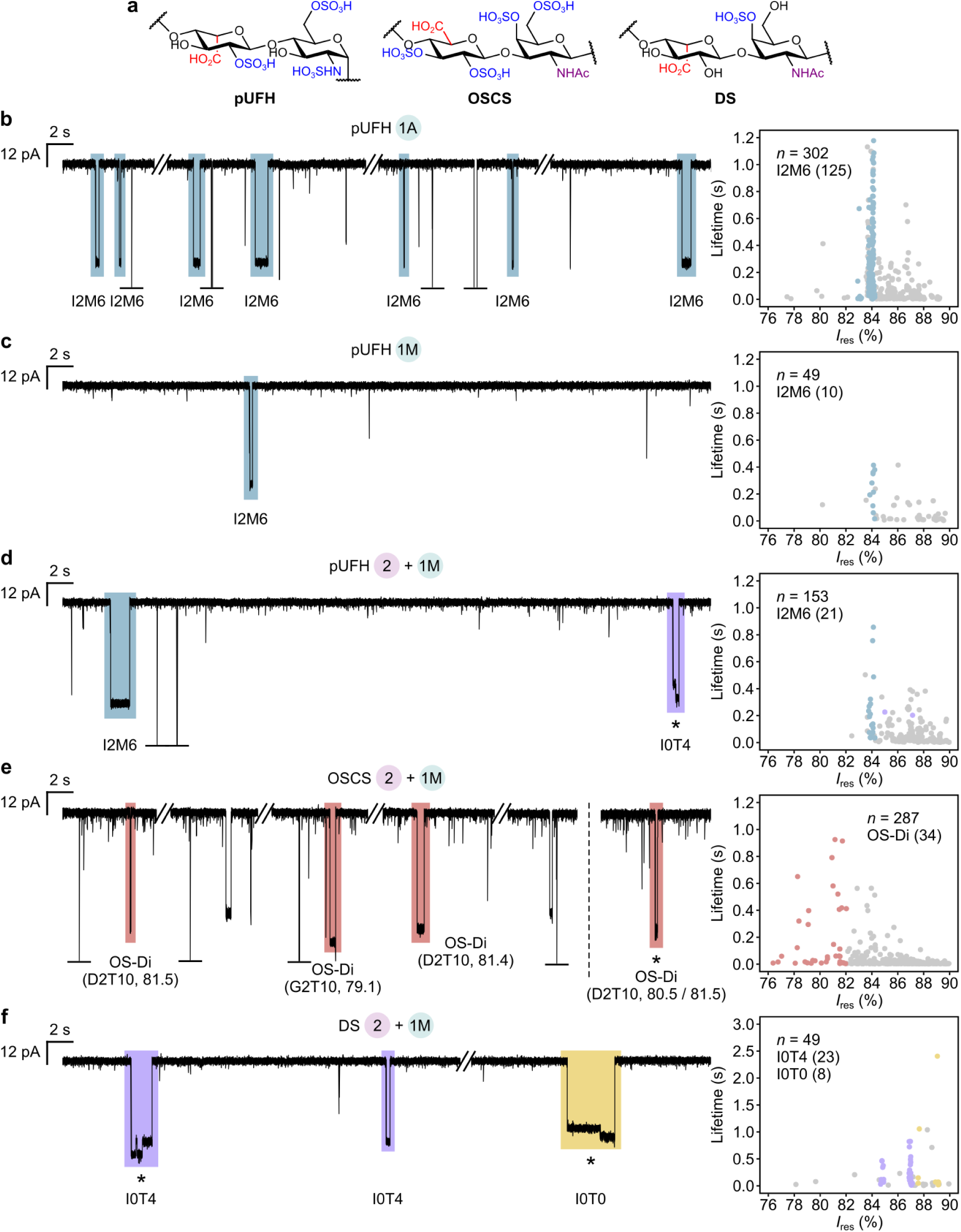
Nanopore fingerprints of pUFH, OSCS, and DS. **a,** The structures of pUFH, OSCS, and DS (representative). **b–f,** The nanopore fingerprints of pUFH (**b–d**), OSCS (**e**), and DS (**f**) revealed various characteristic anhydrosugars. Conditions **1A**, **1M**, and **2** were differentially applied to these polysaccharides; the resulting product mixtures (∼0.3–0.7 mg) were probed by the αHL-Cys115 pore (Supplementary Methods). Representative current traces are shown, along with assignments for I2M6 (to guide the eye; see Supplementary Methods for assignment criteria). The *I*_res_ values of OSCS-derived sugars in the *I*_res_ 76–82 % cluster (OS-Di) are reported to one decimal place. D2T10 and G2T10 are tentatively assigned (see Extended Data Fig. 10b for sugar structures and *I*_res_ predictions). The displayed interconverting D2T10 event is extracted from a separate trace. I0T4 and I0T0 in the DS fingerprint are also tentatively assigned (see Extended Data Fig. 10c for sugar structures and *I*_res_ predictions). Interconversion events are marked with an asterisk. Graphically truncated events are marked with a horizontal bar. The scatter plot in **b** shows event levels from a single pore. The scatter plots in **c–f** show combined event levels from three, three, four, and three pores, respectively. Event levels in the scatter plots are colour-coded as in the traces, and levels from all other sugars are depicted in grey. Within the scatter plots, event counts for the specified sugars are given in parentheses. *n*, total number of events. **Recording conditions:** 4 M LiCl, 20 mM HEPBS, 40 μM EDTA, titrated to pH 8 using KOH; sugars (*cis*); +150 mV (*trans*); 23.8 ± 1 °C. Events with lifetimes under 2 ms were discarded (Methods).

**Extended Data Fig. 10.**
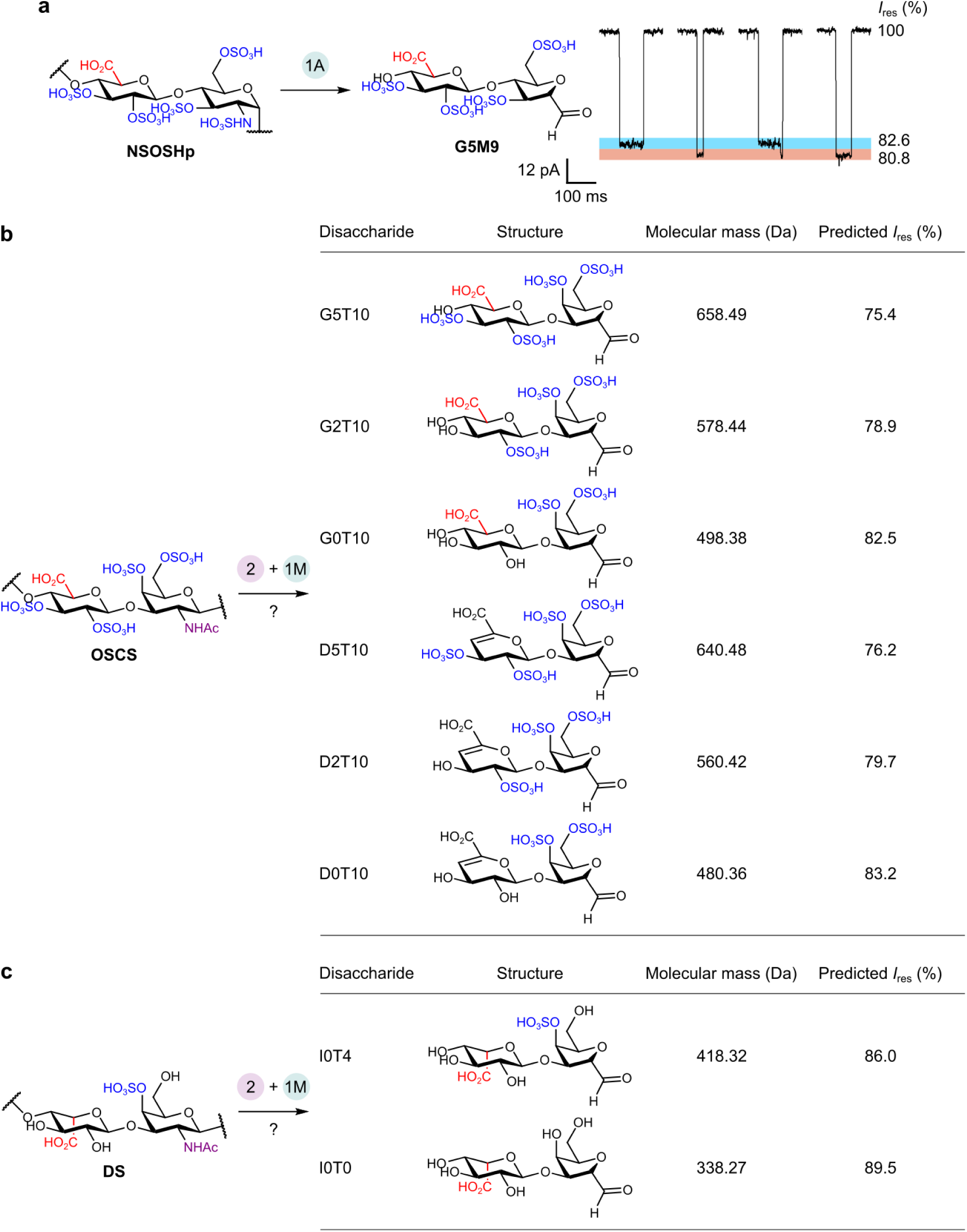
*I*_res_ data for G5M9 and predictions for potential OSCS- and DS-derived sugars. **a,** Representative events (levels highlighted in blue and pink) for the tetrasulfated AMan disaccharide G5M9, generated from the ring contraction of NSOSHp at pH 1.5 (condition **1A**) and probed using the αHL-Cys115 pore. *I*_res_ values are reported to one decimal place and represent means from *N* = 3 pores (Table S6). **Recording conditions:** 4 M LiCl, 20 mM HEPBS, 40 μM EDTA, titrated to pH 8 using KOH; sugars (*cis*); +150 mV (*trans*); 23.8 ± 1 °C. **b–c,** *I*_res_ values for potential OSCS- (**b**) and DS-derived sugars (**c**) were predicted using the linear fit reported in Extended Data Fig. 6b. The inclusion of ΔUA disaccharides in **b** was prompted by NMR analysis of hydrazinolysed OSCS, which suggested partial conversion of UA into ΔUA concomitant with polymer cleavage (Fig. S18). The structure of DS depicted in **c** is representative.

**Extended Data Fig. 11.**
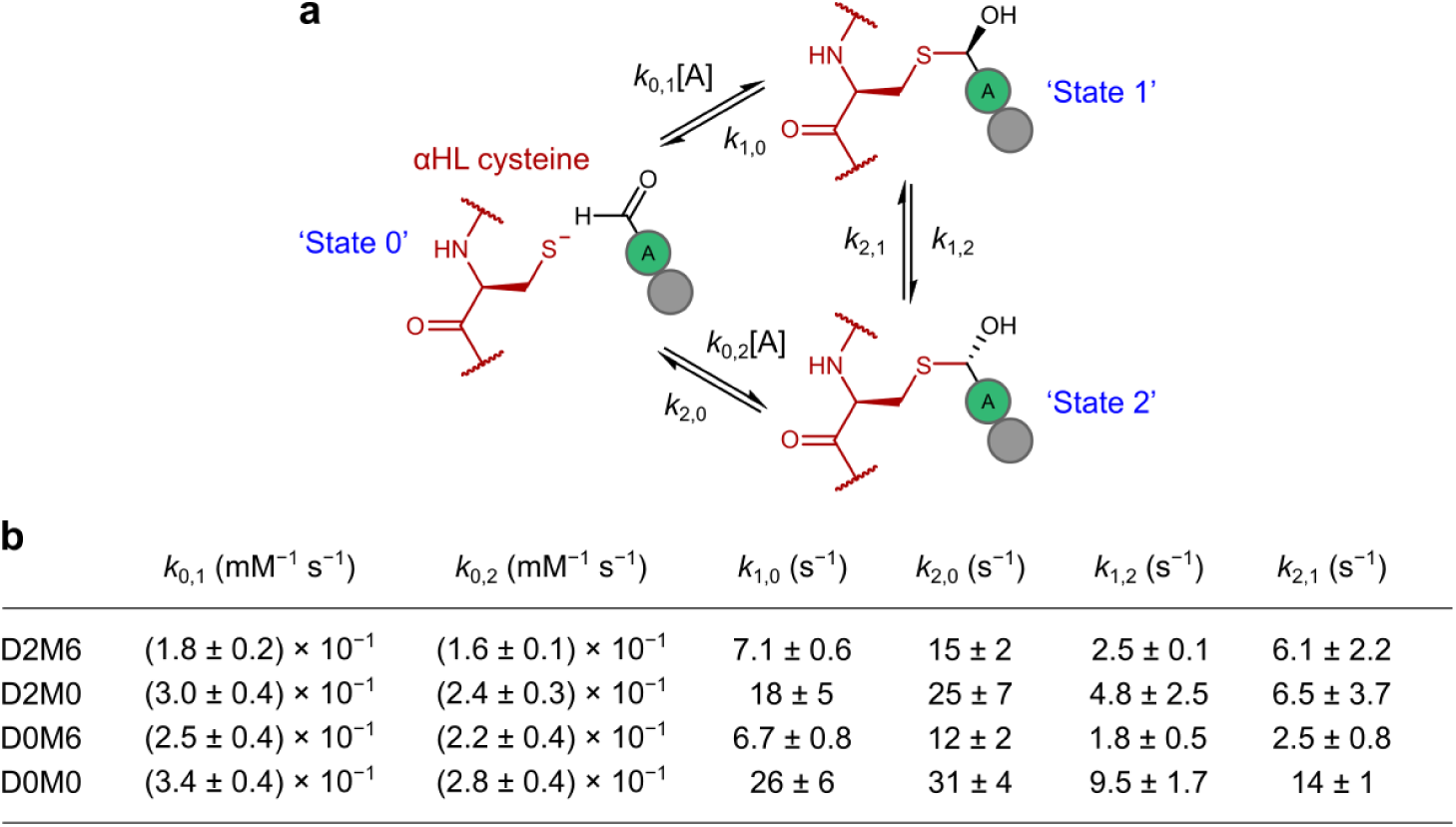
Kinetics of disaccharide sensing. **a,** Thiohemiacetal formation, dissociation, and stereogenic inversion (where ‘A’ = anhydrosugar). AMan and non-AMan residues are depicted as green and grey circles, respectively. **b,** Rate constants (mean ± s.d. across *N* ≥ 3 pores) for the reactions between the αHL-Cys115 pore and each of four ΔUA disaccharides, extracted as detailed in Supplementary Methods. For each disaccharide, state 1 corresponds to the event level with the higher *I*_res_ value.

**Extended Data Table 1.** Comparison of methods for the identification of contaminants in heparin. References for each method are given within the main text. A MinION flow cell features 512 simultaneously active pores connected to a single 0.1 mL analyte reservoir^35^, representing one-fifth the volume of the individual compartments used in our experimental setup (Methods). MM, multivariate modelling; 2D, two-dimensional; LOD, limit of detection.

|  | <sup>1</sup>H NMR<br>(USP) | <sup>1</sup>H NMR<br>+ MM | 2D NMR | LC-MS | IR | Nanopores<br>(this work) |
| --- | --- | --- | --- | --- | --- | --- |
| Contaminant<br>scope | OSCS +<br>some NAc<br>sugars | General | General | General | General | General |
| Structural<br>information | Minimal | Yes | Yes | Yes | Minimal | Yes |
| Sample quantity | > 10 mg | 20 mg | 10–25 mg | ~10 µg | 1–10 mg | ~0.3–0.7 mg<br>per fingerprint<sup>[a]</sup> |
| Contaminant<br>LOD<sup>[b]</sup> | 0.25 %<br>(500 MHz) | 0.25–10 %<br>(600 MHz) | 2–4 %<br>(500 MHz) | 0.1 % | 0.5–5 % | Only ~20 %<br>tested |
| Sample preparation<br>and analysis time | Hours for<br>exchange<br>into D<sub>2</sub>O | Hours for<br>exchange<br>into D<sub>2</sub>O | Hours for<br>exchange<br>into D<sub>2</sub>O | ~3.5 h | ~70 s | ~1.5–5.5 h <sup>[a] [c]</sup> |
| Instrument size | Large | Large | Large | Large | Small<br>(benchtop) | Smallest<br>(pocket size)<sup>[d]</sup> |
| Instrument cost<sup>[e]</sup> | High<br>(> \$500k) | High<br>(> \$500k) | High<br>(> \$500k) | High<br>(> \$500k) | Moderate<br>(> \$15k) | Low<br>(< \$5k)<sup>[d]</sup> |
| Operator expertise | Minimal | High | High | Moderate | Moderate | Minimal |

<sup>[a]</sup> The MinION may reduce these sample quantities and sensing times by a combined ~2,500-fold
<sup>[b]</sup> Concentrations are (w/w)
<sup>[c]</sup> Reactions **1A** and **1M** are complete within 10 min, whereas reaction **2** is carried out for 4 h
<sup>[d]</sup> Adaptation to the MinION
<sup>[e]</sup> Prices are in US dollars (\$) as of May 2026

