## Supplementary Information for "Nanopore-based sequence deconvolution of diverse glycosaminoglycans"

**Supplementary Discussion 1**

To date, 27 unique disaccharide units have been identified as products of HS biosynthesis (Table S1). MS, in its simplest form, can resolve at most 11 of these disaccharides because, in their physiological protonation states, they yield only 11 unique molecular masses. By extension, MS can resolve a maximum of 11 out of the 18 ΔUA disaccharides generated via eliminative digestion (Table S2), and only four out of the 15 AMan disaccharides produced via Tiffeneau–Demjanov ring contraction (Table S3).

More advanced MS configurations offer only moderate improvements in distinguishing HS disaccharides. While MS/MS, through ion activation and subsequent diagnostic fragmentation, can pinpoint the positions of sulfate groups in GAG di- and oligosaccharides, it often struggles to determine UA stereochemistry[1]. Although it has long been appreciated that UA stereochemistry can influence fragmentation pathways[2,3], this effect is weak and difficult to capture in practice[4–6]. Consequently, in the absence of orthogonal validation, UA stereochemistry is typically either omitted entirely or assigned based on prescriptive biosynthetic assumptions that may or may not hold true[7,8]. Even in rare instances of successful MS/MS-only assignment, highly specific constraints emerge, as seen when multivariate modelling elucidated 15 unique HS disaccharide units present within a synthetic tetrasaccharide library[9]. Central to this approach was the use of 'symmetry-breaking' reducing-end labels, which enabled fragment ions originating from the two termini of each tetrasaccharide to be clearly distinguished. However, because only reducing-end UA residues could be reliably assigned, the generality of this approach may be limited.

Coupling MS with liquid-phase fractionation can provide orthogonal dimensions for characterizing regio- and stereoisomeric GAGs[10] while simultaneously reducing matrix effects during ionization[11], yet numerous practical challenges remain. Widely used fractionation setups (e.g., C18 reversed-phase HPLC) struggle immensely with the extreme polarity of many GAGs[1]. While more specialized approaches—including reversed-phase ion-pairing (RPIP)[12,13], hydrophilic interaction (HILIC)[5,14–16], and porous graphitic carbon (PGC)[17,18] chromatographies as well as capillary electrophoresis[6]—have achieved greater success, co-elution, non-elution, variation in retention times, and analyte peak splitting (e.g., due to the presence of α and β anomers) remain commonplace. A particular HILIC-MS platform has demonstrated a rare, impressive breadth of resolution, quantifying eight HS ΔUA disaccharides and eight AMan disaccharides within their respective mixtures when surveyed in parallel[14]. Even so, this analytical capability seems to have approached a physical ceiling: a ninth and tenth ΔUA disaccharide (ΔUA-GlcN6S and ΔUA2S-GlcN6S), which were isomeric with some of those previously tested (ΔUA-GlcNS and ΔUA2S-GlcNS, respectively), entirely eluded chromatographic and MS resolution.

In contrast to standard MS, which solely measures mass-to-charge ratios, ion mobility MS also measures size- and shape-dependent gas-phase mobilities and is therefore capable of resolving regio- and stereoisomeric GAGs[1,10]. However, once again, achieving this in practice can be challenging, in no small part due to the relatively low resolving power of current instrumentation along the ion mobility axis—a widely recognized limitation of the technique[4,16,19–22]. Currently, a difference in collision cross-section (CCS) values of at least ~0.5–1 % is necessary for the reliable differentiation of isomers, a threshold that may not always be met; for context, the isomeric HS disaccharides ΔUA-GlcNS6S and ΔUA2S-GlcNS exhibit a CCS difference of only 0.8 $Å^2$ or ~0.6 % (both in 1− charge states, measured in helium buffer gas)[21]. In a recent state-of-the-art example, trapped ion mobility spectrometry was used to resolve and quantify 12 HS ΔUA disaccharides within mixtures[22]. Notably, most disaccharides in their unmodified reducing forms were poorly distinguished due to the presence of multiple gas-phase conformers and anomers for each species, resulting in complex mobilograms with overlapping peaks. Although adequate resolution of the 12 disaccharides was achieved via a two-step chemical derivatization procedure, resorting to such protocols—which can yield incomplete reactions and product distributions[23,24]—can introduce additional experimental error and hamper throughput.

Gas-phase IR spectroscopy, which integrates MS and IR action spectroscopy, is capable of distinguishing regio- and stereoisomeric GAGs by their vibrational modes[1,10]. This capability, however, can be constrained by spectral congestion, a major source of which is ion heating upon irradiation; indeed, initial approaches employing multiphoton irradiation at ambient temperatures yielded well-resolved and diagnostic spectral features only for a few simple GAG mono- and disaccharides[25,26]. More sophisticated cryogenic approaches—exemplified by helium nanodroplet IR spectroscopy—have enabled the differentiation of a broader array of more complex structures, with several leading examples in HS characterization having resolved a cumulative total of eight ΔUA disaccharides, six tetrasaccharides, a pentasaccharide, and a hexasaccharide[27–29]. Although potentially powerful, these approaches currently depend on highly specialized light sources and ion traps. This dependence has, until only recently[30], confined their use to free-electron laser facilities and home-built experimental setups in a select few laboratories worldwide.

**Supplementary Discussion 2**

Nanopores are broadly classified into solid-state and protein variants[31]. Solid-state pores exhibit exceptional mechanical and chemical stability; however, fabrication constraints typically lead to pores with physical dimensions that exceed those of single biomolecules manyfold, which can compromise structural resolution[31,32] (see below). By contrast, although protein pores require a physical support more prone to disruption (e.g., a lipid bilayer), they are generally better size-matched with single biomolecules and therefore offer higher resolution[31,33].

Solid-state nanopores were initially used to sense GAGs of different subclasses and chain lengths (~20-mers to polysaccharides)[34–37], and subsequently, GAGs within the same subclass and length regime (HS ~80-mers) but bearing broad differences in sulfation and epimerization[38]. In all instances, structurally heterogeneous samples were used; their translocation via unassisted electrophoresis produced signals with overlapping amplitudes and predominantly sub-millisecond lifetimes that precluded the resolution of individual sugar residues. Similar results were obtained when these experiments were repeated using protein pores[39–41], despite their better size match; this outcome persisted even when structurally homogeneous HS 4–12-mer standards were employed[42].

In a recent attempt to map sulfation within GAG chains, a library of structure-defined HS 9- and 18-mer standards in various sulfation states was ligated at both termini to DNA handles, then translocated through protein nanopores via DNA-directed motor proteins[43]. This approach captured individual HS translocation events spanning up to hundreds of milliseconds, which a transformer model assigned to the standards with moderately high accuracy. While this suggested a genuine, rather than random, variation in the resulting event amplitudes according to the underlying sulfation pattern, these amplitudes remained too stochastically distributed and overlapping to indicate either the presence or location of individual sulfate groups—this was exemplified by the high, symmetric misassignment rates of standards exhibiting progressive 6-*O*-sulfation across the same backbone. These results illustrate the critical distinction between macroscopic classification and structural resolution; despite numerous attempts, the latter remains an entirely uncharted frontier.

### Supplementary Discussion 3

Previously, this benchmark was held by an online HILIC-MS platform capable of identifying eight HS ΔUA disaccharides and eight AMan disaccharides[14] (see Supplementary Discussion 1 for details). By applying our designed deconvolutive strategy (see main text), these eight AMan variants can cover 18 biosynthetic HS disaccharide units; consequently, the total number of these units and their eliminative digestion products tractable by this MS-based platform is 18 + 8 = 26.

### Supplementary Note 1

$I_{res}$ data acquired from single pores were generally well modelled by Gaussian distributions with HWHM values of ~0.1 % (Figs S5–6 and Tables S5–6). Considering only single-level (non-interconverting) events observed within a given pore, the normalized area of each Gaussian distribution extending beyond its intercept with a neighbouring distribution represents the one-tailed probability of event misassignment under equal sampling counts[44]. For a pair of idealized distributions with centroids displaced by 0.2 % and identical HWHM values of 0.1 %, this individual misassignment probability is only 12.0 % (corresponding to a total shared error of 23.9 %); in other words, the vast majority (88.0 %) of events are correctly assigned. Importantly, the proportion of correctly assigned events increases when also considering multi-level (interconverting) events, as the temporal pairing of two distinct levels forces a stricter criterion for assignment. Using this framework as a point of reference, and accounting for some small pore-to-pore centroid variation (Tables S5–6), we selected an $I_{res}$ separation of 0.2 % as a threshold for resolving individual events with satisfactory confidence. Hence, the αHL-Cys117 pore sufficed to resolve:

- all pairs of disaccharide standards except the G0M0/G0M9 pair (level 2 of G0M0 and level 1 of G0M9 are identical at $I_{res}$ 87.1 %); and
- any given disaccharide from the five saturated UA variants I2M6, G0M6, I0M6, G0M0, and I0M0, with a minimum $I_{res}$ separation of 0.3 % (occurring between level 1 of G0M0 at 88.1 % and level 1 of I0M0 at 88.4 %).

Conversely, the αHL-Cys115 pore sufficed to resolve:

- any given disaccharide from the four ΔUA variants D2M6, D2M0, D0M6, and D0M0, with a minimum $I_{res}$ separation of 0.2 % (occurring between level 1 of D2M6 at 84.8 % and level 2 of D2M0 at 85.0 %, and between level 1 of D0M6 at 88.2 % and level 2 of D0M0 at 88.4 %); and
- any given disaccharide from the six saturated UA variants I2M6, I2M0, G0M6, G0M0, I0M0, and G0M9 (despite complete overlap between level 1 of I2M0 and level 2 of G0M6 at $I_{res}$ 85.4 %, and between level 1 of G0M0 and level 2 of I0M0 at $I_{res}$ 87.9 %, all these disaccharides were straightforwardly resolved by their remaining levels).

Thus, in combination, the two pores sufficed to resolve any given disaccharide across our entire 11-member panel. Moreover, the αHL-Cys115 pore is capable of fully resolving, from this core panel, the NSOSHp-derived non-natural disaccharide G5M9.

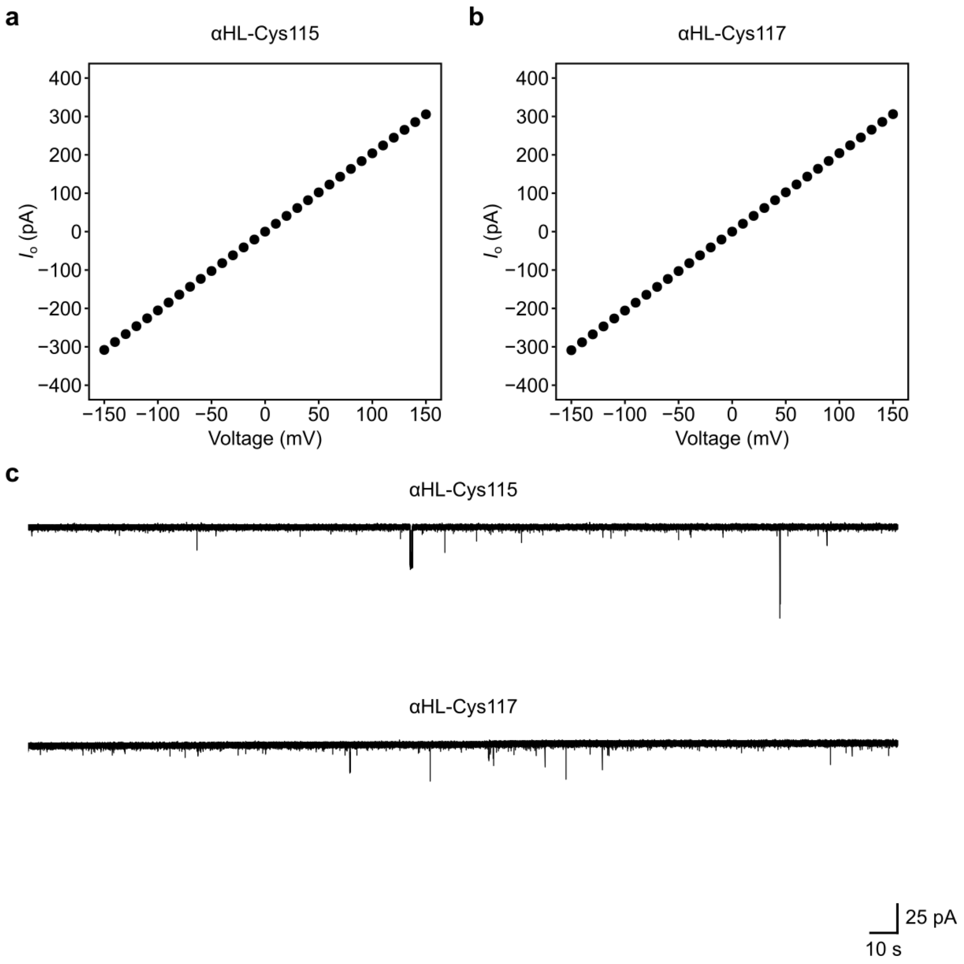

**Fig. S1 | Open-pore characteristics of the αHL-Cys115 and αHL-Cys117 pores. a–b,** The open-pore currents ($I_o$) of the αHL-Cys115 (**a**) and αHL-Cys117 (**b**) pores are plotted against voltage. Points indicate the mean $I_o$ from $N$ = 10 pores; vertical error bars (not visible at the present scale) indicate s.d. At +150 mV, the mean $I_o$ is 305.6 ± 2.8 pA and 306.1 ± 5.0 pA for αHL-Cys115 and αHL-Cys117, respectively. **c,** Representative current traces of each pore variant recorded over a span of 5 min. Both variants produce stable open-pore baseline currents, with rare gating events and microsecond-long current blockades ('spikes') immediately distinguishable from covalent events. **Recording conditions:** 4 M LiCl, 20 mM HEPBS, 40 μM EDTA, titrated to pH 8 using KOH; +150 mV (*trans*); 23.8 ± 1 °C.

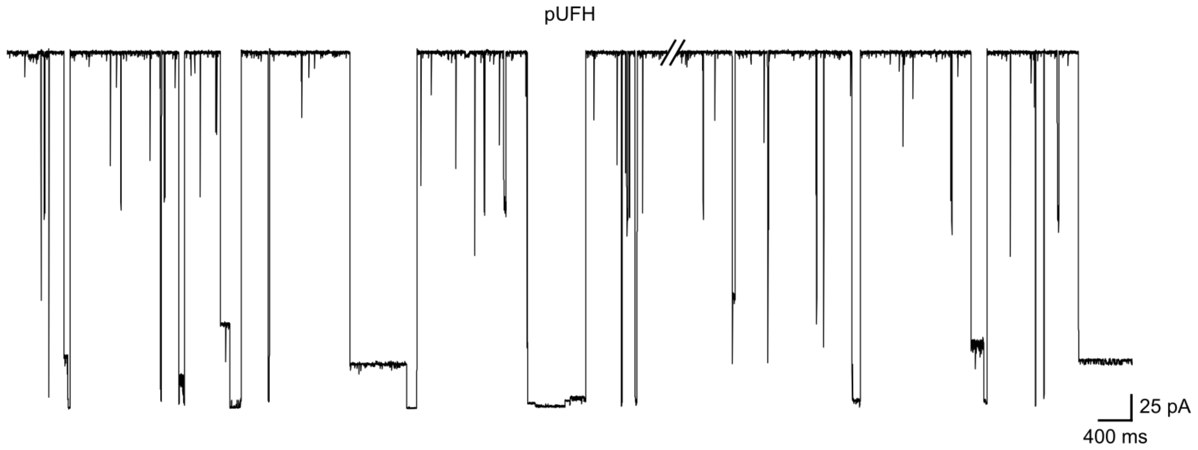

**Fig. S2 | Lack of covalent events with non-ring-contracted polysaccharidic heparin.** pUFH (150 µg) was introduced to the αHL-Cys117 pore. The observed events were predominantly 'spikes' with sub-millisecond lifetimes or longer-lived blockades with highly variable amplitudes (consistent with early glycan sensing approaches[39,45]); no clear covalent events emerged. **Recording conditions:** 4 M LiCl, 20 mM HEPBS, 40 µM EDTA, titrated to pH 8 using KOH; sugars (*cis*); +150 mV (*trans*); 23.8 ± 1 °C.

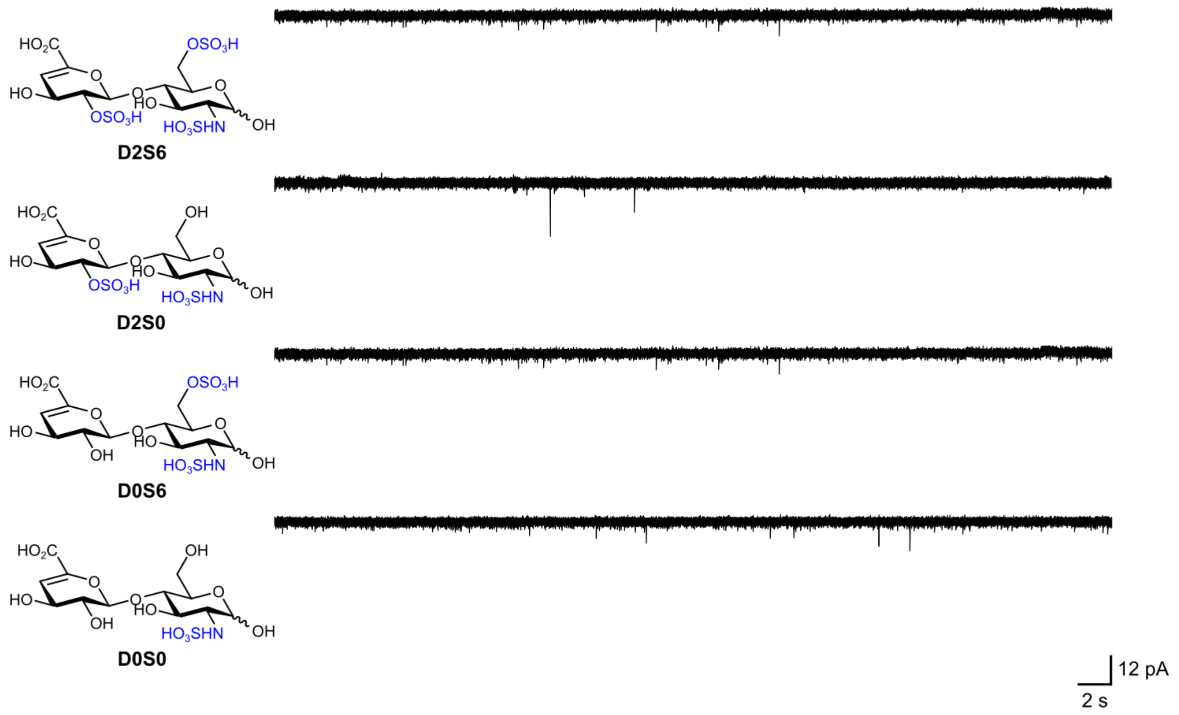

**Fig. S3 | Lack of covalent events with sugars bearing free anomeric centres.** No covalent events emerged following the separate introduction of ΔUA2S-GlcNS6S (D2S6), ΔUA2S-GlcNS (D2S0), ΔUA-GlcNS6S (D0S6), and ΔUA-GlcNS (D0S0) to the αHL-Cys117 pore. Sugar concentrations were 0.54 mM, 0.68 mM, 0.68 mM, and 0.84 mM, respectively. In all instances, the subsequent addition of AMan standards yielded covalent events, thereby confirming pore viability (data not shown). **Recording conditions:** 4 M LiCl, 20 mM HEPBS, 40 µM EDTA, titrated to pH 8 using KOH; sugars (*cis*); +150 mV (*trans*); 23.8 ± 1 °C.

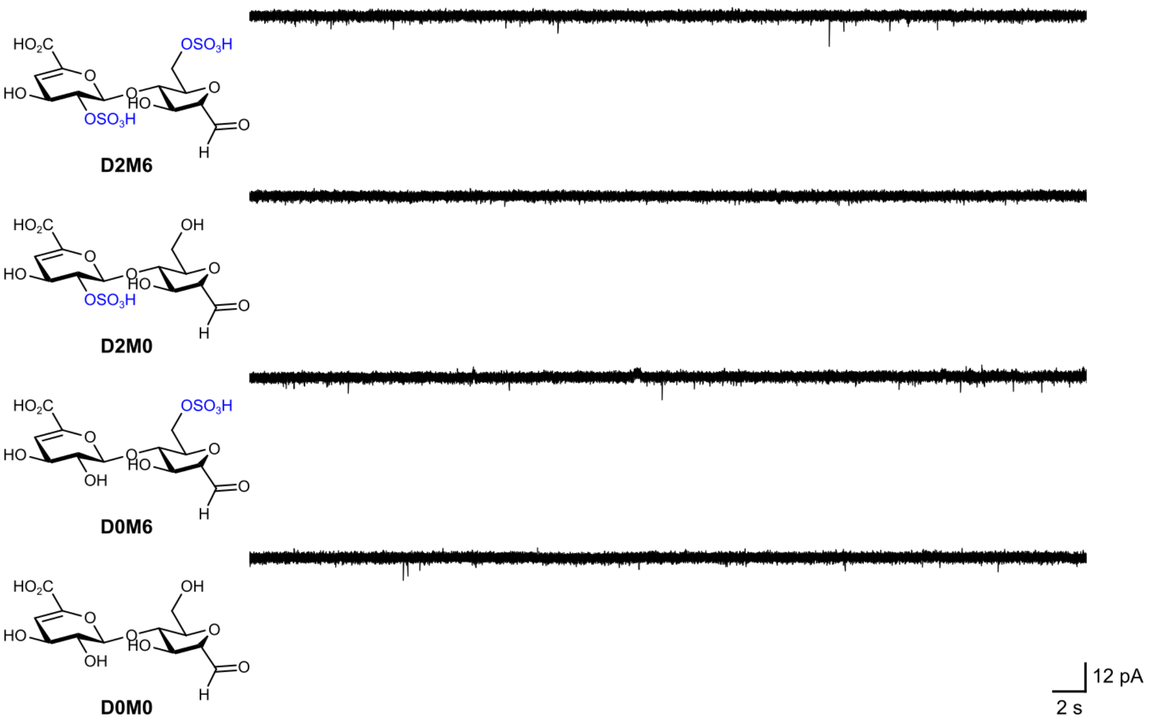

**Fig. S4 | Lack of covalent events with a cysteine-free pore variant.** No covalent events emerged following the separate introduction of D2M6, D2M0, D0M6, and D0M0 to the cysteine-free αHL-$A_7$ pore (Methods). Sugar concentrations were 0.64 mM, 0.57 mM, 0.76 mM, and 0.84 mM, respectively. **Recording conditions:** 4 M LiCl, 20 mM HEPBS, 40 μM EDTA, titrated to pH 8 using KOH; sugars (*cis*); +150 mV (*trans*); 23.8 ± 1 °C.

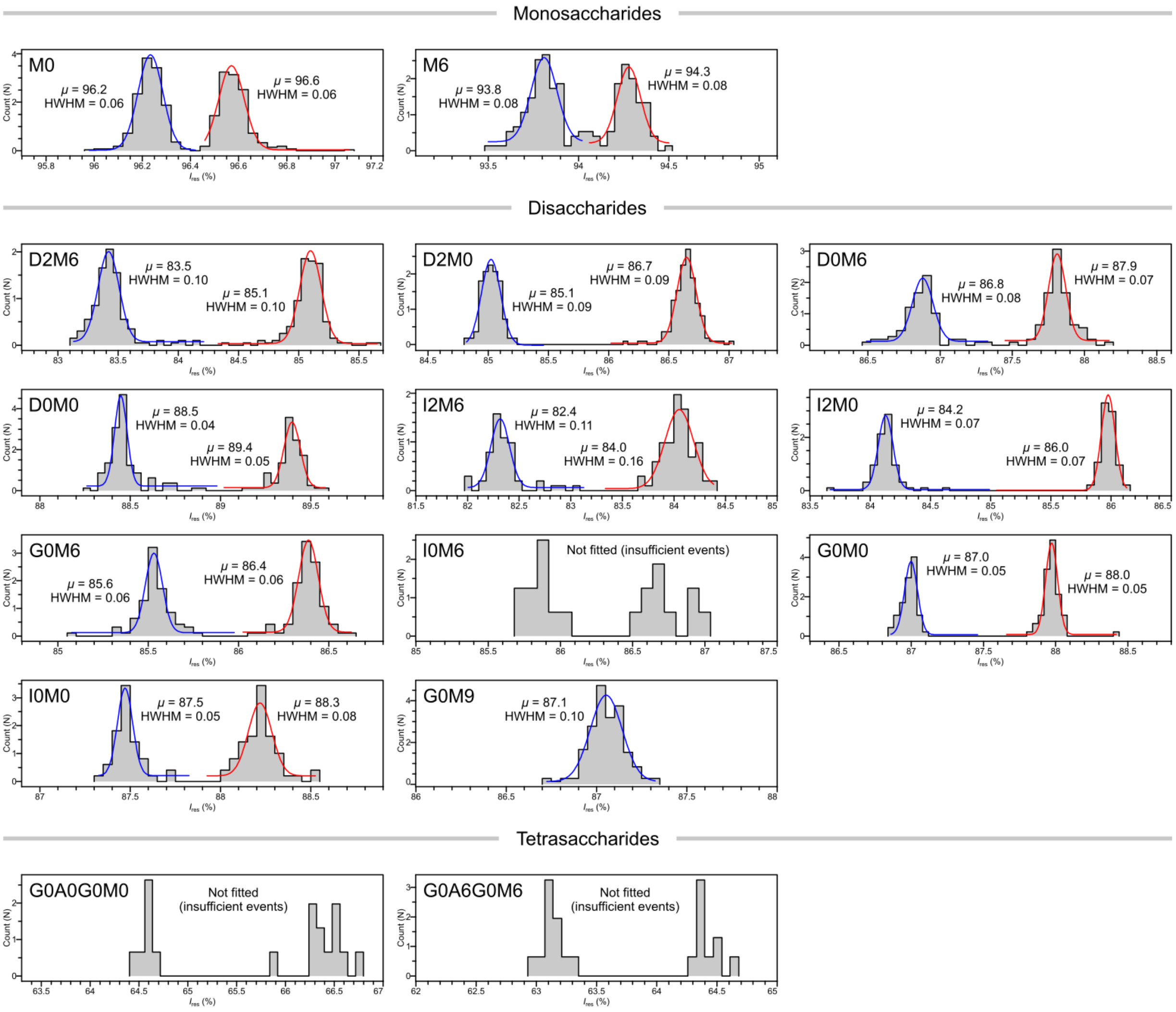

**Fig. S5 | $I_{res}$ levels are measured with high precision (αHL-Cys117).** Representative $I_{res}$ data for each AMan standard acquired from single pores are overlaid with single-term Gaussian fits (one for each level or 'half-doublet') computed using the Clampfit Fit function with bin widths of 0.04–0.08 %, where all other parameters were set to their default values. Gaussian s.d. values were multiplied by 1.1774 to yield HWHM values. $\mu$, centroid. $\mu$ and HWHM values are reported to one and two decimal places, respectively (also see Table S5).

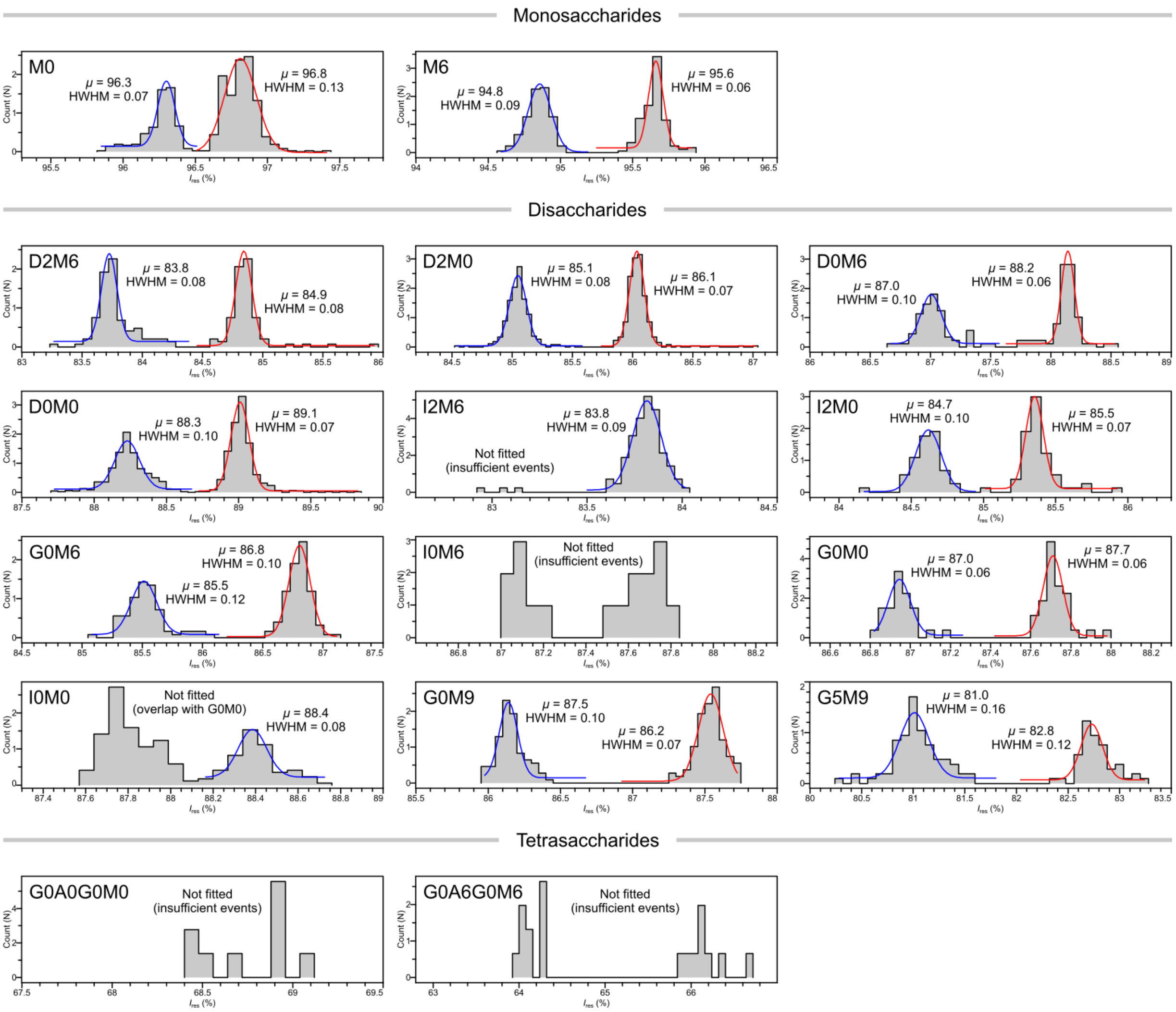

**Fig. S6 | $I_{res}$ levels are measured with high precision (αHL-Cys115).** Representative $I_{res}$ data for each AMan standard (including G5M9) acquired from single pores are overlaid with single-term Gaussian fits (one for each level or 'half-doublet') computed using the Clampfit Fit function with bin widths of 0.04–0.08 %, where all other parameters were set to their default values. Gaussian s.d. values were multiplied by 1.1774 to yield HWHM values. $\mu$, centroid. $\mu$ and HWHM values are reported to one and two decimal places, respectively (also see Table S6).

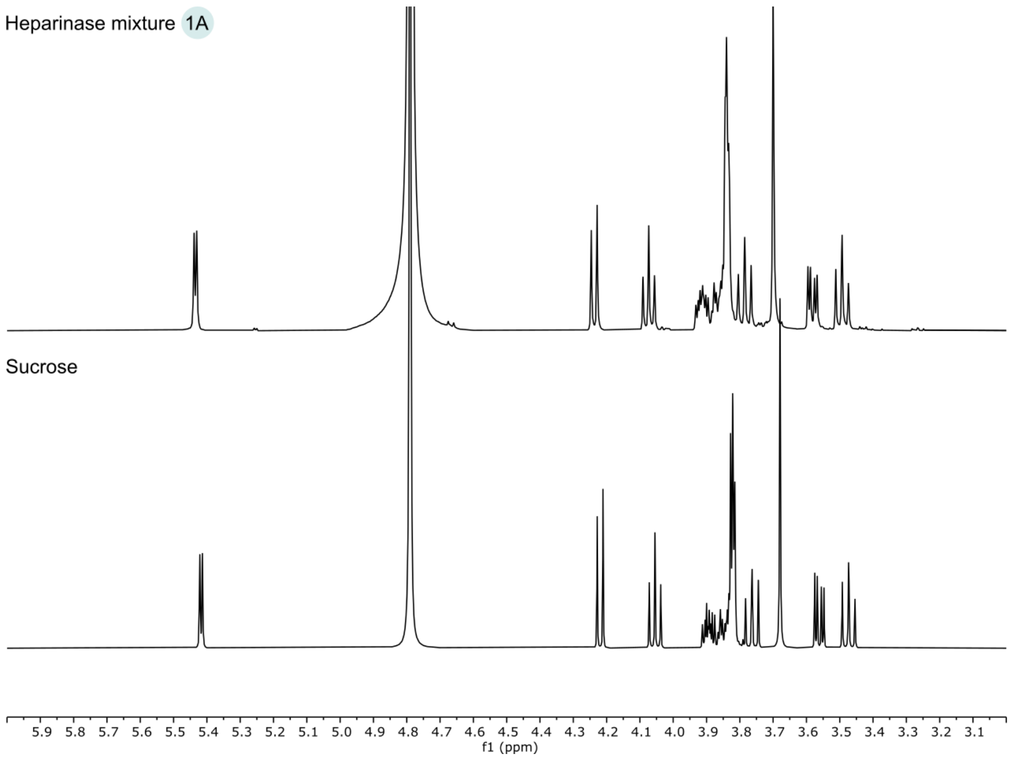

**Fig. S7 | Commercial enzyme formulations contain sucrose as a cryoprotectant.** $^{1}H$ NMR spectra (500 MHz, $D_2O$, 298 K) of a mixture of commercial heparinases I, II, and III, prepared as detailed in Methods, after condition **1A** (top) and a sucrose standard (bottom). Chemical shifts were referenced to the residual HOD signal. Sucrose signals in the spectra are slightly offset due to a difference in pD.

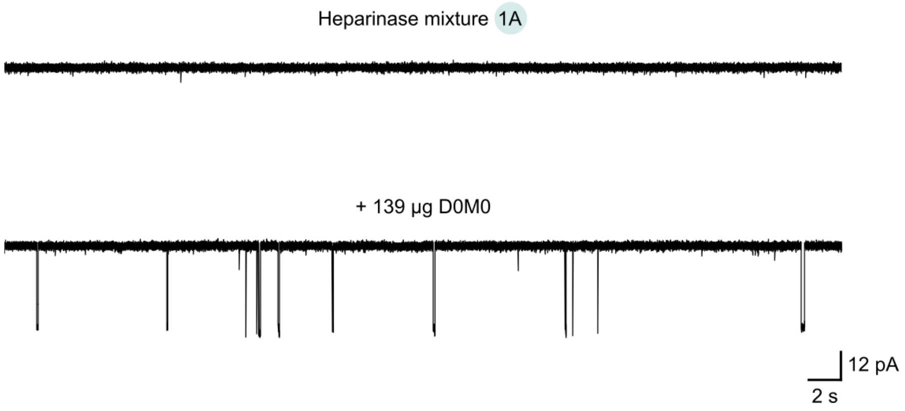

**Fig. S8 | Buffer salts, enzymes, and cryoprotectant sugars do not interfere with sensing.** A heparinase mixture (Methods and Fig. S7) was subjected to condition **1A**; 20 µL of solution was subsequently directly introduced to the αHL-Cys115 pore. No covalent events were observed. The subsequent addition of D0M0 (139 µg) yielded numerous covalent events, thereby confirming pore viability. **Recording conditions:** 4 M LiCl, 20 mM HEPBS, 40 µM EDTA, titrated to pH 8 using KOH; analyte (*cis*); +150 mV (*trans*); 23.8 ± 1 °C.

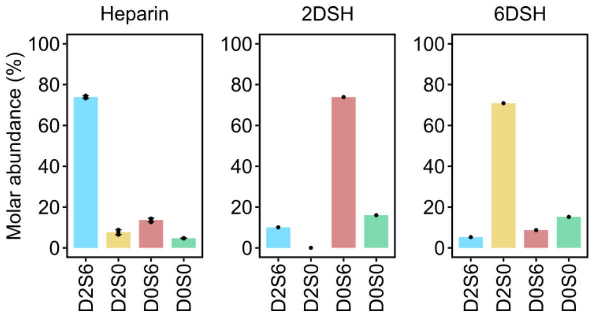

**Fig. S9 | Disaccharide compositions of pUFH, 2DSH, and 6DSH determined by HPLC.** Compositions of porcine intestinal heparin (here 'heparin'), measured by RPIP-HPLC following eliminative digestion, were obtained from ref. [46], where bars represent means across three samples from different manufacturers, points indicate measurements from independent runs, and vertical error bars (not visible at the present scale) indicate s.d. Compositions of 2DSH and 6DSH, measured by strong anion-exchange HPLC following eliminative digestion, were provided by the supplier, with the extent of 6DSH digestion specified as 93 %. The 2DSH and 6DSH bar plots show data from single HPLC runs.

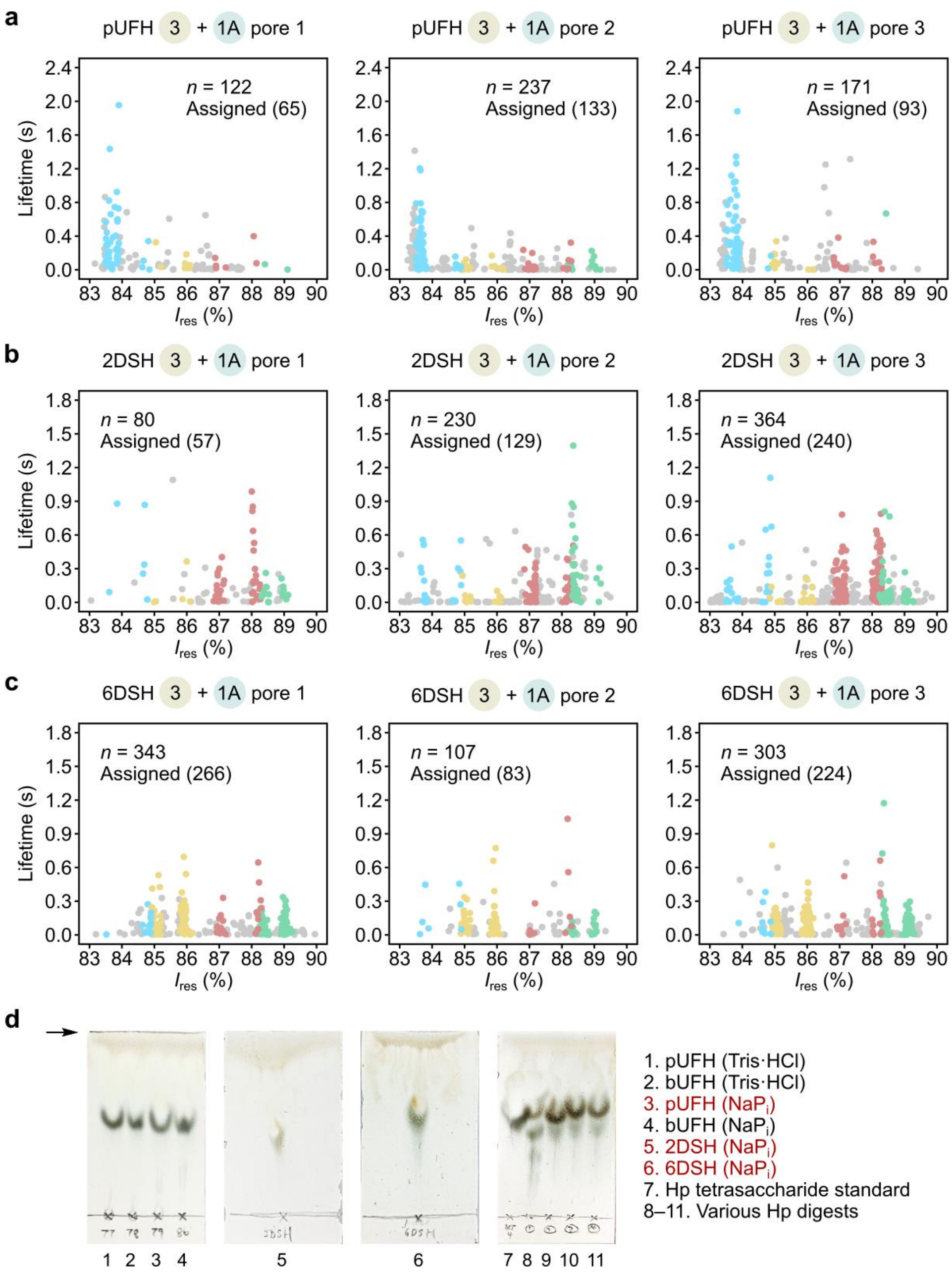

**Fig. S10 | Nanopore fingerprints of pUFH, 2DSH, and 6DSH with all disaccharide events. a–c,** pUFH (**a**), 2DSH (**b**), and 6DSH (**c**) fingerprints were generated using the αHL-Cys115 pore under conditions **3 + 1A** (Fig. 3 and Table S8). The scatter plots show event levels: D2M6 (blue), D2M0 (yellow), D0M6 (red), D0M0 (green), all other disaccharides (grey). *n*, total number of events. The number of assigned events is given in parentheses. **d,** Analysis of pUFH, 2DSH, 6DSH, unfractionated bovine lung heparin (bUFH), and Hp digests using high-performance thin-layer chromatography according to a literature protocol[47]. The heparins were digested under various conditions; 'NaPi' refers to the general procedure (Methods), whereas 'Tris·HCl' refers to a variant that used a 10 mM Tris·HCl pH 7 buffer but was otherwise identical. The presence of a single spot after each chromatographic run indicated exhaustive digestion. For comparison, a series of incomplete Hp digests, prepared under alternative digestion conditions, is shown. Digests that were subsequently ring-contracted are indicated in red. The arrow indicates the position of the mobile phase front.

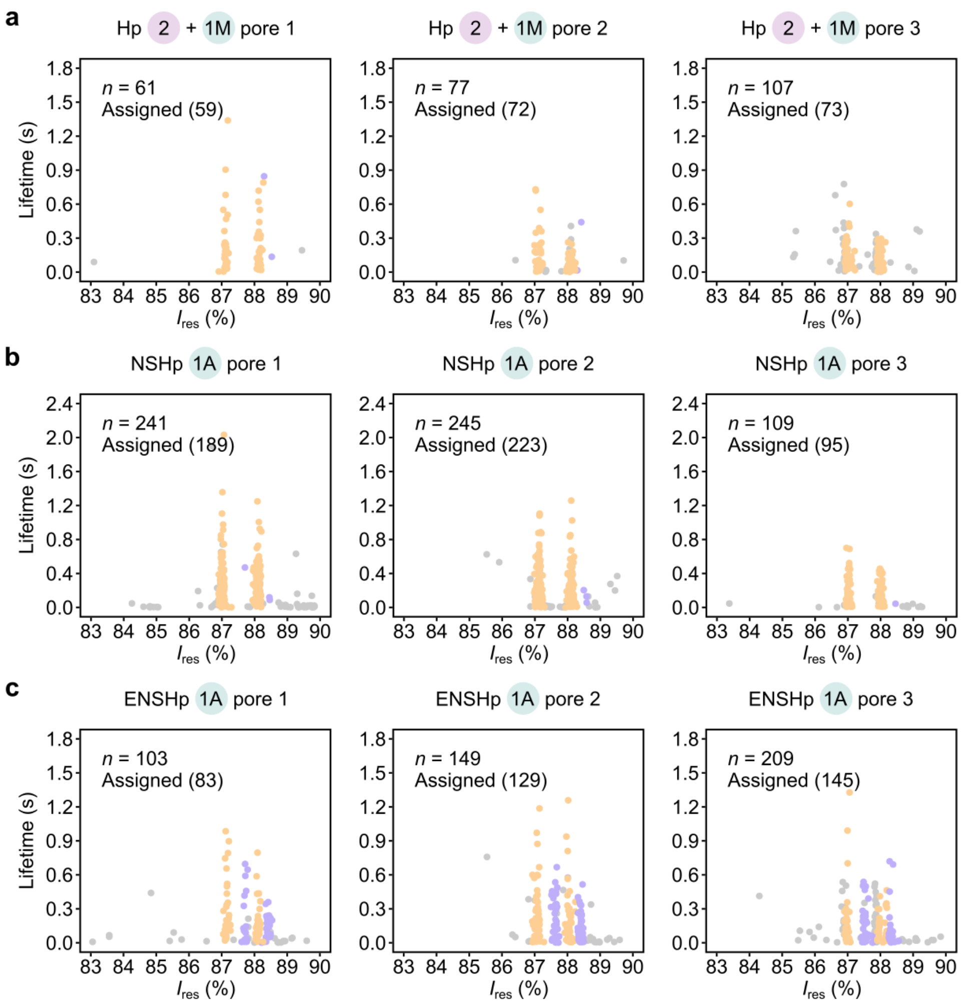

**Fig. S11 | Nanopore fingerprints of Hp, NSHp, and ENSHp with all disaccharide events. a–c,** Hp (**a**), NSHp (**b**), and ENSHp (**c**) fingerprints were generated using the αHL-Cys117 pore under condition **1A** or conditions **2 + 1M** (Fig. 4 and Table S9). The scatter plots show event levels: G0M0 (orange), I0M0 (purple), all other disaccharides (grey). *n*, total number of events. The number of assigned events is given in parentheses.

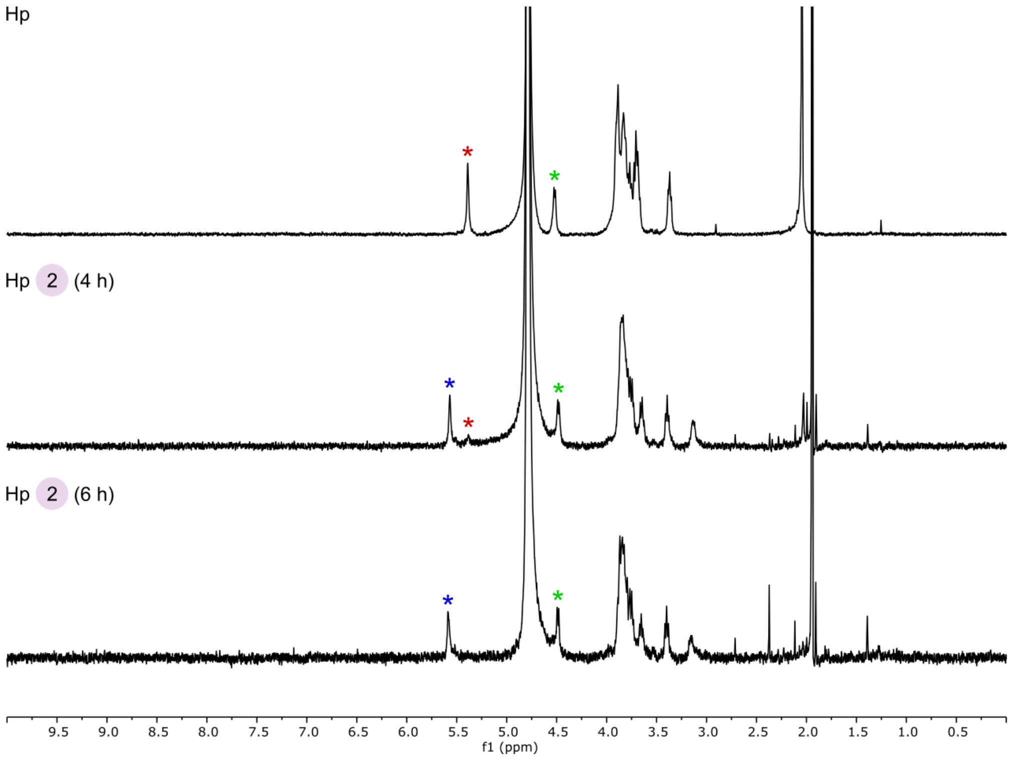

**Fig. S12 | Hp before and after hydrazinolysis.** $^{1}H$ NMR spectra (500 MHz, $D_2O$, 298 K) of Hp before and after hydrazinolysis, with reaction times indicated on the panels. Chemical shifts were referenced to the residual HOD signal. The coloured asterisks indicate the anomeric signals of GlcNAc (red), GlcN (blue), and GlcA residues (green).

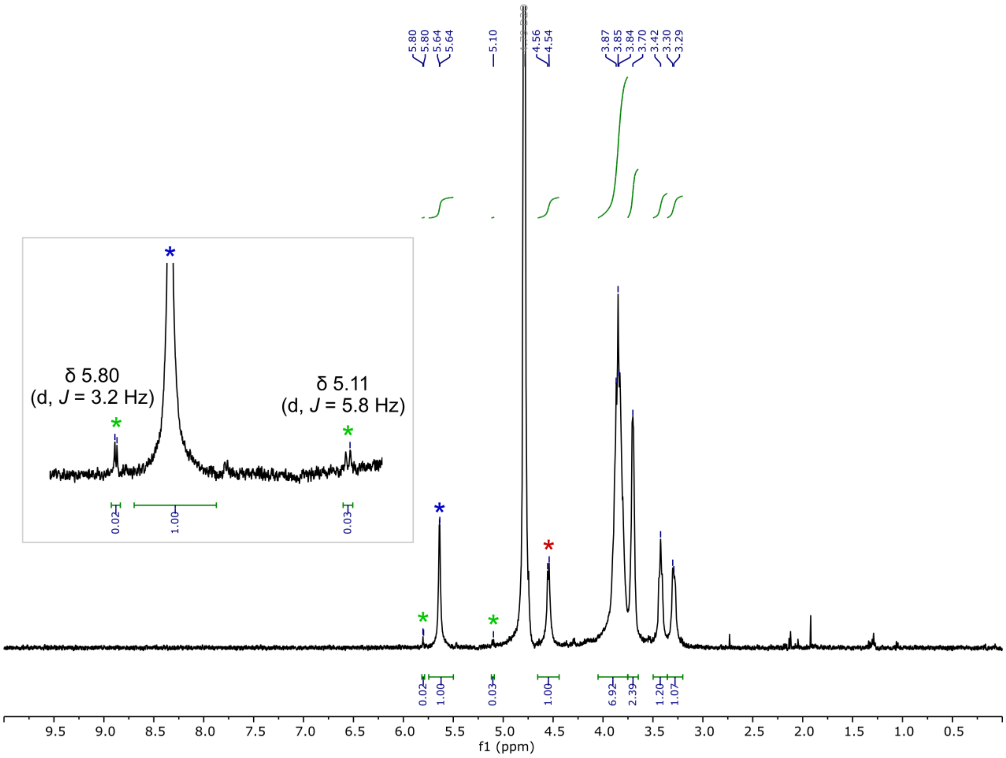

**Fig. S13 | NSHp.** $^1$H NMR spectrum (500 MHz, $D_2O$, 298 K) of NSHp; chemical shifts were referenced to the residual HOD signal. The blue and red asterisks indicate the anomeric signals of GlcNS and GlcA residues, respectively, whereas the green asterisks indicate ΔUA signals, present at ~2 % abundance (for reference, see D0m6 and D0m0 NMR data in Supplementary Methods). IdoA signals were not observed (hence, the degree of epimerization is zero).

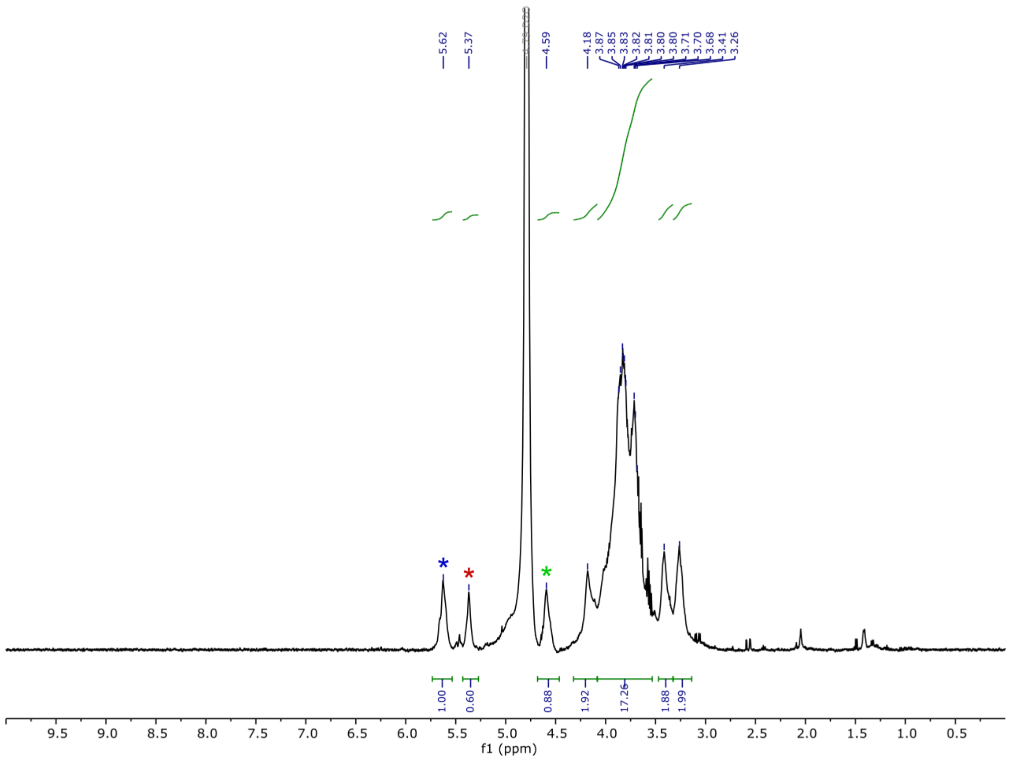

**Fig. S14 | ENSHp.** $^{1}$H NMR spectrum (500 MHz, $D_2O$, 298 K) of ENSHp. Chemical shifts were referenced to the residual HOD signal. The coloured asterisks indicate the anomeric signals of GlcNS in the sequence GlcNS-GlcA (blue), GlcNS in the sequence GlcNS-IdoA (red), and GlcA (green); signal assignments were based on the literature[48]. The degree of epimerization, calculated from the integrals of the GlcNS signals, was 38 %.

### Completely *N*-Sulfated K5 Polysaccharide, Na Salt

**Catalog Number:** C-CNSK5PS-1MG **Quantity:** 1 mg
C-CNSK5PS-2MG **Quantity:** 2 mg

**Lot Number:** 1B30C2H
**CAS Number:** None listed
**Reaction:**

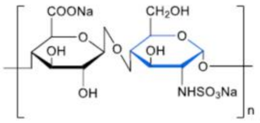

**Results:**

| Tests | Specifications | Results |
|---|---|---|
| Purity | >98% | 99% |
| Average Molecular Weight ($M_w$, Da) | ~50,000-80,000 | 59,200 |
| ($M_n$, Da) | | 57,850 |
| Uronic Acid | 30%-40% | 35% |
| *N*-sulfate content | >95% | 97% |
| Amino Content | <2% | 1% |
| *N*-acetyl Content | <2% | 1% |
| ∆ Di-NS content in disaccharide compositional analysis (%) | ≥95% | 96% |
| Structure Analysis | Pass | Conforms |
| Solubility, 70 mg/mL, $H_2O$ | Clear, colorless | Conforms |
| Appearance | White to faint yellow powder | Conforms |

1. HPGPC result of completely *N*-Sulfated K5 polysaccharide

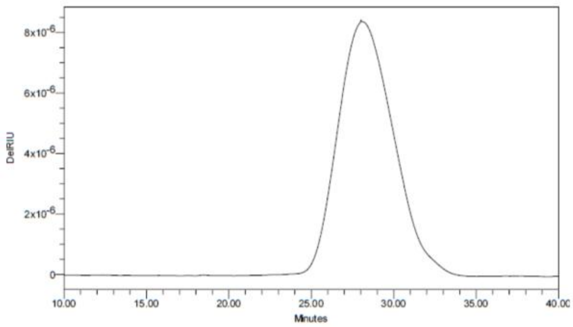

2. The [1]H NMR spectrum of completely *N*-Sulfated K5 polysaccharide

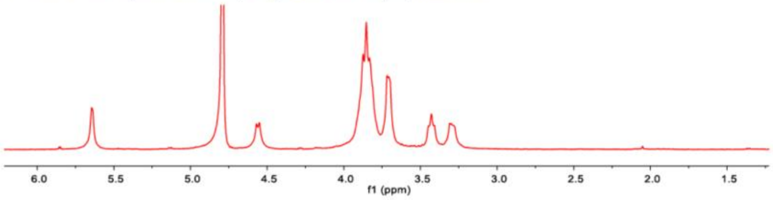

3. The [13]C NMR spectrum of the completely *N*-Sulfated K5 polysaccharide

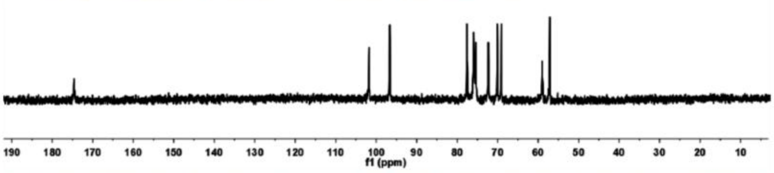

4. Unsaturated disaccharide analysis of the completely *N*-sulfated K5 polysaccharide degraded products by the mixture of recombinant heparinase II (Cat#: E-REHEPII) and heparinase III (Cat#: E-REHEPIII) with SAX-HPLC

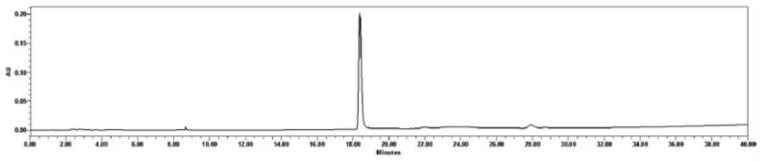

**Fig. S15 | NSHp supplier certificate.** Disaccharide compositional analysis of NSHp following eliminative digestion revealed only ΔUA-GlcNS. HPGPC, high-performance gel permeation chromatography (or SEC).

### Epimerized Completely *N*-Sulfated K5 Polysaccharide, Na Salt

GlycoNovo.com

**Catalog Number:** C-EPICNSK5PS-1MG **Quantity:** 1 mg
C-EPICNSK5PS-2MG **Quantity:** 2 mg

**Lot Number:** 1B60H1H
**CAS number:** None list
**Source:** Derived from K5 polysaccharide from *E. coli* 010:K5:H4
**Reaction:**

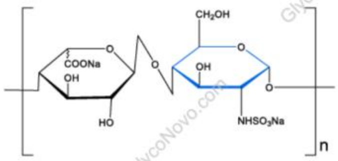

**Results:**

| Tests | Specifications | Results |
|---|---|---|
| Purity | >97% | 98.7% |
| Average Molecular Weight ($M_w$, Da) | 55,000-85,000 | 84,000 |
| Amino Content | <2% | N.D. |
| *N*-acetyl Content | <2% | N.D. |
| Sulfur Content | 3-5% | 4.8% |
| IdoA conversion rate | >40% | 46.8% |
| Structure Analysis | Pass | Pass |
| Solubility, 70 mg/mL, $H_2O$ | Clear, colorless to faint yellow | Clear, colorless |
| Appearance | White to faint yellow powder | White powder |

1. Purity analysis of the Epimerized completely *N*-sulfated K5 polysaccharide by HPGPC

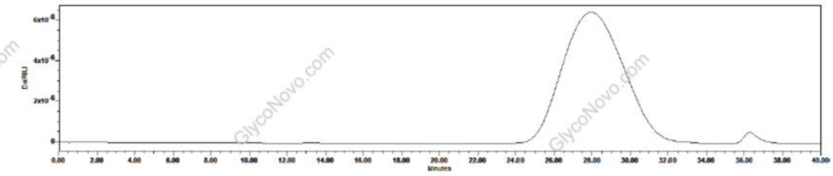

2. The ¹H-NMR spectrum of the Epimerized completely *N*-sulfated K5 polysaccharide

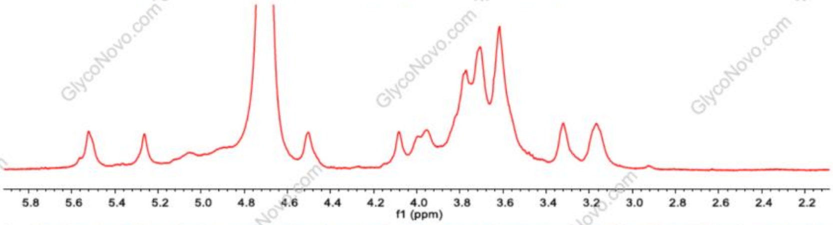

3. Unsaturated disaccharide compositional analysis of the epimerized completely N-sulfated K5 polysaccharide degraded products by mixture of recombinant heparinase II (Cat#: E-REHEPII) and heparinase III (Cat#: E-REHEPIII) with SAX-HPLC (96%)

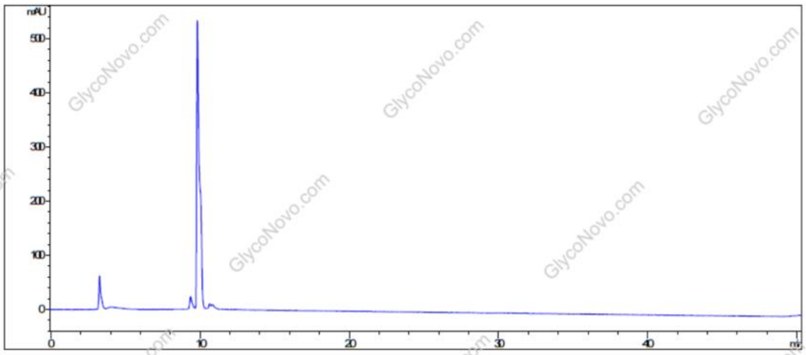

**Fig. S16 | ENSHp supplier certificate.** Disaccharide compositional analysis of ENSHp following eliminative digestion revealed only ΔUA-GlcNS; the extent of digestion was 96 %. HPGPC, high-performance gel permeation chromatography (or SEC).

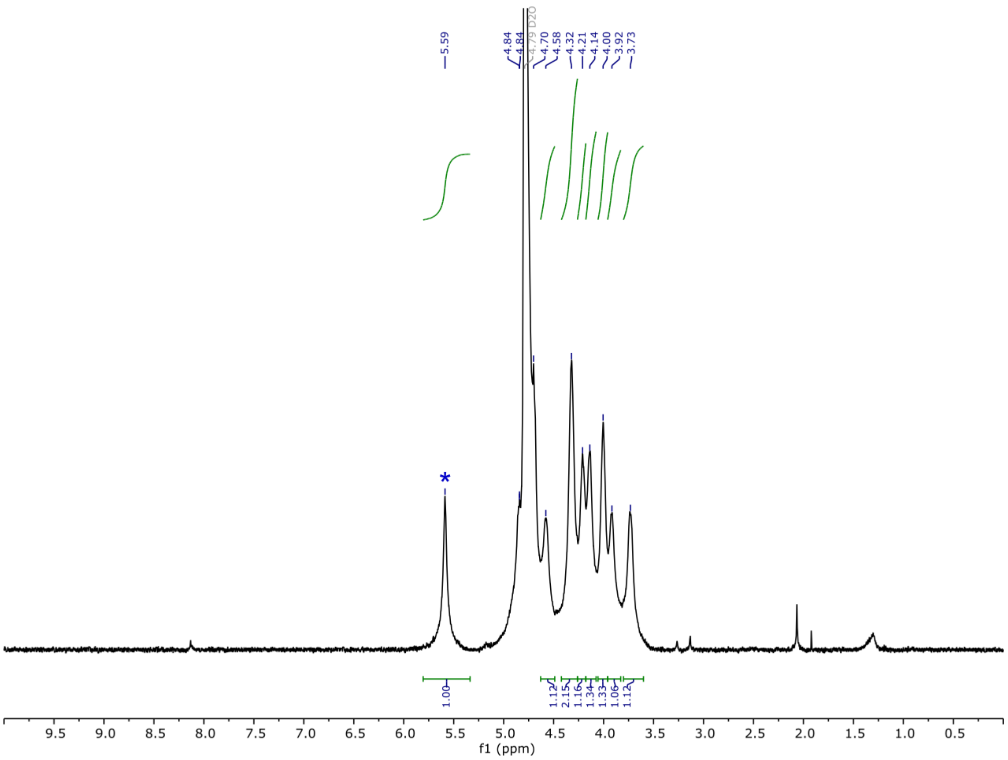

**Fig. S17 | NSOSHp.** $^1$H NMR spectrum (500 MHz, $D_2O$, 298 K) of NSOSHp. Chemical shifts were referenced to the residual HOD signal. The GlcNS3S6S anomeric signal is highlighted with a blue asterisk.

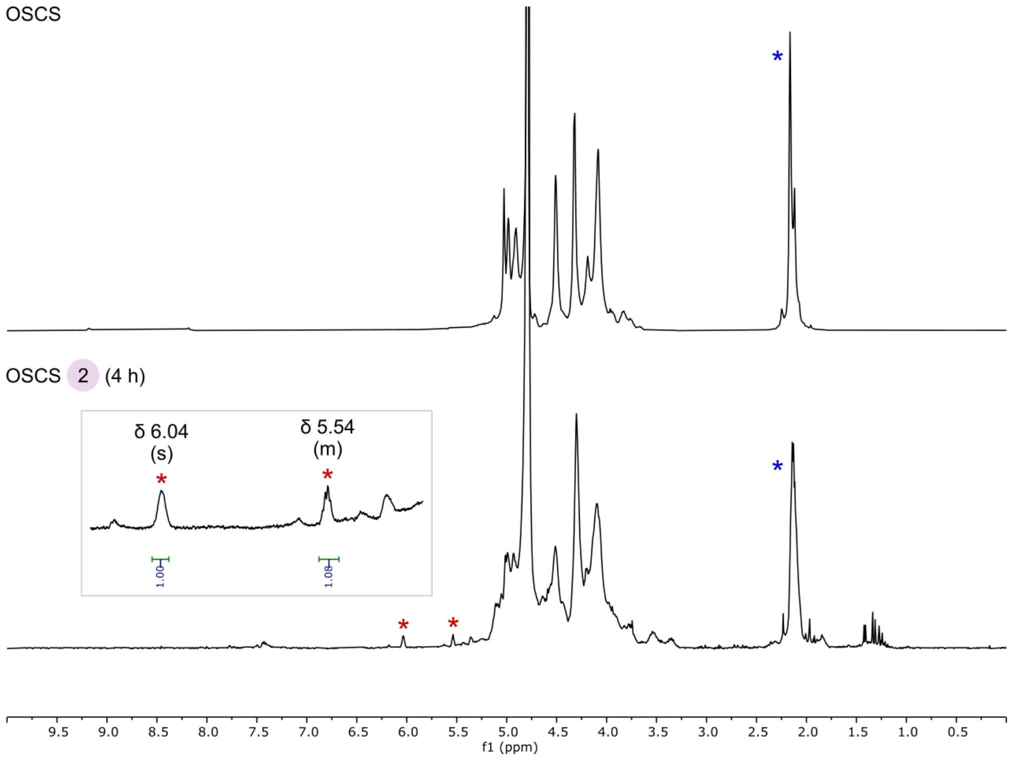

**Fig. S18 | OSCS before and after hydrazinolysis.** $^{1}H$ NMR spectra (500 MHz, $D_2O$, 298 K) of OSCS before and after hydrazinolysis and subsequent exhaustive dialysis. The reaction time is indicated on the panel. Chemical shifts were referenced to the residual HOD signal. The blue asterisks indicate the *N*-acetyl signals of *N*-acetylgalactosamine (GalNAc) residues, whereas red asterisks indicate signals consistent with ΔUA residues (for reference, see D2m6 NMR data in Supplementary Methods). The degree of OSCS deacetylation could not be accurately determined.

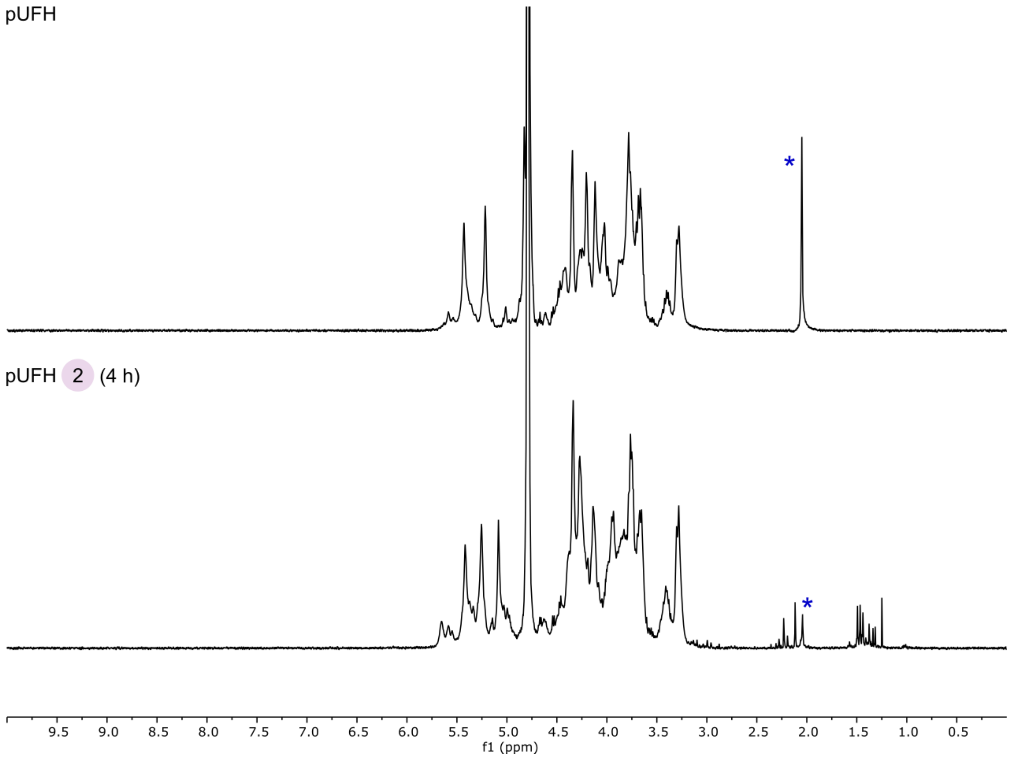

**Fig. S19 | pUFH before and after hydrazinolysis.** $^{1}$H NMR spectra (500 MHz, $D_2O$, 298 K) of pUFH before and after hydrazinolysis and subsequent exhaustive dialysis. The reaction time is indicated on the panel. Chemical shifts were referenced to the residual HOD signal. The blue asterisks indicate the *N*-acetyl signals of GlcNAc residues.

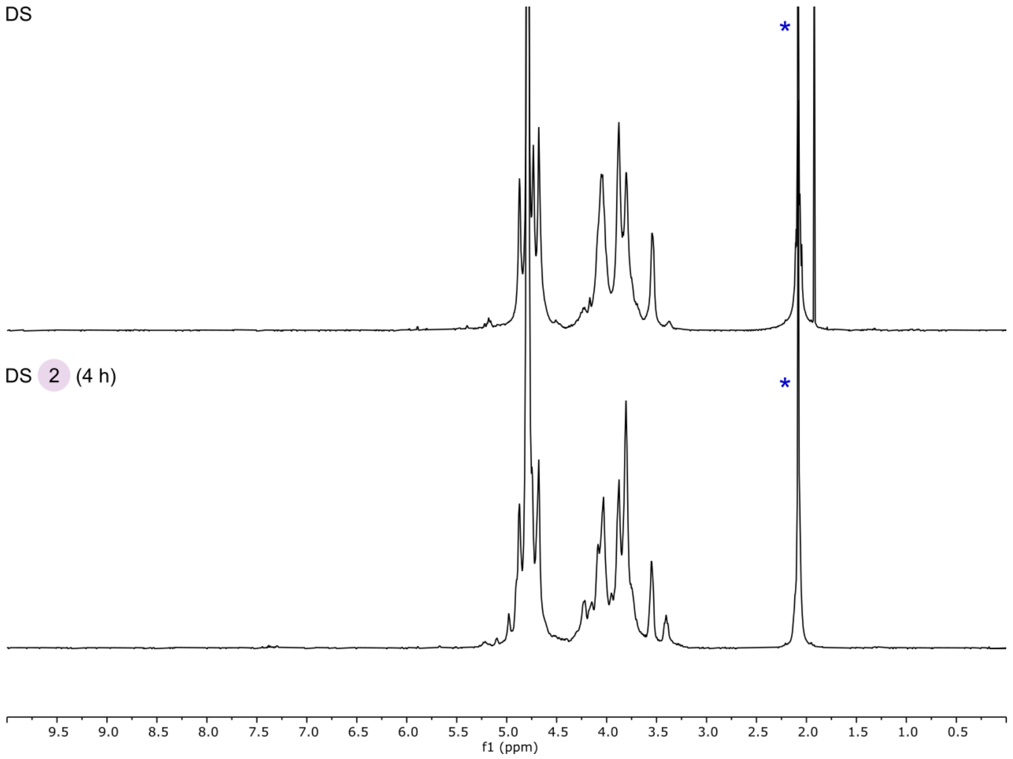

**Fig. S20 | DS before and after hydrazinolysis.** $^{1}H$ NMR spectra (500 MHz, $D_2O$, 298 K) of DS before and after hydrazinolysis and subsequent exhaustive dialysis. The reaction time is indicated on the panel. Chemical shifts were referenced to the residual HOD signal. The blue asterisks indicate the *N*-acetyl signals of GalNAc residues.

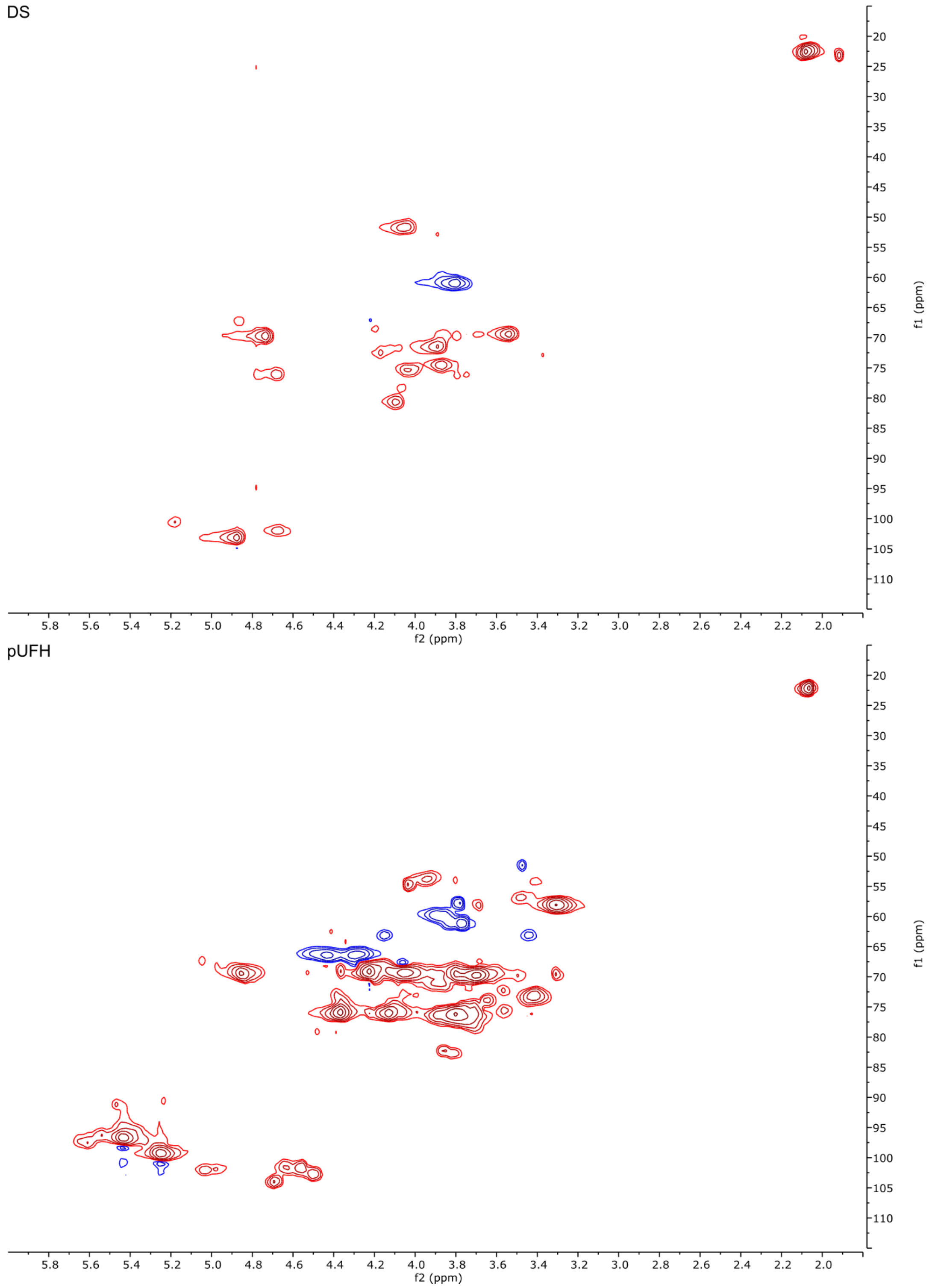

**Fig. S21 | Dermatan sulfate was not detected in pUFH by two-dimensional NMR spectroscopy.** Heteronuclear single quantum coherence (HSQC) spectra of a dermatan sulfate standard (DS; top) and pUFH (bottom), recorded under the following conditions**:** 500 MHz, $D_2O$, 298 K, 10 mg of DS or 40 mg of pUFH, 80 scans. Red and blue contours indicate positive (CH and $CH_3$) and negative ($CH_2$) signals, respectively.

| Disaccharide | Sequence | Molecular mass[a] (Da) | Reference |
|---|---|---|---|
| Non-3-*O*-sulfated | | | |
| G0S0 | GlcA-GlcNS | 415.32 | Esko *et al.* (2002) Thacker *et al.* (2014) |
| G0A0 | GlcA-GlcNAc | 378.31 | |
| G0H0 | GlcA-GlcN | 337.28 | |
| G0S6 | GlcA-GlcNS6S | 494.37 | |
| G0A6 | GlcA-GlcNAc6S | 457.36 | |
| G2S0 | GlcA2S-GlcNS | 494.37 | |
| G2S6 | GlcA2S-GlcNS6S | 573.42 | |
| I0S0 | IdoA-GlcNS | 415.32 | |
| I0A0 | IdoA-GlcNAc | 378.31 | |
| I0S6 | IdoA-GlcNS6S | 494.37 | |
| I0A6 | IdoA-GlcNAc6S | 457.36 | |
| I2S0 | IdoA2S-GlcNS | 494.37 | |
| I2A0 | IdoA2S-GlcNAc | 457.36 | |
| I2S6 | IdoA2S-GlcNS6S | 573.42 | |
| I2A6 | IdoA2S-GlcNAc6S | 536.41 | |
| 3-*O*-sulfated | | | |
| G0S3 | GlcA-GlcNS3S | 494.37 | |
| G0S9 | GlcA-GlcNS3S6S | 573.42 | |
| G2S3 | GlcA2S-GlcNS3S | 573.42 | |
| I0S3 | IdoA-GlcNS3S | 494.37 | |
| I0S9 | IdoA-GlcNS3S6S | 573.42 | |
| I2S3 | IdoA2S-GlcNS3S | 573.42 | |
| I2H3 | IdoA2S-GlcN3S | 495.38 | |
| I2H9 | IdoA2S-GlcN3S6S | 574.43 | |
| I2H0 | IdoA2S-GlcN | 416.33 | Westling *et al.* (2002) |
| I2H6 | IdoA2S-GlcN6S | 495.38 | |
| G0H6 | GlcA-GlcN6S | 416.33 | Shi *et al.* (2009) |
| I2S9 | IdoA2S-GlcNS3S6S | 652.47 | Thacker *et al.* (2014) |

[a] Calculated for disaccharides in their physiological protonation states positioned internally in a chain

**Table S1 | Disaccharides that emerge from HS biosynthesis.** A total of 27 unique disaccharide units have been identified as products of HS biosynthesis[49–52]. Disaccharides are referred to by their disaccharide structure code[53] and colour-coded by molecular mass.

| ΔUA disaccharide | Sequence | Molecular mass[a] (Da) |
|---|---|---|
| Non-3-*O*-sulfated | | |
| D0S0 | ΔUA-GlcNS | 415.32 |
| D0A0 | ΔUA-GlcNAc | 378.31 |
| D0H0 | ΔUA-GlcN | 337.28 |
| D0S6 | ΔUA-GlcNS6S | 494.37 |
| D0A6 | ΔUA-GlcNAc6S | 457.36 |
| D0H6 | ΔUA-GlcN6S | 416.33 |
| D2S0 | ΔUA2S-GlcNS | 494.37 |
| D2A0 | ΔUA2S-GlcNAc | 457.36 |
| D2H0 | ΔUA2S-GlcN | 416.33 |
| D2S6 | ΔUA2S-GlcNS6S | 573.42 |
| D2A6 | ΔUA2S-GlcNAc6S | 536.41 |
| D2H6 | ΔUA2S-GlcN6S | 495.38 |
| 3-*O*-sulfated | | |
| D0S3 | ΔUA-GlcNS3S | 494.37 |
| D0S9 | ΔUA-GlcNS3S6S | 573.42 |
| D2S3 | ΔUA2S-GlcNS3S | 573.42 |
| D2H3 | ΔUA2S-GlcN3S | 495.38 |
| D2S9 | ΔUA2S-GlcNS3S6S | 652.47 |
| D2H9 | ΔUA2S-GlcN3S6S | 574.43 |

[a] Calculated for disaccharides in their physiological protonation states

**Table S2 | ΔUA disaccharides derived from the biosynthetic HS disaccharide pool.** The eliminative digestion of HS generates 18 unique ΔUA disaccharides. Disaccharides are referred to by their disaccharide structure code[53] and colour-coded by molecular mass.

| AMan disaccharide | Sequence | Molecular mass[a] (Da) |
|---|---|---|
| Non-3-*O*-sulfated | | |
| G0M0 | GlcA-AMan | 337.26 |
| G0M6 | GlcA-AMan6S | 416.31 |
| G2M0 | GlcA2S-AMan | 416.31 |
| G2M6 | GlcA2S-AMan6S | 495.36 |
| I0M0 | IdoA-AMan | 337.26 |
| I0M6 | IdoA-AMan6S | 416.31 |
| I2M0 | IdoA2S-AMan | 416.31 |
| I2M6 | IdoA2S-AMan6S | 495.36 |
| 3-*O*-sulfated | | |
| G0M3 | GlcA-AMan3S | 416.31 |
| G0M9 | GlcA-AMan3S6S | 495.36 |
| G2M3 | GlcA2S-AMan3S | 495.36 |
| I0M3 | IdoA-AMan3S | 416.31 |
| I0M9 | IdoA-AMan3S6S | 495.36 |
| I2M3 | IdoA2S-AMan3S | 495.36 |
| I2M9 | IdoA2S-AMan3S6S | 574.41 |

[a] Calculated for disaccharides in their physiological protonation states

**Table S3 | AMan disaccharides derived from the biosynthetic HS disaccharide pool.** Tiffeneau–Demjanov ring contraction of HS generates 15 unique AMan disaccharides. Disaccharides are referred to by their disaccharide structure code[53] and colour-coded by molecular mass.

| Sample | Disaccharides[a] | Molar abundance[b] (%) | Reference |
|---|---|---|---|
| Bovine aortic endothelial cell HS | I2M6, I2M0, G0M6, I0M6, G0M9 | 89–91 | Marcum *et al*. (1986) |
| *Tivela mactroides* heparin | I2M6, I2M0, G0M6, I0M6, G0M9 | 94–95 | Pejler *et al*. (1987) |
| *Anomalocardia brasiliana* heparin | | 84 | |
| Porcine intestinal HS | I2M6, I2M0, G0M6, I0M6, G0M0, I0M0, G0M9 | 98 | Guo *et al*. (1989) |
| Rat microvascular endothelial cell HS | I2M6, I2M0, G0M6, I0M6, G0M9 | 87–95 | Kojima *et al*. (1992) |
| CHO cell HS | I2M6, I2M0, G0M6, I0M6, G0M0, I0M0 | 97–100 | Bai *et al*. (1996) |
| Rat granulosa cell HS | I2M6, I2M0, G0M6, I0M6, G0M9 | 92–100 | Hosseini *et al*. (1996) |
| Mouse mastocytoma cell heparin | I2M6, I2M0, G0M6, I0M6, G0M0, I0M0, G0M9 | ~100 | Uhlin-Hansen *et al*. (1997) |
| Bovine renal HS | I2M6, I2M0, G0M6, I0M6, G0M0, I0M0 | ~100 | Kariya *et al*. (1998) |
| Bovine renal HS | I2M6, I2M0, G0M6, I0M6, G0M0, I0M0, G0M9 | 99 | Gill *et al*. (2012) |
| Porcine intestinal HS | | 91 | |
| Porcine intestinal heparin | | 92 | |

[a] AMan disaccharides also examined in the present work
[b] AMan disaccharides in [a] relative to total disaccharides in sample

**Table S4 | Disaccharide molar abundances in diversely surveyed HS.** HS and heparin polysaccharides were quantitatively converted to AMan disaccharides either via hydrazinolysis followed by ring contraction at all GlcNS and GlcN residues[17,54–60] or, in the specific cases of *Tivela mactroides* and *Anomalocardia brasiliana* heparin, solely via ring contraction at GlcNS[61]. The seven saturated AMan UA disaccharides examined in the present work collectively account for ~84–100 % by molar abundance of all disaccharides found in these samples. CHO, Chinese hamster ovary.

| | Level 1 $I_{res}$ (%) | | Level 2 $I_{res}$ (%) | |
|---|---|---|---|---|
| | Mean[a] | HWHM[a] | Mean[a] | HWHM[a] |
| Monosaccharides | | | | |
| M0 | 96.6 ± 0.0 | 0.07 ± 0.01 | 96.2 ± 0.0 | 0.06 ± 0.00 |
| M6 | 94.4 ± 0.1 | 0.08 ± 0.01 | 93.8 ± 0.1 | 0.08 ± 0.01 |
| ΔUA disaccharides | | | | |
| D2M6 | 85.0 ± 0.1 | 0.13 ± 0.03 | 83.4 ± 0.1 | 0.12 ± 0.03 |
| D2M0 | 86.7 ± 0.1 | 0.10 ± 0.03 | 85.1 ± 0.1 | 0.10 ± 0.03 |
| D0M6 | 87.8 ± 0.1 | 0.08 ± 0.01 | 86.8 ± 0.1 | 0.09 ± 0.02 |
| D0M0 | 89.4 ± 0.0 | 0.07 ± 0.02 | 88.5 ± 0.0 | 0.09 ± 0.04 |
| UA disaccharides | | | | |
| I2M6 | 84.1 ± 0.1 | 0.12 ± 0.04 | 82.4 ± 0.1 | 0.08 ± 0.03 |
| I2M0 | 86.0 ± 0.0 | 0.07 ± 0.01 | 84.2 ± 0.0 | 0.08 ± 0.02 |
| G0M6 | 86.3 ± 0.1 | 0.07 ± 0.01 | 85.5 ± 0.1 | 0.10 ± 0.07 |
| I0M6* | 86.7 ± 0.2 | 0.18 ± 0.01 | 85.9 ± 0.1 | 0.13 ± 0.01 |
| G0M0 | 88.1 ± 0.1 | 0.06 ± 0.01 | 87.1 ± 0.1 | 0.05 ± 0.01 |
| I0M0* | 88.4 ± 0.1 | 0.09 ± 0.02 | 87.6 ± 0.1 | 0.08 ± 0.03 |
| G0M9 | 87.1 ± 0.0 | 0.10 ± 0.02 | | |
| Tetrasaccharides | | | | |
| G0A0G0M0* | 66.5 ± 0.0 | 0.13 ± 0.11 | 64.6 ± 0.0 | 0.07 ± 0.04 |
| G0A6G0M6* | 64.3 ± 0.1 | 0.15 ± 0.05 | 63.1 ± 0.1 | 0.16 ± 0.08 |

[a] Values represent the Gaussian centroid and HWHM; for analytes with low event counts (indicated by an asterisk), values represent the arithmetic mean and a proxy HWHM calculated from the sample s.d.

**Table S5 | AMan mono-, di-, and tetrasaccharide $I_{res}$ statistics (αHL-Cys117).** For each sugar with paired interconverting levels, level 1 was chosen as the level with the higher $I_{res}$ value. Means and HWHM values are reported to one and two decimal places, respectively; all values represent mean ± s.d. across $N \geq 3$ pores, except for I0M6 and G0A6G0M6 ($N$ = 2) due to limited material.

| | Level 1 $I_{res}$ (%) | | Level 2 $I_{res}$ (%) | |
|---|---|---|---|---|
| | Mean[a] | HWHM[a] | Mean[a] | HWHM[a] |
| Monosaccharides | | | | |
| M0 | 96.8 ± 0.1 | 0.09 ± 0.04 | 96.3 ± 0.1 | 0.09 ± 0.04 |
| M6 | 95.8 ± 0.1 | 0.06 ± 0.02 | 95.0 ± 0.1 | 0.09 ± 0.01 |
| ΔUA disaccharides | | | | |
| D2M6 | 84.8 ± 0.1 | 0.08 ± 0.01 | 83.7 ± 0.1 | 0.08 ± 0.03 |
| D2M0 | 86.0 ± 0.1 | 0.07 ± 0.00 | 85.0 ± 0.1 | 0.10 ± 0.02 |
| D0M6 | 88.2 ± 0.1 | 0.10 ± 0.03 | 87.0 ± 0.1 | 0.11 ± 0.01 |
| D0M0 | 89.1 ± 0.1 | 0.08 ± 0.01 | 88.4 ± 0.1 | 0.09 ± 0.03 |
| UA disaccharides | | | | |
| I2M6 | 84.0 ± 0.1 | 0.16 ± 0.06 | 83.0 ± 0.1 | 0.12 ± 0.01 |
| I2M0 | 85.4 ± 0.1 | 0.07 ± 0.01 | 84.6 ± 0.1 | 0.09 ± 0.02 |
| G0M6 | 86.7 ± 0.2 | 0.09 ± 0.03 | 85.4 ± 0.2 | 0.10 ± 0.03 |
| I0M6* | 87.8 ± 0.1 | 0.15 ± 0.04 | 87.1 ± 0.1 | 0.11 ± 0.10 |
| G0M0 | 87.9 ± 0.1 | 0.06 ± 0.00 | 87.0 ± 0.1 | 0.08 ± 0.02 |
| I0M0[b] | 88.5 ± 0.1 | 0.10 ± 0.02 | 87.9 ± 0.1 | |
| G0M9 | 87.5 ± 0.0 | 0.09 ± 0.01 | 86.2 ± 0.0 | 0.09 ± 0.02 |
| G5M9 | 82.6 ± 0.1 | 0.12 ± 0.01 | 80.8 ± 0.2 | 0.16 ± 0.01 |
| Tetrasaccharides | | | | |
| G0A0G0M0* | 68.8 ± 0.1 | 0.27 ± 0.03 | | |
| G0A6G0M6* | 66.3 ± 0.1 | 0.10 ± 0.06 | 64.4 ± 0.2 | 0.13 ± 0.04 |

[a] Values represent the Gaussian centroid and HWHM; for analytes with low event counts (indicated by an asterisk), values represent the arithmetic mean and a proxy HWHM calculated from the sample s.d.
[b] I0M0 level 2 HWHM not calculated due to overlap with G0M0 level 1

**Table S6 | AMan mono-, di-, and tetrasaccharide $I_{res}$ statistics (αHL-Cys115).** For each sugar with paired interconverting levels, level 1 was chosen as the level with the higher $I_{res}$ value. Means and HWHM values are reported to one and two decimal places, respectively; all values represent mean ± s.d. across $N \geq 3$ pores, except for I2M0, G0M6, and G0A0G0M0 ($N = 2$) due to limited material.

| | Events | Event proportion (%) Mono[a] | Di[b] | Oligo[c] (tetra[d]) |
|---|---|---|---|---|
| Pore 1 | 307 | 11.7 | 68.1 | 20.2 (16.0) |
| Pore 2 | 407 | 9.1 | 73.5 | 17.4 (13.3) |
| Pore 3 | 329 | 9.4 | 68.7 | 21.9 (14.0) |
| Mean | | 10.1 | 70.1 | 19.8 (14.4) |
| s.d. | | 1.4 | 2.9 | 2.2 (1.4) |

[a] $I_{res}$ 90–97 %
[b] $I_{res}$ 80–90 %
[c] $I_{res}$ < 80 %
[d] $I_{res}$ 60–70 %

**Table S7 | Mono-, di-, and oligosaccharide event proportions for ring-contracted pUFH (condition 1A).** Sugars were probed by the αHL-Cys115 pore (Extended Data Figs 4c–d).

| | | Events | Assigned | % assigned | Event proportion (%) D2M6 | D2M0 | D0M6 | D0M0 |
|---|---|---|---|---|---|---|---|---|
| pUFH | Pore 1 | 122 | 65 | 53.3 | 75.4 | 12.3 | 9.2 | 3.1 |
| | Pore 2 | 237 | 133 | 56.1 | 60.9 | 15.0 | 18.0 | 6.0 |
| | Pore 3 | 171 | 93 | 54.4 | 71.0 | 12.9 | 15.1 | 1.1 |
| | Mean | | | 54.6 | 69.1 | 13.4 | 14.1 | 3.4 |
| | s.d. | | | 1.4 | 7.4 | 1.4 | 4.5 | 2.5 |
| 2DSH | Pore 1 | 80 | 57 | 71.3 | 10.5 | 8.8 | 61.4 | 19.3 |
| | Pore 2 | 230 | 129 | 56.1 | 14.7 | 10.9 | 41.9 | 32.6 |
| | Pore 3 | 364 | 240 | 65.9 | 9.2 | 10.0 | 62.1 | 18.8 |
| | Mean | | | 64.4 | 11.5 | 9.9 | 55.1 | 23.5 |
| | s.d. | | | 7.7 | 2.9 | 1.0 | 11.5 | 7.8 |
| 6DSH | Pore 1 | 343 | 266 | 77.6 | 6.4 | 48.9 | 10.9 | 33.8 |
| | Pore 2 | 107 | 83 | 77.6 | 6.0 | 61.4 | 9.6 | 22.9 |
| | Pore 3 | 303 | 224 | 73.9 | 4.5 | 49.6 | 8.0 | 37.9 |
| | Mean | | | 76.3 | 5.6 | 53.3 | 9.5 | 31.6 |
| | s.d. | | | 2.1 | 1.0 | 7.1 | 1.4 | 7.8 |

**Table S8 | Disaccharide event proportions in the pUFH, 2DSH, and 6DSH fingerprints.** Fingerprints were generated using the αHL-Cys115 pore under conditions **3 + 1A** (Figs 3 and S10a–c). Events were assigned as detailed in Supplementary Methods.

| | | Events | Assigned | % assigned | Event proportion (%) G0M0 | I0M0 |
|---|---|---|---|---|---|---|
| Hp | Pore 1 | 61 | 59 | 96.7 | 96.6 | 3.4 |
| | Pore 2 | 77 | 72 | 93.5 | 97.2 | 2.8 |
| | Pore 3 | 107 | 73 | 68.2 | 100.0 | 0.0 |
| | Mean | | | 86.2 | 97.9 | 2.1 |
| | s.d. | | | 15.6 | 1.8 | 1.8 |
| NSHp | Pore 1 | 241 | 189 | 78.4 | 98.4 | 1.6 |
| | Pore 2 | 245 | 223 | 91.0 | 98.7 | 1.3 |
| | Pore 3 | 109 | 95 | 87.2 | 98.9 | 1.1 |
| | Mean | | | 85.5 | 98.7 | 1.3 |
| | s.d. | | | 6.5 | 0.3 | 0.3 |
| ENSHp | Pore 1 | 103 | 83 | 80.6 | 63.9 | 36.1 |
| | Pore 2 | 149 | 129 | 86.6 | 56.6 | 43.4 |
| | Pore 3 | 209 | 145 | 69.4 | 58.6 | 41.4 |
| | Mean | | | 78.8 | 59.7 | 40.3 |
| | s.d. | | | 8.7 | 3.7 | 3.7 |

**Table S9 | Disaccharide event proportions in the Hp, NSHp, and ENSHp fingerprints.** Fingerprints were generated using the αHL-Cys117 pore under condition **1A** or conditions **2 + 1M** (Figs 4 and S11). Events were assigned as detailed in Supplementary Methods.

### Supplementary Methods

#### 1. Preparation and sensing of AMan standards

A total of 15 AMan standards (comprising two monosaccharides, 11 disaccharides, and two tetrasaccharides) were prepared from their aminosugar precursors through the differential application of conditions **1A**, **1M**, and **2**. Six standards were obtained and subjected to covalent sensing in pure form (Fig. S22). The monosaccharide M0 was prepared from monosaccharidic GlcN. The four ΔUA disaccharides D2M6, D2M0, D0M6, and D0M0 were prepared from their *N*-sulfated precursors. G0M0 was prepared from Hp via conditions **2 + 1M**. These disaccharides were also characterized by NMR spectroscopy and MS (see below).

The remaining nine standards were obtained from ring contraction at pH 1.5 (condition **1A**) and subjected to sensing as part of mixtures of known compositions. G0M6 was prepared from the synthetic heptasaccharide 6S 7-mer C (Fig. S23). I2M0 and G0A0G0M0 were prepared from the synthetic heptasaccharide 2S 7-mer A (Fig. S24). I0M6 and G0A6G0M6 were derived from the synthetic heptasaccharide 6S IdoA 7-mer B (Fig. S25). The di- and tetrasaccharides generated from 2S 7-mer A and 6S IdoA 7-mer B produced discrete event clusters; within each mixture, the component that yielded smaller current blockades (i.e., higher $I_{res}$ values) was assigned as the disaccharide, whereas the component that yielded larger blockades (~2-fold larger in all cases) was assigned as the tetrasaccharide. The glucuronides liberated from the heptasaccharide reducing ends were not removed, as they predictably failed to elicit sensing events.

I0M0 was prepared from ENSHp under condition **1A** with the concomitant generation of G0M0; no attempt was made to fractionate the sugars, as they were amenable to simultaneous analysis. I0M0 was identified by comparing $I_{res}$ data from ENSHp and Hp (see Fig. 4d for αHL-Cys117 data).

I2M6, G0M9, and M6 were prepared from the synthetic pentasaccharide anticoagulant fondaparinux under condition **1A**. As before, no attempt was made to fractionate the sugars. Sensing of this mixture revealed three major components (Fig. S26). The component that yielded the smallest current blockades ($I_{res}$ ~95 %) was assigned as M6. I2M6 was identified by comparing $I_{res}$ data from fondaparinux and pUFH, given that standard unfractionated heparin is predominantly cleaved into I2M6 under condition **1A**[62] (see Extended Data Fig. 9b and Fig. S27 for αHL-Cys115 and αHL-Cys117 data, respectively). The remaining component in the fondaparinux-derived mixture was assigned as G0M9.

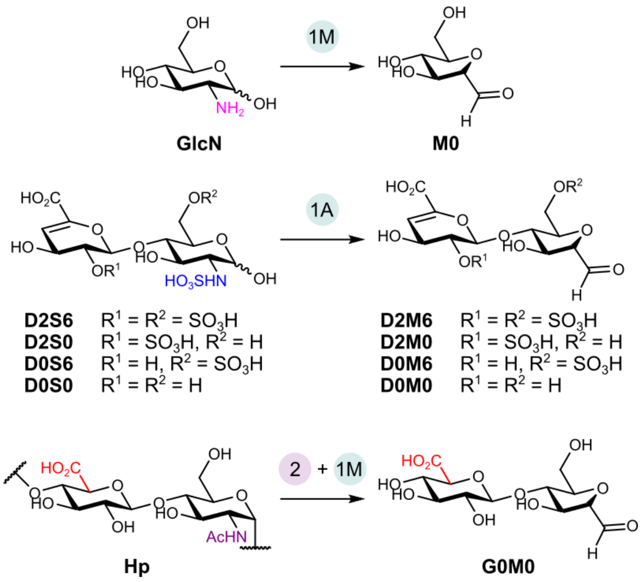

**Fig. S22 | Preparation of pure AMan standards.** Reactions were performed according to the general procedures detailed in Methods; Hp was reacted with hydrazine for 6 h instead of the default duration of 4 h (Fig. S12).

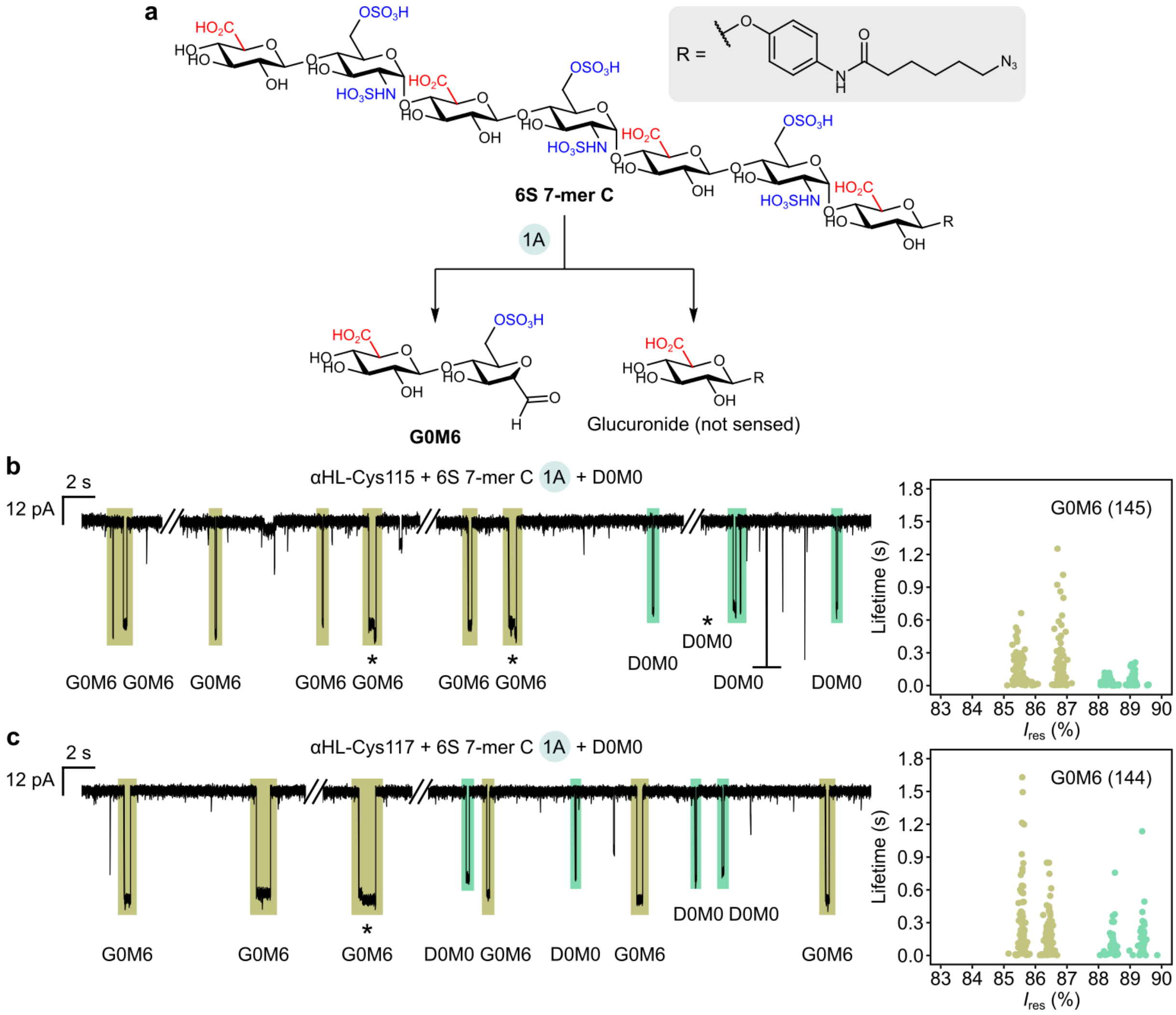

**Fig. S23 | Preparation and sensing of G0M6. a,** G0M6 was prepared from 6S 7-mer C as part of a mixture under condition **1A**. **b–c,** The mixture was directly loaded into our nanopore setup, with D0M0 serving as an internal standard. Representative current traces for the αHL-Cys115 (**b**) and αHL-Cys117 (**c**) pores are shown, along with disaccharide assignments. Interconversion events are marked with an asterisk. A graphically truncated event is marked with a horizontal bar. The scatter plots show assigned event levels from single pores, and levels are colour-coded as in the traces. The number of G0M6 events is given in parentheses. **Recording conditions:** 4 M LiCl, 20 mM HEPBS, 40 µM EDTA, titrated to pH 8 using KOH; sugars (*cis*); +150 mV (*trans*); 23.8 ± 1 °C. G0M6 and D0M0 concentrations were 0.43 mM and 0.14–0.46 mM, respectively. Events with lifetimes under 2 ms were discarded (Methods).

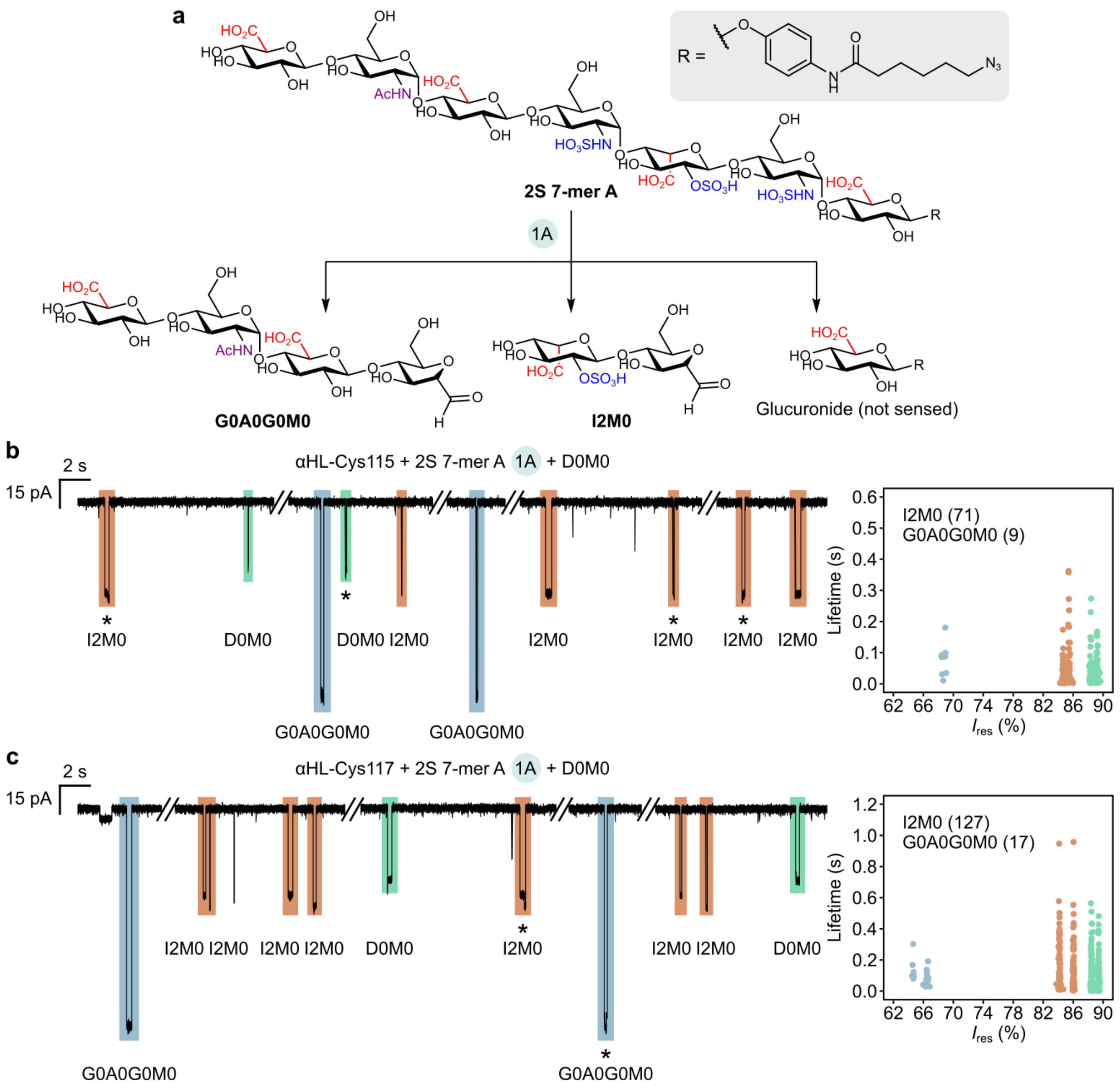

**Fig. S24 | Preparation and sensing of I2M0 and G0A0G0M0. a,** The two AMan standards were prepared from 2S 7-mer A as part of a mixture under condition **1A**. **b–c,** The mixture was directly loaded into our nanopore setup, with D0M0 serving as an internal standard. Representative current traces for the αHL-Cys115 (**b**) and αHL-Cys117 (**c**) pores are shown, along with event assignments. Interconversion events are marked with an asterisk. The scatter plots show assigned event levels from single pores, and levels are colour-coded as in the traces. The number of I2M0 and G0A0G0M0 events is given in parentheses. **Recording conditions:** 4 M LiCl, 20 mM HEPBS, 40 µM EDTA, titrated to pH 8 using KOH; sugars (*cis*); +150 mV (*trans*); 23.8 ± 1 °C. I2M0, G0A0G0M0, and D0M0 concentrations were 0.19 mM, 0.19 mM, and 0.23 mM, respectively. Events with lifetimes under 2 ms were discarded (Methods).

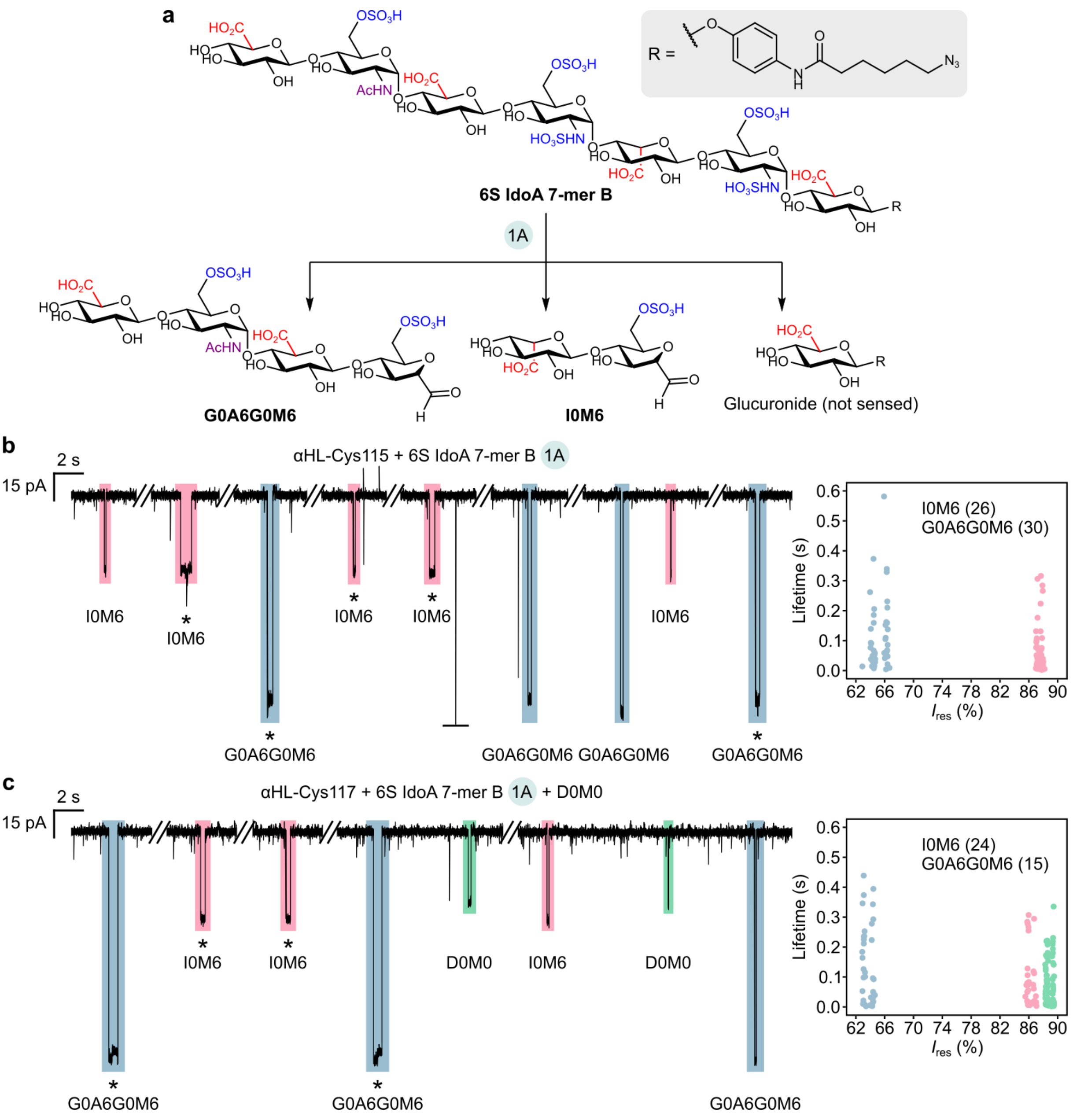

**Fig. S25 | Preparation and sensing of I0M6 and G0A6G0M6. a,** The two AMan standards were prepared from 6S IdoA 7-mer B as part of a mixture under condition **1A**. **b–c,** The mixture was directly loaded into our nanopore setup. Representative current traces for the αHL-Cys115 (**b**) and αHL-Cys117 (**c**) pores are shown, along with event assignments. In the illustrated αHL-Cys117 experiment, D0M0 was used as an internal standard. Interconversion events are marked with an asterisk. A graphically truncated event is marked with a horizontal bar. Due to low event counts, the scatter plots in **b** and **c** show combined event levels from three and two pores, respectively. Levels are colour-coded as in the traces. The number of I0M6 and G0A6G0M6 events is given in parentheses. **Recording conditions:** 4 M LiCl, 20 mM HEPBS, 40 μM EDTA, titrated to pH 8 using KOH; sugars (*cis*); +150 mV (*trans*); 23.8 ± 1 °C. I0M6, G0A6G0M6, and D0M0 concentrations were 0.15–0.22 mM, 0.15–0.22 mM, and 0.14–0.23 mM, respectively. Events with lifetimes under 2 ms were discarded (Methods).

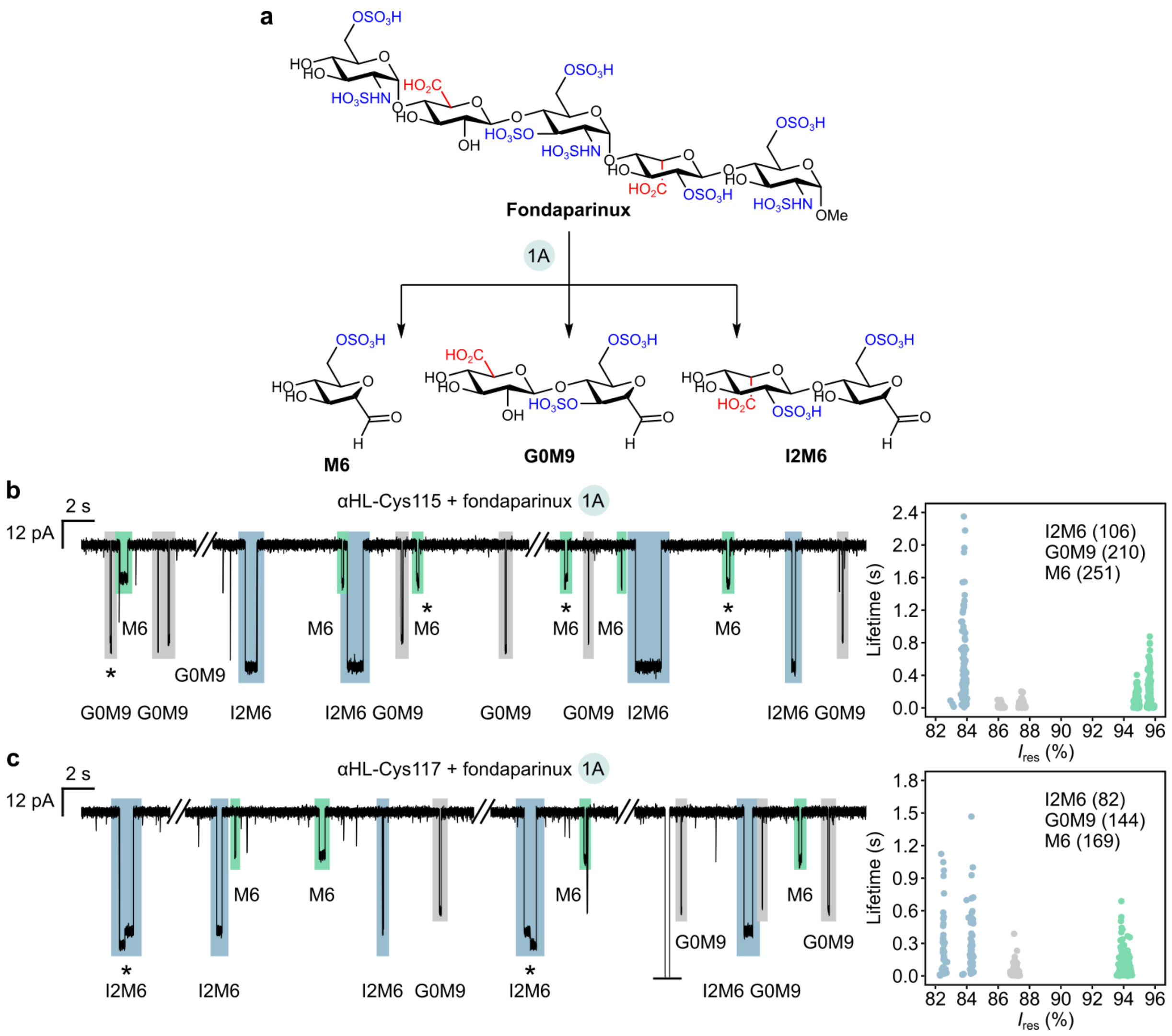

**Fig. S26 | Preparation and sensing of I2M6, G0M9, and M6. a,** The three AMan standards were prepared from fondaparinux as part of a mixture under condition **1A**. **b–c,** The mixture was directly loaded into our nanopore setup. Representative current traces for the αHL-Cys115 (**b**) and αHL-Cys117 (**c**) pores are shown, along with event assignments. Interconversion events are marked with an asterisk. A graphically truncated event is marked with a horizontal bar. The scatter plots show assigned event levels from single pores, and levels are colour-coded as in the traces. The number of sugar events is given in parentheses. **Recording conditions:** 4 M LiCl, 20 mM HEPBS, 40 μM EDTA, titrated to pH 8 using KOH; sugars (*cis*); +150 mV (*trans*); 23.8 ± 1 °C. All sugar concentrations were 0.39 mM. Events with lifetimes under 2 ms were discarded (Methods).

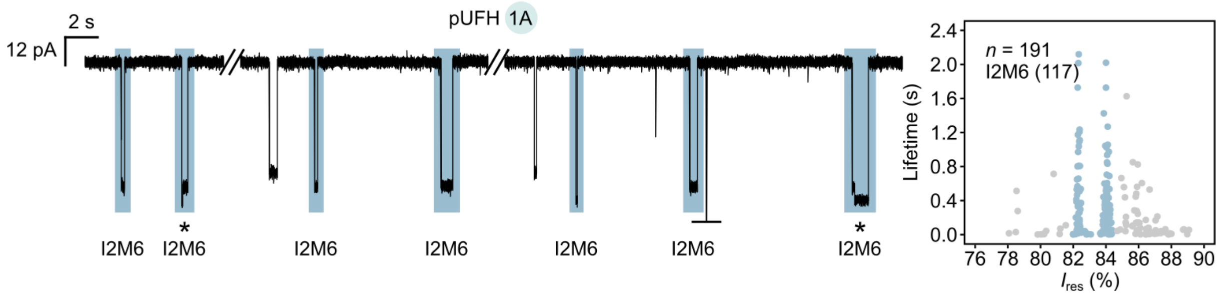

**Fig. S27 | Ring-contracted pUFH probed by the αHL-Cys117 pore.** pUFH was subjected to condition **1A** and ~0.3 mg was subsequently probed without further fractionation. A representative current trace is shown, along with assignments for I2M6 (compare to fondaparinux data in Fig. S26c). Interconversion events are marked with an asterisk. A graphically truncated event is marked with a horizontal bar. The scatter plot shows event levels from a single pore; the number of I2M6 events is given in parentheses (see below for assignment criteria). I2M6 levels are coloured in slate blue, whereas levels from all other sugars are coloured in grey. *n*, total number of events. **Recording conditions:** 4 M LiCl, 20 mM HEPBS, 40 µM EDTA, titrated to pH 8 using KOH; sugars (*cis*); +150 mV (*trans*); 23.8 ± 1 °C. Events with lifetimes under 2 ms were discarded (Methods).

### 2. Assignment of events in polysaccharide fingerprints

For each fingerprint in Figs 3–4, a basis set of structures was selected from our resolved panel of disaccharide standards according to the nature of the polysaccharide sample and/or the deconvolutive module(s) applied. For the heparins, the basis set comprised D2M6, D2M0, D0M6, and D0M0; for the heparosans, it comprised G0M0 and I0M0. To account for some small pore-to-pore variation, the $I_{res}$ values of the basis set structures (Tables S5–6) were rounded to one decimal place to give a grid of reference values against which raw, continuous event values in the fingerprints were evaluated.

In an initial round of assignment, only event $I_{res}$ values within ± 0.2 % of reference values—henceforth termed 'within range'—were assigned. This ± 0.2 % assignment window was selected in accordance with our threshold for resolving individual events with high confidence (Supplementary Note 1). Single-level events within range of only one reference were directly assigned to the corresponding sugar. Single-level events within range of multiple references were assigned to the structure corresponding to the closest reference (representing the maximum likelihood). Paired interconversion events were assigned provided all levels remained within range of the same structure, thereby enabling assignments of maximum confidence.

In a second round of assignment, unassigned events were evaluated against all other standards, and identical assignment criteria were adopted. This tiered screening strategy revealed unanticipated low-abundance sugars, many of which were unambiguously identified via interconversion events (Extended Data Fig. 8).

I2M6 in selected fingerprints (e.g., that in Fig. 5d) was assigned using a ± 0.25 % assignment window. I0T4 in the pUFH fingerprint generated under conditions **2 + 1M** (Extended Data Fig. 9d) was assigned using a ± 0.2 % window.

#### 3. Calculation of rate constants

Idealized lifetime data were imported into QuB[63,64], within which rates were extracted using the Maximum Interval Likelihood function[65,66] based on a three-state Markovian system, comprising the open-pore state and two thiohemiacetal adduct states, with six possible transitions[67,68] (Extended Data Fig. 11a). The corresponding rates are:

1. The rate of formation of adduct 1 ($v_{0,1}$)
2. The rate of formation of adduct 2 ($v_{0,2}$)
3. The rate of dissociation of adduct 1 ($v_{1,0}$)
4. The rate of dissociation of adduct 2 ($v_{2,0}$)
5. The rate of stereogenic inversion from adduct 1 to adduct 2 ($v_{1,2}$)
6. The rate of stereogenic inversion from adduct 2 to adduct 1 ($v_{2,1}$)

For each sugar, adduct 1 was chosen as the adduct that returned the higher $I_{res}$ value. Initial 'seed' values of 1 were used for all rates. Rate constants of thiohemiacetal formation were calculated from the rates using the following equations:

$$k_{0,1} = v_{0,1}/[A]$$

$$k_{0,2} = v_{0,2}/[A]$$

where [A] refers to the anhydrosugar concentration in the *cis* compartment (0.5 mL) in our experimental setup. Rate constants of thiohemiacetal dissociation and stereogenic inversion were taken as equivalent to the rates, that is:

$$k_{1,0} = v_{1,0}$$

$$k_{2,0} = v_{2,0}$$

$$k_{1,2} = v_{1,2}$$

$$k_{2,1} = v_{2,1}$$

#### 4. Ring contraction and sensing of selected polysaccharide samples

**NSOSHp:pUFH (condition 1A).** A sample of pUFH (4 mg) spiked with NSOSHp (1 mg) was subjected to condition **1A** according to the general procedure with volumes scaled correspondingly. The resulting solution was passed through an Amicon Ultra-0.5 Centrifugal Filter 3K Device (Millipore) at 14,000 × *g* and 4 °C, and 25–30 µL (corresponding to ~0.3 mg of starting material) was probed using the αHL-Cys115 pore (Fig. 5d). The remainder was stored at −80 °C until use.

**Adulterated heparin (condition 1A).** 300 µL of adulterated pharmaceutical heparin solution (lot 107031), containing no more than ~10.7 mg of active ingredient, was lyophilized and placed on ice. 500 µL of $NaNO_2$ solution was added to 500 µL of $H_2SO_4$ solution; the mixture (pH 1.5) was swirled for 3 s, then a 540 µL aliquot was immediately added to the sample. The reaction mixture was protected from direct light and allowed to warm to room temperature over 10 min, followed by the addition of 94.5 µL of 2 M $Na_2CO_3$ to quench the reaction and adjust its pH to 8. The resulting solution was filtered and probed as detailed above; in each sensing run, 30 µL (corresponding to no more than ~0.3 mg of starting material) was used.

**Adulterated heparin (condition 1M).** 110 µL of adulterated pharmaceutical heparin solution (lot 107031), containing no more than ~3.9 mg of active ingredient, was lyophilized. 500 µL of 5.5 M $NaNO_2$ solution was added to 200 µL of 0.5 M $H_2SO_4$ solution; the mixture (pH 4) was swirled for 3 s, then a 198.2 µL aliquot was immediately added to the sample. Protected from direct light, the mixture was allowed to react at room temperature for 10 min, followed by the addition of 23.1 µL of 2 M $Na_2CO_3$ to quench the reaction and adjust its pH to 8. The resulting solution was filtered and probed as detailed above; in each sensing run, 30 µL (corresponding to no more than ~0.5 mg of starting material) was used.

**Adulterated heparin (conditions 2 + 1M).** 150 µL of adulterated pharmaceutical heparin solution (lot 107031), containing no more than ~5.4 mg of active ingredient, was lyophilized. Hydrazinolysis was performed according to the general procedure (Methods) using 300 µL of hydrazine reagent. The mixture was dried under a stream of $N_2$. 500 µL of 5.5 M $NaNO_2$ solution was added to 200 µL of 0.5 M $H_2SO_4$ solution; the mixture was swirled for 3 s, then a 180 µL aliquot was immediately added to the sugars. 17.5 µL of 3 M $H_2SO_4$ was added to adjust the pH to ~4. The solution was protected from direct light and allowed to react at room temperature for 10 min, followed by the addition of 21 µL of 2 M $Na_2CO_3$ to quench the reaction and adjust its pH to 8. The resulting solution was filtered and probed as detailed above; in each sensing run, 30 µL (corresponding to no more than ~0.7 mg of starting material) was used.

**Deconvolutive analyses of standalone pUFH, NSOSHp, OSCS, and DS.** These polysaccharides were reacted according to the general procedures (Methods). To maintain consistency with the methods used to analyse the spiked and adulterated heparin samples, the ring-contraction products of pUFH, NSOSHp, OSCS, and DS were filtered before sensing as detailed above. Consistent with the lack of compounds inherent to the filter devices that may interfere with sensing, pUFH fingerprints acquired with and without such filtering were essentially identical.

### 5. NMR and MS characterization of selected disaccharides

The novel disaccharides D2M6, D2M0, D0M6, D0M0, alongside the previously reported G0M0 (ref. [69]), were reduced to their corresponding anhydromannitol (AManR) forms using $NaBH_4$ and characterized by NMR spectroscopy and MS. Removal of the AMan C1 aldehyde via reduction prevents both gradual, base-catalysed C2 epimerization[70] and potential downstream retro-aldol degradation; furthermore, eliminating *gem* diol formation[69] minimizes spectral complexity. In all cases, the NMR and MS data confirmed a single AManR product with complete preservation of fine structural features, including sulfate groups, glycosidic linkages, and UA state.

**General procedure for reduction and characterization.** The protocol for reduction was adapted from the literature[71]. 20 µL of freshly prepared $NaBH_4$ reagent (0.5 M $NaBH_4$ in 0.1 M NaOH) was added to 1 mg of anhydrosugars. The mixture was incubated at 50 °C for 15 min (500 rpm) and subsequently quenched with 10 µL of 3 M $H_2SO_4$. Following lyophilization, the products were reconstituted in $H_2O$ or $D_2O$ and directly characterized. NMR spectra were recorded on Bruker Avance III HD instruments (500 MHz). Chemical shifts were referenced to the residual protonated solvent signal. Correlation spectroscopy (COSY), heteronuclear single quantum coherence (HSQC), heteronuclear multiple bond correlation (HMBC), and nuclear Overhauser effect spectroscopy (NOESY) experiments were performed for signal assignment where necessary. NMR data are reported as follows: chemical shift, multiplicity (s = singlet, d = doublet, t = triplet, dd = doublet of doublets, m = multiplet, etc.), coupling constants (if any), integration, and assignment. High-resolution MS

(HRMS) data were acquired on a Thermo Scientific Exactive Orbitrap LC-MS instrument via electrospray ionization.

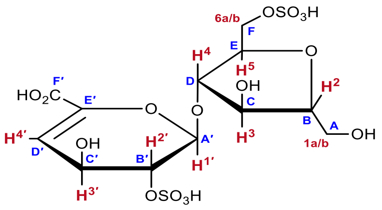

**D2m6.** Prepared from D2S6 (bn7) according to the general procedures for ring contraction at pH 1.5 (condition **1A**) and reduction. Note that AManR residues are designated as ‘m’ in the disaccharide structure code[53].

$^{1}$H NMR (500 MHz, $D_2O$, 298 K): δ 6.05 (dd, $J_{3',4'}$ = 4.7 Hz, $J_{2',4'}$ = 1.4 Hz, 1H, H-4′), 5.52 (dd, $J_{1',2'}$ = 2.4 Hz, $J_{1',3'}$ = 1.0 Hz, 1H, H-1′), 4.61 (td, $J_{1',2'}$ = $J_{2',3'}$ = 2.4 Hz, $J_{2',4'}$ = 1.4 Hz, 1H, H-2′), 4.31 (ddd, $J_{3',4'}$ = 4.7 Hz, $J_{2',3'}$ = 2.4 Hz, $J_{1',3'}$ = 1.0 Hz, 1H, H-3′), 4.27 (m, 1H, H-4), 4.24 (m, 2H, H-6a, H-6b), 4.22 (m, 1H, H-5), 4.08 (dd, $J_{2,3}$ = 7.1 Hz, $J_{3,4}$ = 5.0 Hz, 1H, H-3), 4.00 (ddd, $J_{2,3}$ = 7.1 Hz, $J_{1a,2}$ = 5.3 Hz, $J_{1b,2}$ = 3.0 Hz, 1H, H-2), 3.76 (dd, $J_{1a,1b}$ = 12.6 Hz, $J_{1b,2}$ = 3.0 Hz, 1H, H-1b), 3.68 (dd, $J_{1a,1b}$ = 12.6 Hz, $J_{1a,2}$ = 5.3 Hz, 1H, H-1a). Non-D2m6 signals: δ 8.47 ($NaHCO_2$). H-1a and H-1b assignments are consistent with the literature for IdoA2S-AManR6S[72].

$^{13}$C NMR (126 MHz, $D_2O$, 298 K): δ 169.4 (C-F′), 144.0 (C-E′), 106.7 (C-D′), 97.2 (C-A′), 86.9 (C-D), 82.9 (C-B), 79.4 (C-E), 75.1 (C-C), 74.2 (C-B′), 67.7 (C-F), 62.5 (C-C′), 60.7 (C-A). Non-D2m6 signals: δ 171.1 ($NaHCO_2$), 160.8 ($Na_2CO_3$/$NaHCO_3$). C-E′ assignment is consistent with the literature for ΔUA2S-GlcNS6S[73].

HRMS *m/z*: $[M-2H]^{2-}$ calcd for $C_{12}H_{18}O_{16}S_2$, 239.9945; found, 239.9944.

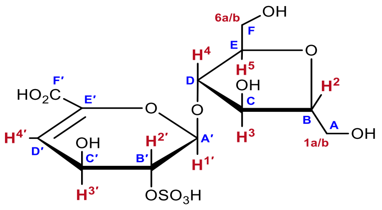

**D2m0.** Prepared from D2S0 (bn5) according to the general procedures for ring contraction at pH 1.5 (condition **1A**) and reduction.

$^{1}$H NMR (500 MHz, $D_2O$, 298 K): δ 6.03 (dd, $J_{3',4'}$ = 4.7 Hz, $J_{2',4'}$ = 1.4 Hz, 1H, H-4′), 5.48 (d, $J_{1',2'}$ = 2.4 Hz, 1H, H-1′), 4.58 (td, $J_{1',2'}$ = $J_{2',3'}$ = 2.4 Hz, $J_{2',4'}$ = 1.4 Hz, 1H, H-2′), 4.30 (ddd, $J_{3',4'}$ = 4.7 Hz, $J_{2',3'}$ = 2.4 Hz, $J_{1',3'}$ = 1.0 Hz, 1H, H-3′), 4.14 (dd, $J$ = 6.5, 5.1 Hz, 1H, H-4), 4.09–4.07 (m, 2H, H-3, H-5), 3.95 (ddd, $J_{2,3}$ = 5.5–6.0 Hz, $J_{1b,2}$ = 5.5 Hz, $J_{1a,2}$ = 3.1 Hz, 1H, H-2), 3.79 (dd, $J_{6a,6b}$ = 12.4 Hz, $J_{5,6a}$ = 3.6 Hz, 1H, H-6a), 3.77 (d, $J_{1b,2}$ = 5.5 Hz, 1H, H-1b), 3.74 (d, $J_{1a,2}$ = 3.1 Hz, 1H, H-1a), 3.67 (dd, $J_{6a,6b}$ = 12.4 Hz, $J_{5,6b}$ = 5.6 Hz, 1H, H-6b). Non-D2m0 signals: δ 8.46 ($NaHCO_2$). H-1a, H-1b, H-6a, and H-6b assignments were aided by NOESY and are consistent with the literature for IdoA-AManR[74].

$^{13}$C NMR (126 MHz, $D_2O$, 298 K): δ 144.1 (C-E′), 106.6 (C-D′), 97.1 (C-A′), 86.8 (C-D), 82.6 (C-B), 81.5 (C-E), 75.4 (C-C), 74.4 (C-B′), 62.7 (C-C′), 61.2 (C-A), 61.0 (C-F). Non-D2m0 signals: δ 171.1 ($NaHCO_2$), 161.1 ($Na_2CO_3$/$NaHCO_3$). C-F′ was not observed.

HRMS *m/z*: $[M-H]^{-}$ calcd for $C_{12}H_{18}O_{13}S$, 401.0395; found, 401.0390.

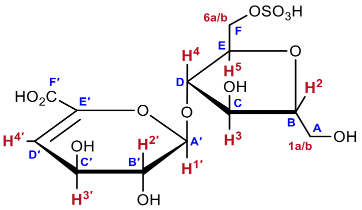

**D0m6.** Prepared from D0S6 (bn16) according to the general procedures for ring contraction at pH 1.5 (condition **1A**) and reduction.

$^{1}$H NMR (500 MHz, $D_2O$, 298 K): δ 5.91 (dd, $J_{3',4'}$ = 4.0 Hz, $J_{2',4'}$ = 0.8 Hz, 1H, H-4′), 5.18 (dd, $J_{1',2'}$ = 5.1 Hz, $J_{1',3'}$ = 0.9 Hz, 1H, H-1′), 4.25 (m, 1H, H-5), 4.24 (m, 2H, H-6a, H-6b), 4.23 (m, 1H, H-4), 4.20 (m, 1H, H-3′), 4.17 (m, 1H, H-3), 3.99 (ddd, $J_{2,3}$ = 7.6 Hz, $J_{1b,2}$ = 5.3 Hz, $J_{1a,2}$ = 3.0 Hz, 1H, H-2), 3.88 (dd, $J_{1',2'}$ = 5.1 Hz, $J_{2',3'}$ = 4.7 Hz, 1H, H-2′), 3.78 (dd, $J_{1a,1b}$ = 12.6 Hz, $J_{1a,2}$ = 3.0 Hz, 1H, H-1a), 3.70 (dd, $J_{1a,1b}$ = 12.6 Hz, $J_{1b,2}$ = 5.3 Hz, 1H, H-1b). Non-D0m6 signals: δ 8.46 ($NaHCO_2$). H-4 assignment was based on that in D2m6.

$^{13}$C NMR (126 MHz, $D_2O$, 298 K): δ 169.5 (C-F′, observed in HMBC spectrum), 144.2 (C-E′), 107.7 (C-D′), 100.5 (C-A′), 86.8 (C-D), 82.7 (C-B), 79.1 (C-E), 75.3 (C-C), 69.7 (C-B′), 67.6 (C-F), 66.2 (C-C′), 60.8 (C-A). Non-D0m6 signals: δ 171.1 ($NaHCO_2$), 161.2 ($Na_2CO_3$/$NaHCO_3$).

HRMS *m/z*: $[M-H]^-$ calcd for $C_{12}H_{18}O_{13}S$, 401.0395; found, 401.0394.

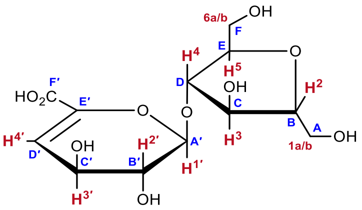

**D0m0.** Prepared from D0S0 (bn5) according to the general procedures for ring contraction at pH 1.5 (condition **1A**) and reduction.

$^{1}$H NMR (500 MHz, $D_2O$, 298 K): δ 5.90 (dd, $J_{3',4'}$ = 3.8 Hz, $J_{2',4'}$ = 0.7 Hz, 1H, H-4′), 5.15 (dd, $J_{1',2'}$ = 5.3 Hz, $J_{1',3'}$ = 0.8 Hz, 1H, H-1′), 4.21 (ddd, $J_{2',3'}$ = 4.5 Hz, $J_{3',4'}$ = 3.8 Hz, $J_{1',3'}$ = 0.8 Hz, 1H, H-3′), 4.16 (m, 1H, H-3), 4.14 (m, 1H, H-4), 4.08 (td, $J$ = 5.8, 3.5 Hz, 1H, H-5), 3.95 (ddd, $J_{2,3}$ = 7.1 Hz, $J_{1a,2}$ = 5.5 Hz, $J_{1b,2}$ = 3.1 Hz, 1H, H-2), 3.86 (ddd, $J_{1',2'}$ = 5.3 Hz, $J_{2',3'}$ = 4.5 Hz, $J_{2',4'}$ = 0.7 Hz, 1H, H-2′), 3.79 (m, 2H, H-6a, H-1b), 3.74 (dd, $J_{1a,1b}$ = 12.4 Hz, $J_{1a,2}$ = 5.5 Hz, 1H, H-1a), 3.70 (dd, $J_{6a,6b}$ = 12.5 Hz, $J_{5,6b}$ = 5.5 Hz, 1H, H-6b). Non-D0m0 signals: δ 8.46 ($NaHCO_2$). H-4 assignment was based on that in D2m0.

$^{13}$C NMR (126 MHz, $D_2O$, 298 K): δ 169.4 (C-F′), 144.2 (C-E′), 107.8 (C-D′), 100.4 (C-A′), 86.7 (C-D), 82.5 (C-B), 81.4 (C-E), 75.7 (C-C), 69.9 (C-B′), 66.4 (C-C′), 61.2 (C-A), 61.0 (C-F). Non-D0m0 signals: δ 171.1 ($NaHCO_2$), 160.9 ($Na_2CO_3$/$NaHCO_3$).

HRMS *m/z*: $[M-H]^-$ calcd for $C_{12}H_{18}O_{10}$, 321.0827; found, 321.0826.

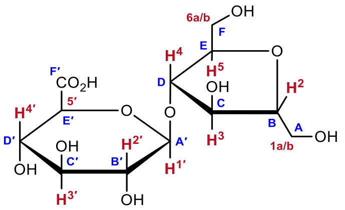

**G0m0.** Prepared from Hp according to the general procedures for conditions **2 + 1M** and reduction.

$^1$H NMR (500 MHz, $D_2O$, 298 K): δ 4.58 (d, $J_{1′,2′}$ = 8.0 Hz, 1H, H-1′), 4.26 (dd, $J_{2,3}$ = 6.7 Hz, $J_{3,4}$ = 5.2 Hz, 1H, H-3), 4.18 (dd, $J_{4,5}$ = 5.8 Hz, $J_{3,4}$ = 5.2 Hz, 1H, H-4), 4.10 (td, $J_{4,5}$ = $J_{5,6a}$ = 5.8 Hz, $J_{5,6b}$ = 3.4 Hz, 1H, H-5), 3.97 (td, $J_{1a,2}$ = $J_{2,3}$ = 6.7 Hz, $J_{1b,2}$ = 3.4 Hz, 1H, H-2), 3.94 (d, $J_{4′,5′}$ = 9.5 Hz, 1H, H-5′), 3.83–3.78 (m, $J_{1a,1b}$ = $J_{6a,6b}$ = 12.5 Hz, $J_{1b,2}$ = $J_{5,6b}$ = 3.4 Hz, 2H, H-6a, H-1b), 3.77–3.70 (m, $J_{1a,1b}$ = $J_{6a,6b}$ = 12.5 Hz, $J_{5,6a}$ = 5.8 Hz, 2H, H-1a, H-6b), 3.64–3.54 (m, 2H, H-3′, H-4′), 3.39 (t, $J_{1′,2′}$ = $J_{2′,3′}$ = 8.0 Hz, 1H, H-2′). Note pD = 4 after reconstitution in $D_2O$. Assignments are consistent with those of D2m0 and D0m0.

$^{13}$C NMR (126 MHz, $D_2O$, 298 K): δ 162.3 (C-F′, observed in HMBC spectrum), 102.1 (C-A′), 85.8 (C-D), 82.6 (C-B), 81.5 (C-E), 75.7 (C-C), 75.3 (C-C′), 74.6 (C-E′), 72.8 (C-B′), 71.4 (C-D′), 61.1 (C-A), 60.9 (C-F).

HRMS *m*/*z*: $[M+Na]^+$ calcd for $C_{12}H_{20}O_{11}$, 363.0898; found, 363.0897. No ions were observed in negative mode.

D2m6 $^{1}H$ NMR spectrum

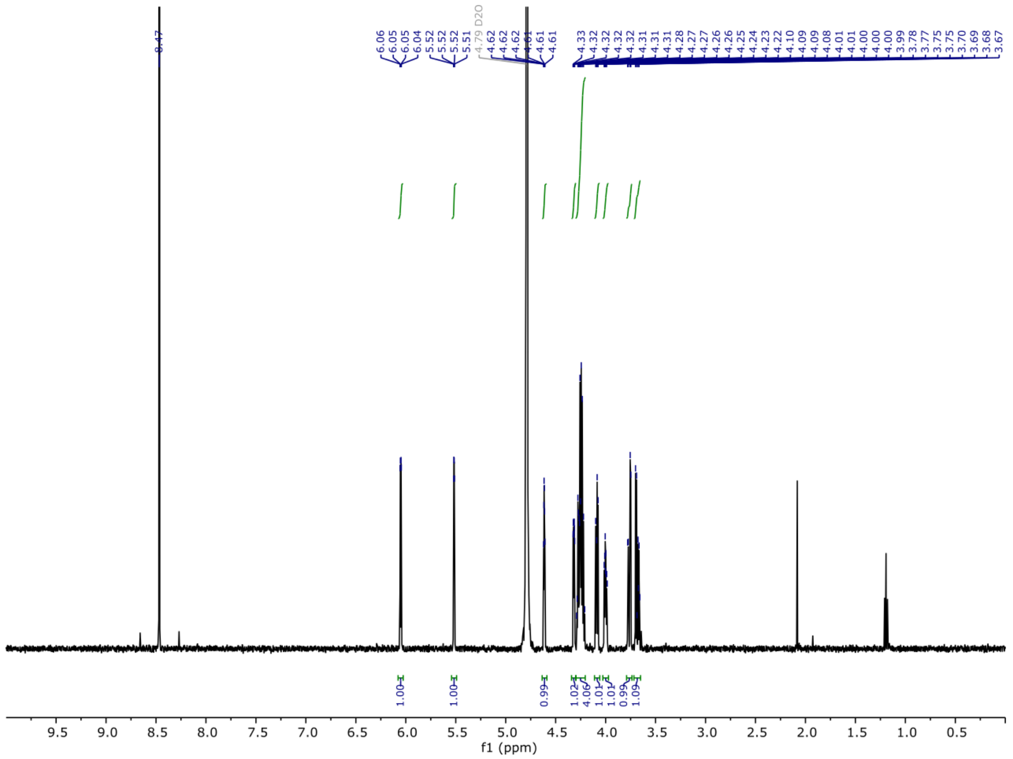

D2m6 $^{13}C$ NMR spectrum

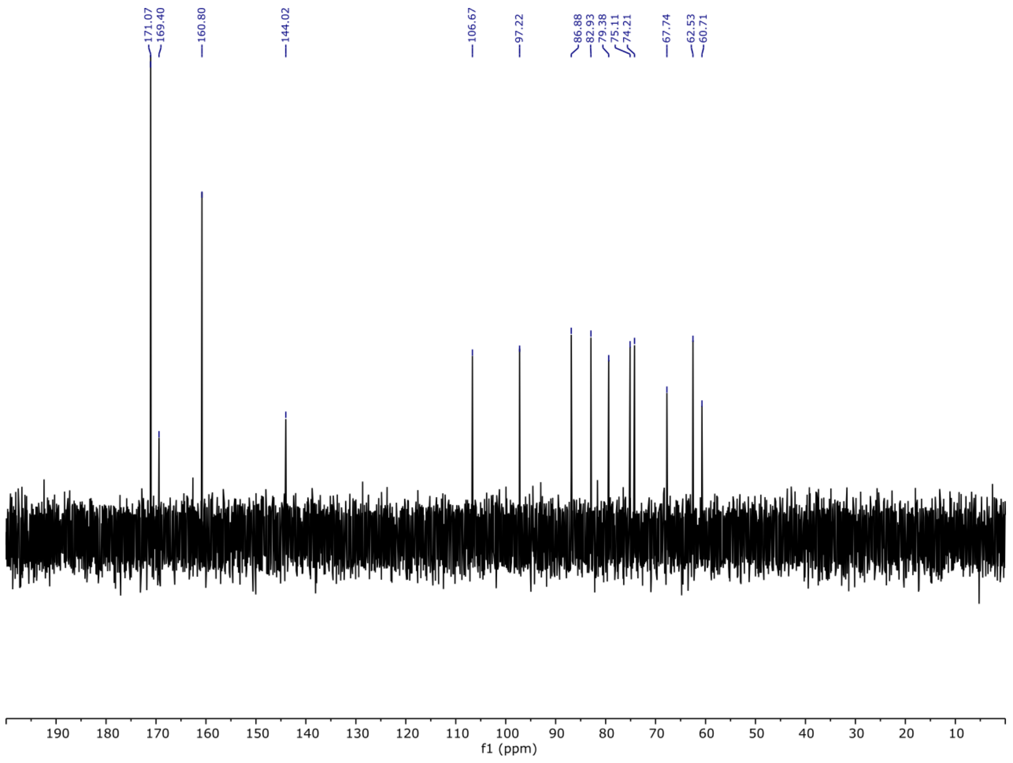

D2m0 $^{1}H$ NMR spectrum

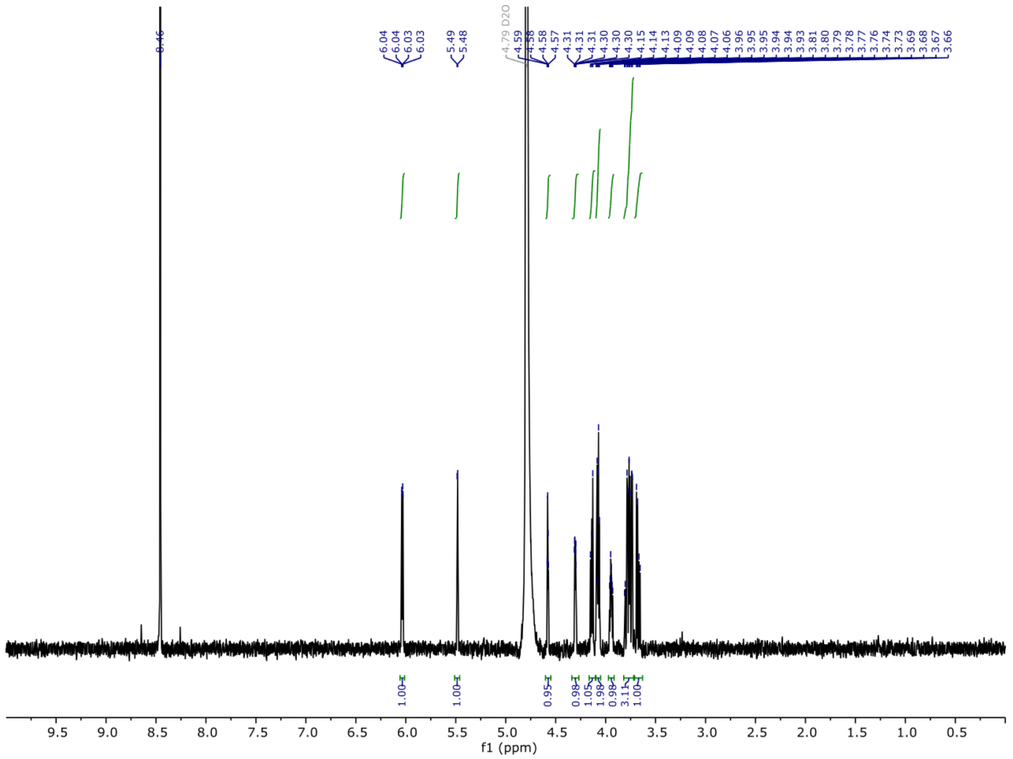

D2m0 $^{13}C$ NMR spectrum

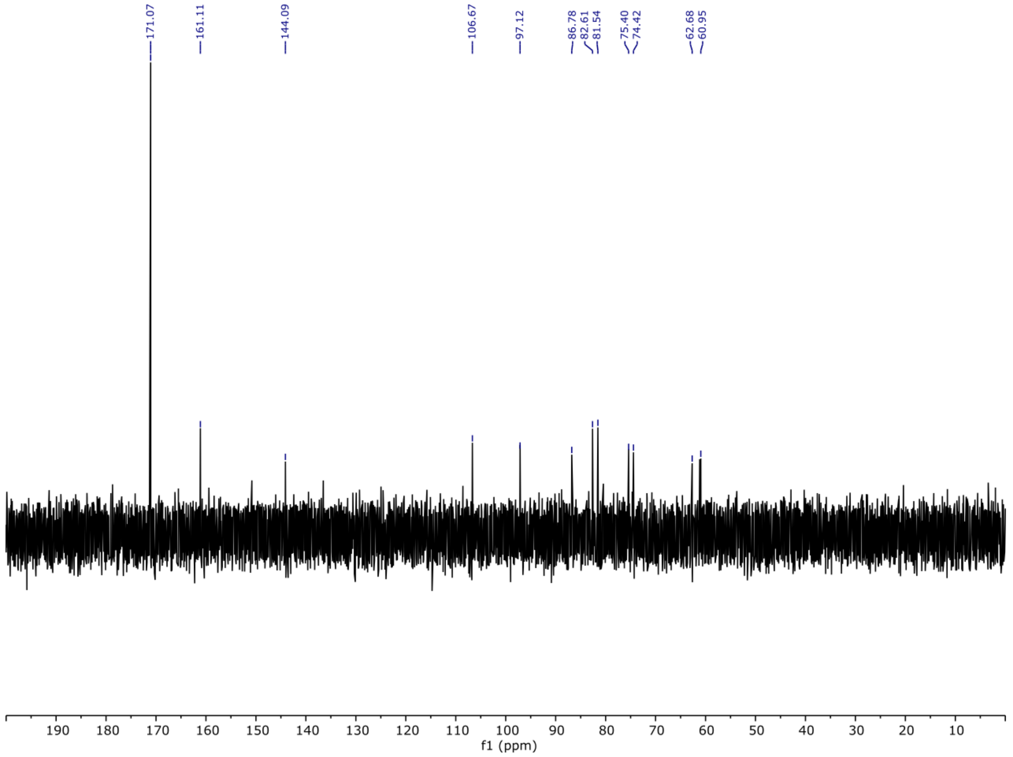

D0m6 $^{1}$H NMR spectrum

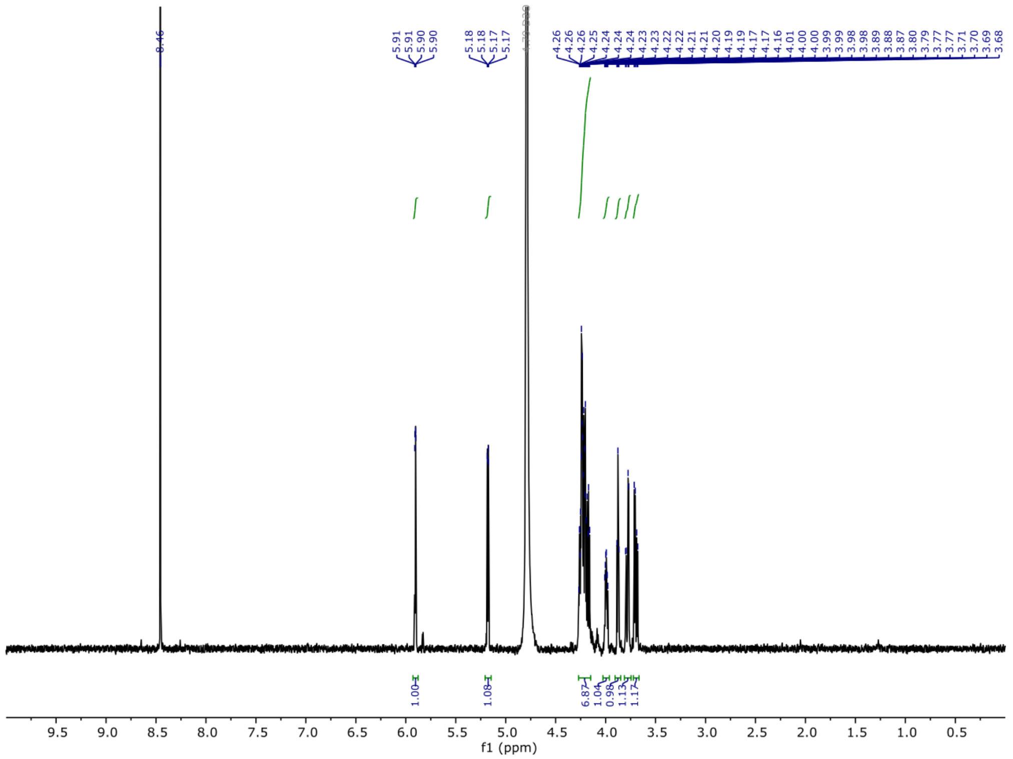

D0m6 $^{13}$C NMR spectrum

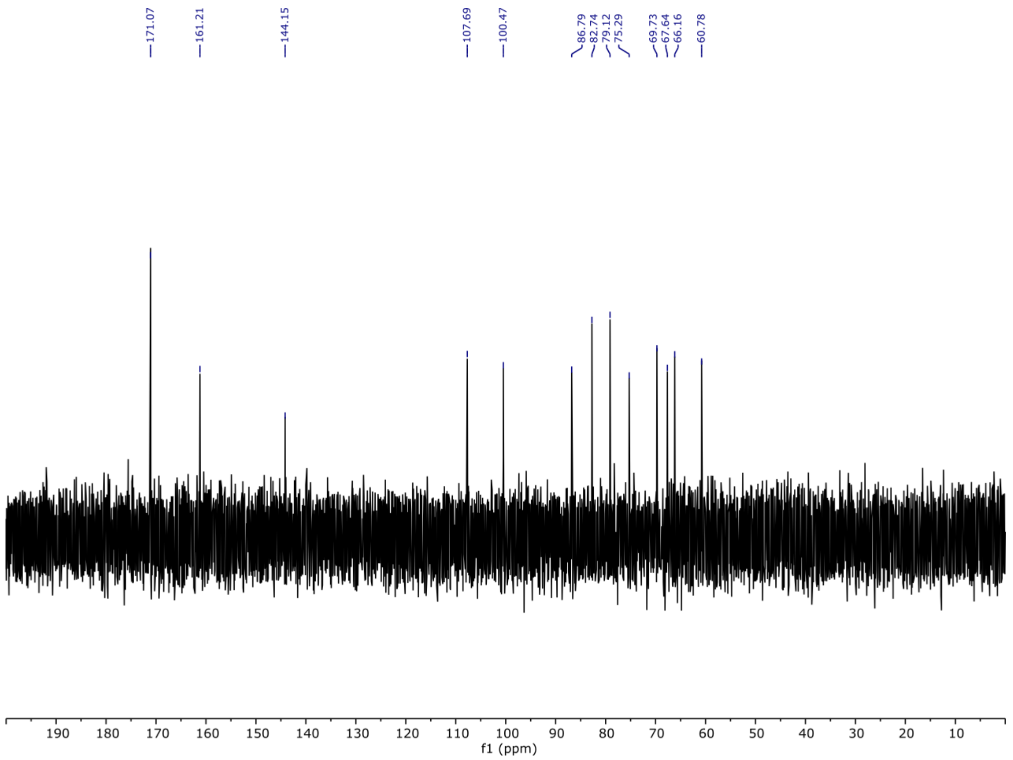

D0m0 $^{1}H$ NMR spectrum

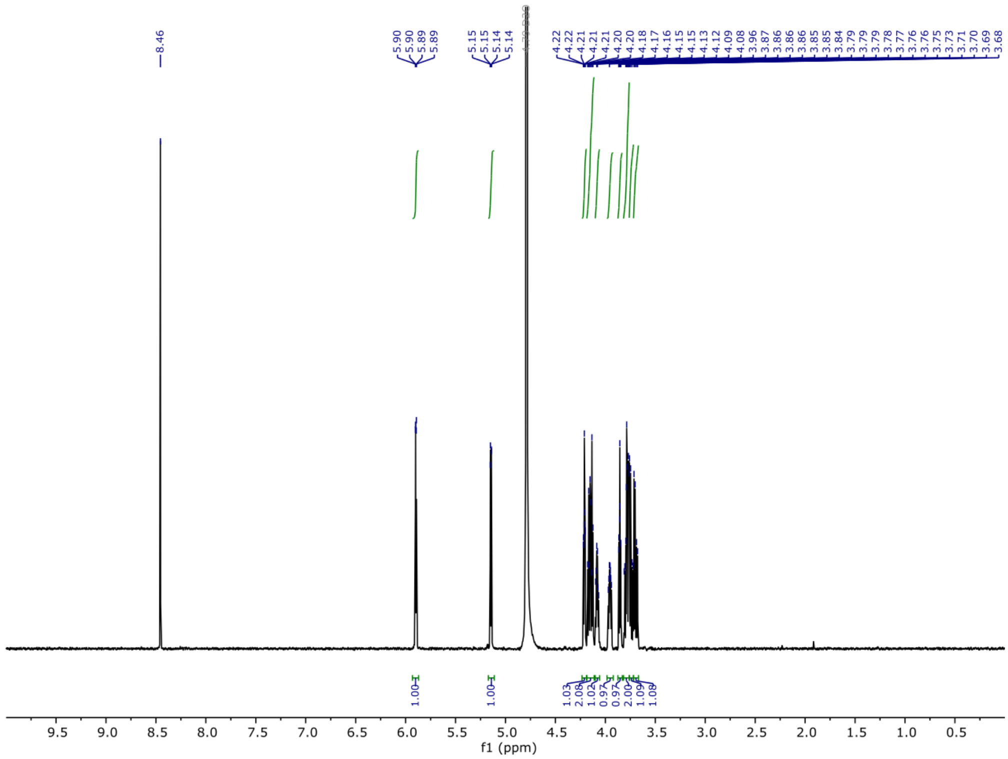

D0m0 $^{13}C$ NMR spectrum

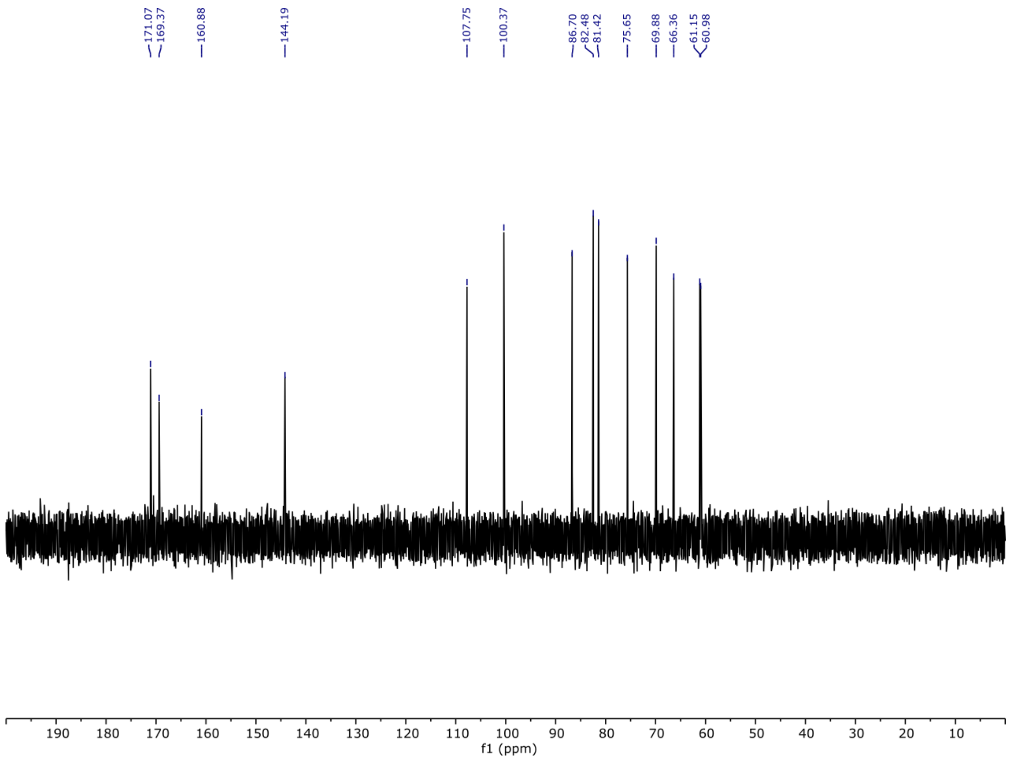

G0m0 $^1$H NMR spectrum

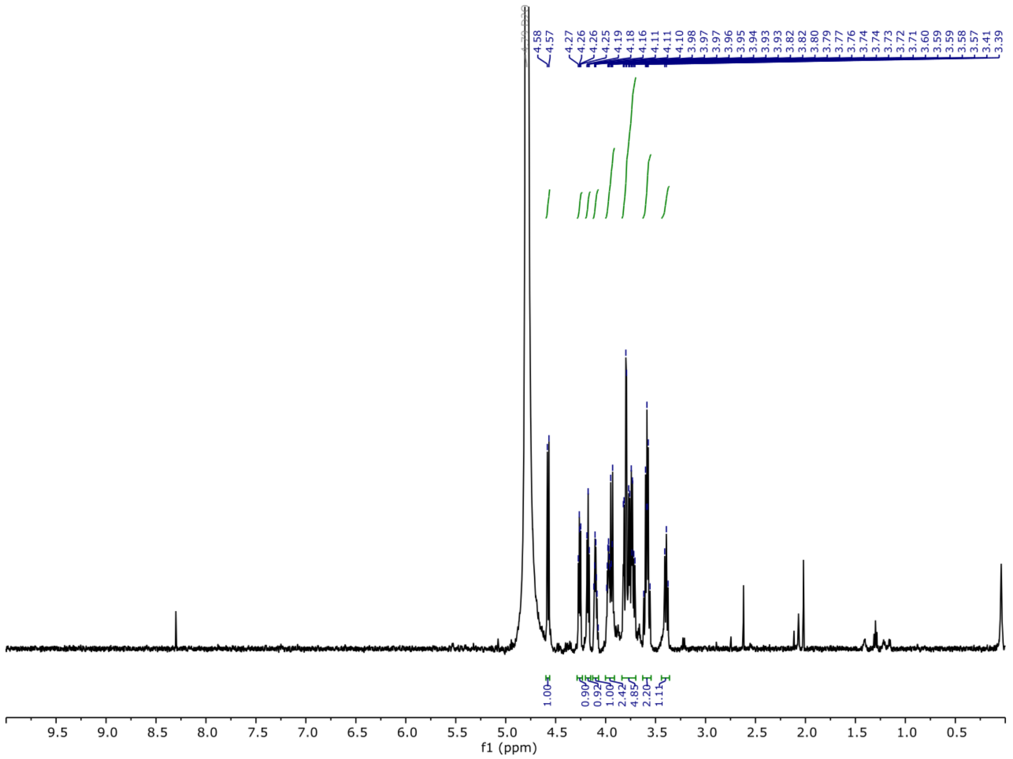

G0m0 $^{13}$C NMR spectrum

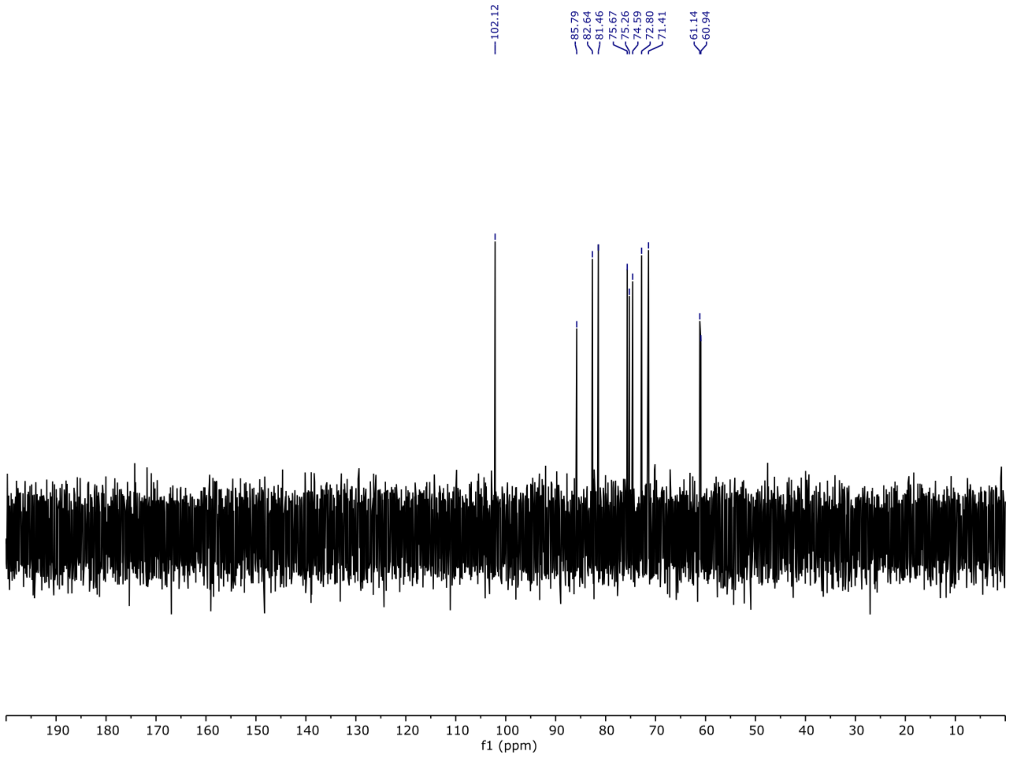
